# Microbial evolution, biogeochemical functions, and environmental adaptations in a desert saline lake on the Qinghai–Tibet Plateau

**DOI:** 10.64898/2026.07.27.740809

**Authors:** Haoran Wang, Chunhui Ai, Alexandru S. Barcan, Zhuobo Li, Binghe Zhao, Yinghui He, Yong Wang

## Abstract

The Eboliang Hu saline lakes in the hyper-arid Qaidam Basin is a high-altitude, weakly acidic hypersaline system with strong environmental gradients and limited nitrogen availability. To resolve its microbial ecology and evolutionary context, we performed genome-resolved metagenomic sequencing across four distinct habitats, reconstructing 46 medium- to high-quality metagenome-assembled genomes (MAGs) and a comprehensive gene catalog. The community shows pronounced spatial heterogeneity and is dominated by Thermodesulfobacteriota, Pseudomonadota, Bacteroidota, and archaeal lineages. Phylogenomic placement and large-scale sequence comparisons indicate that multiple dominant taxa exhibit affinity to marine- and subsurface-associated reference lineages, consistent with long-term isolation of a marine-derived ecosystem about 10–11 million years ago. Functional reconstruction reveals a distributed metabolic system in which carbon, nitrogen, and sulfur cycling are partitioned across taxa. Notably, hydrogen oxidation and arsenite oxidation are recurrent energy-producing strategies across dominant lineages, indicating redox flexibility under oligotrophic conditions. Comparative genomics further suggests lineage-specific adaptations to osmotic stress, UV exposure, and nutrient limitation. Horizontal gene transfer and phylogenetic incongruence among key metabolic genes indicate that co-evolutionary processes and gene exchange have contributed to functional innovation. These findings provide a framework for understanding microbial persistence and evolution in isolated extreme environments and offer potential analogs for extraterrestrial habitability.

## 1. Introduction

The Qaidam Basin occupies a large, intramontane depression in the northern sector of the Qinghai-Tibetan Plateau, located in north-western Qinghai Province, China. The basin floor lies at high elevation (∼2.8–2.9 km above sea level) and is surrounded by mountain ranges exceeding 5 to 6 km; its modern environment is hyper-arid, cold, and wind-dominated, with very low annual precipitation (Kong et al., 2018), high evaporation and extensive playa and salt-lake development (Stober et al., 2023). Geochemical and biomarker studies demonstrate that at least episodic marine influence into the basin occurred during the Cenozoic (notably inferred mid-Miocene seawater incursions), implying substantially lower basin elevation in parts of its history (Sun et al., 2023).

Eboliang Hu saline lake system is named after the Eboliang area located in the northwest of the Qaidam Basin. This area has an altitude of approximately 2.8 km and is more arid than the southeastern part of the basin (Liu et al., 2022). The Yadan landform makes this area known as the place in China that most resembles the surface landscape of Mars. The upper lake is fed by a hilltop spring, while the lower one lies downstream and receives water from the upper lake. These two small and shallow lakes are far away from the large salt lakes within the basin, such as Gas-Hure Hu lake and Xi Taijnar Hu lake (Stober et al., 2023), or large rivers, making it appear to be largely deprived of external organic matter and nitrogen inputs. Its area is in a state of dynamic change, which requires the microbial community in the ecosystem to cope with intermittent droughts and long-term intense ultraviolet radiation. The survival strategies of microorganisms in such extreme environments are of reference significance for the study of Earth life with the potential to colonize beyond the Earth.

We expect the Eboliang Hu lake system to host a specialized community of microorganisms uniquely adapted to its environment, also a living model for how life might endure in evaporated paleolakes on Mars or other planetary bodies. Despite the astrobiological significance, the microbial diversity, functional capacity and evolutionary characteristics of this isolated ecosystem have remained unexplored. Recent comprehensive surveys of aquatic microbial communities in other high-altitude regions of the Qinghai–Tibet Plateau, excluding the Qaidam Basin, have shown that Pseudomonadota, Bacteroidota, Actinomycetota, and Planctomycetota are often dominant in saline lake waters (Cheng et al., 2024), whereas Thermodesulfobacteriota genomes are more readily recovered from sediments (Zhang et al., 2025). In contrast, a metagenomic survey of regolith soil samples from the western region of the Eboliang area reported that Pseudomonadota and Actinomycetota were the most abundant phyla (Liu et al., 2022; Liu et al., 2025). Therefore, the specific community structure and genomic diversity in the Eboliang Hu lakes would be a great complement, and it would be interesting to compare it with previously studied nearby microbial communities and even those in other “Martian bases” on Earth. Moreover, given the basin’s history, present-day community may include relic marine-associated lineages that have persisted after long-term isolation, rather than being assembled exclusively from local terrestrial taxa. Such signals have not been examined or analyzed in other lakes on the Qinghai–Tibet Plateau.

In this study, we performed a genome-resolved metagenomic analysis of sediments-associated microbial communities from Eboliang Hu saline lake system. Microbial community composition across four environmentally distinct sampling sites was reconstructed and metagenome-assembled genomes (MAGs) spanning wide bacterial and archaeal lineages were recovered. We further resolved functional gene distributions at both community and genome levels to infer core metabolic capacities, with emphasis on carbon, nitrogen, and sulfur cycling under extreme salinity and nutrient limitation. In addition, we examined phylogenetic affiliations of dominant taxa and MAGs to evaluate their evolutionary relationships with reference genomes from diverse environment, including potential marine-associated lineages. Finally, we assessed genome-level horizontal gene transfer (HGT) events to explore mechanisms that may contribute to functional innovation and environmental adaptation in this isolated ecosystem. By combining genome-resolved metagenomics with large-scale sequence comparison and phylogenetic placement of both marker genes and functional proteins, this study not only expands the known genomic diversity of high-altitude saline lake microbiomes, but also provides a high-resolution framework for assessing evolutionary origins and functional convergence in extreme terrestrial ecosystems.

## 2. Materials and methods

### 2.1 Sampling, determination of liquid-phase chemical parameters

Fig. 1 illustrates the spatial layout of the Eboliang saline lake system (92.8916 E, 38.3595 N) and the four sampling sites surveyed in July 2022. Fig. 1A shows the position of the lake within the Eboliang area of the Qaidam Basin, elevation data from GEBCO 2024 NetCDF. Fig. 1B presents representative field photographs of the sampling sites (the location of the lake is marked on the satellite image map, which is sourced from the Chinese National Platform for Common GeoSpatial Information Services “t0.tianditu.com/img_c/esri/wmts”).

**Figure 1.**
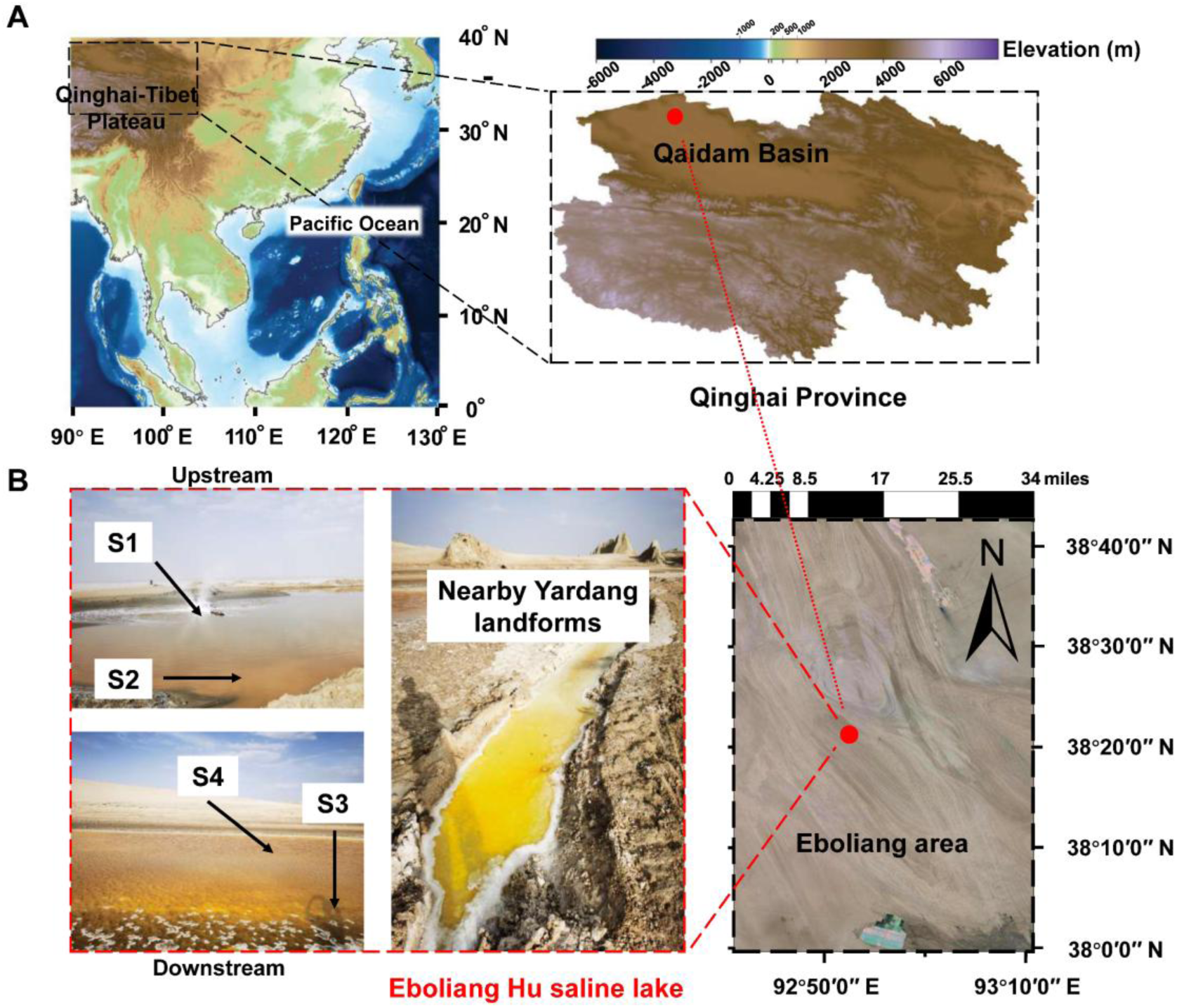
The spatial layout of the Eboliang Hu saline lake system and the four sampling sites. (A) Position of the lakes within the Eboliang area of the Qaidam Basin (Qinghai Province, China). (B) Representative field photographs of the sampling sites.

Samples were kept frozen and transported to the laboratory for subsequent chemical parameter analysis of liquid phase composition. The total salinity is measured by a salinity sensor (SWEVY, Guangzhou, China). The pH value was measured with the FiveEasy Plus pH meter (METTLER TOLEDO, Greifensee, Switzerland). For the analysis of inorganic salts, liquid samples were first filtered through a 0.22-µm membrane (Wang et al., 2024). Concentrations of nitrate, nitrite, ammonium, sulfate, and sulfide were subsequently quantified using Hach reagents (NO_3_^-^-N: 2429800; NO_2_^-^-N: 2107169; NH_4_^+^-N: 2604545; SO_4_²⁻: 2106769; S²⁻: 2244500) in combination with a DR900 colorimeter (Hach, Loveland, USA).

### 2.2. DNA extraction, metagenomic library construction and Illumina sequencing

For each sample, 0.25 g of wet sediment was used for DNA extraction, following the operating instructions of the DNeasy PowerMax Soil Kit (Qiagen, Hilden, Germany). The Covaris M220 Focused ultrasonicator (Covaris, Massachusetts, USA) was used to fragment 100 ng of DNA to ∼350 bp. The high-throughput metagenome libraries were then created using the VAHTS Universal DNA library prep kit for Illumina V3 (Vazyme, Nanjing, China) in accordance with the manufacturer’s instructions. A NovaSeq 6000 System (Illumina, San Diego, USA) was used to sequence the libraries using PE 2 × 150 bp.

### 2.3 Taxonomic classification based on 16S and 18S rRNA gene

Raw reads were processed to obtain clean reads for rRNA gene extraction and metagenome assembly, as previously described (He et al., 2025) (Supplementary methods and results 2).

The 16S and 18S rRNA fragments longer than 100 bp were identified from clean reads by utilizing rRNA_HMM (Huang et al., 2009). As previously described (Zhang et al., 2024), HMMsearch was used to further identify the fragments that mapped to V4 regions of 16S rRNA genes and V9 regions of 18S rRNA.

For species annotation of prokaryotic microbial communities, the V4 region sequence was imported into QIIME2 (v.2024.2.0) (Bolyen et al., 2019), and the sequence was deduplicated using VSEARCH and clustered into operational taxonomic units (OTUs) based on a similarity level of ≥ 97%. The feature classifier “classify sklearn” command in QIIME2 refers to the SILIVA 138 database (Quast et al., 2013) to annotate representative sequences of OTUs. Alpha diversity metrics were computed using the QIIME2 core-metrics-phylogenetic pipeline: a rooted phylogenetic tree of representative sequences was supplied to enable calculation of Faith’s phylogenetic diversity; Shannon entropy and Pielou’s evenness were also generated. BLAST (v.2.16.0+) was used to search the 16S gene sequences against the SILVA SSU rRNA database (release 138), and a threshold of 97% was used to find novel 16S gene sequences. The number of novel 16S gene sequences found in each sample was divided by the total number of 16S gene sequences to determine the novel 16S sequence rates (Zhou et al., 2022). Bray–Curtis dissimilarity matrices were calculated based on order-level relative abundances, and weighted-unifrac distance matrices, incorporating both representative OTU abundances and their phylogenetic relationships, were constructed. These distance matrices were subsequently used for principal coordinates analysis (PCoA).

For species annotation of eukaryotic microorganisms, a similar process was followed. The V9 region sequence was imported into QIIME2, and the annotation of representative sequences was referenced in the PR2 (v.5.0.0) database (Guillou et al., 2013). To more clearly illustrate the phylogenetic positions of the dominant OTU sequences, a phylogenetic tree was constructed using these sequences together with rRNA sequences of the closest matching, taxonomically identified species obtained from the top BLASTn hits. Specifically, multiple sequence alignment of the combined 18S reference sequences and dominant V9 OTU sequences was performed using MAFFT (v.7.505) (Katoh and Standley, 2013) with the L-INS-i strategy, enabling local-pair refinement and up to 1,000 iterative improvement cycles. Poorly aligned or ambiguously aligned positions were trimmed using trimAl (v.1.5) (Capella-Gutiérrez et al., 2009) with the “automated1” heuristic. Maximum-likelihood phylogenetic inference was conducted with IQ-TREE2 (v.2.2.0) (Minh et al., 2020), employing ModelFinder Plus (MFP) for optimal substitution model selection. Branch support was assessed using 1,000 ultrafast bootstrap replicates (-B 1000) and 1,000 SH-aLRT tests (-alrt 1000), with additional NNI optimization of bootstrap trees (--bnni). iTOL v.7 (https://itol.embl.de/) was used to visualize the phylogenetic tree.

Newick Utilities (v.1.6; nw_prune) (Junier and Zdobnov, 2010) was used to prune large phylogenetic trees into simplified representations while retaining their topological structure.

### 2.4 Metagenome assembly and genome binning

Clean reads of each sample were assembled using SPAdes (v.3.15.5) software (Bankevich et al., 2012) with the parameters (-min-contig-len 1300; -k 21, 33, 55, 77, 99, 127). After filtering out eukaryotic contigs using EukRep (v.0.6.7) (West et al., 2018), the prokaryotic contigs were binned using the metaWRAP (v.1.3.2) pipeline (Uritskiy et al., 2018). Initial bins were generated with three independent binning algorithms—MetaBAT2, MaxBin2, and CONCOCT—using contigs ≥ 2 kb and differential coverage profiles derived from paired-end clean reads. Consensus bin refinement was then conducted with metaWRAP’s bin_refinement module, which integrates the three binning outputs and retains bins with ≥ 50% completeness and ≤ 10% contamination as assessed by CheckM (Parks et al., 2015). To further improve genome quality, each refined bin was subjected to read extraction and individual reassembly using the metaWRAP reassemble_bins module, yielding reassembled bins with enhanced contiguity and completeness. MAGs were dereplicated using dRep (v.3.5.0) (parameters: -sa 0.97 -nc 0.30 -cm larger -comp 50 -con 10) to remove redundant genomes and retain a non-redundant representative (with the largest assembly size) set (Olm et al., 2017). To assess genome chimerism and lineage-specific contamination, the dereplicated MAGs were evaluated using GUNC (v. 1.0.6) (Orakov et al., 2021).

### 2.5 Genome abundance and phylogenetic analysis

CoverM (v.0.7.0) (Aroney et al., 2025) was used to estimate the relative abundance of MAGs based on the recruitment rates of clean reads (parameters: --min-read-aligned-length 50 --min-read-percent-identity 0.95 --min-covered-fraction 0.1).

Taxonomic and phylogenetic placement of the dereplicated, GUNC-passed MAGs was performed using GTDB-Tk (v.2.3.2) with the GTDB Release 207_v2 reference (Chaumeil et al., 2020; Parks et al., 2022), yielding standardized GTDB taxonomic assignments. Following classification, phylogenetically proximate reference genomes were obtained from NCBI (RefSeq genomes given priority) and combined with the representative MAGs to generate a genome-resolved phylogenetic tree. Phylogenomic inference was conducted by identifying single-copy marker genes using CheckM’s “lineage_wf” (v.1.2.2), aligning sequences with MAFFT, trimming poorly aligned regions using trimAl, and reconstructing a maximum-likelihood tree with IQ-TREE2.

### 2.6 Functional gene annotation and relative abundance analysis

Prodigal (v.2.6.3) (Hyatt et al., 2010) with option “-p meta” was used to predict open reading frames (ORFs) and translated protein sequences for individual genomes. KofamScan (v.1.1.0) (Aramaki et al., 2020) was used to assign KEGG Orthologs (KOs) to these protein sequences. Clusters of orthologous groups (COGs) were annotated by BLASTp searches against the COG protein-sequence database (v.9.05) with an e-value cutoff of 1e-5. Carbohydrate-active enzymes (CAZymes) were annotated using dbCAN (v.4.1.4) (Zheng et al., 2023), which identifies CAZy families from predicted protein sequences based on the CAZy HMM database.

For clarity and consistency, gene names rather than protein names were consistently used in describing annotation results and in verifying the species origin of gene-encoded protein sequences based on BLASTp searches.

To quantify the distribution of functional genes across phyla, the frequency of each marker gene (with KO) was calculated as the proportion of MAGs within each phylum that encoded the gene (Li et al., 2025b).

Similarly, after the clean reads were assembled using MEGAHIT (v.1.2.9) (Li et al., 2016) ORFs were predicted from the resulting contigs using Prodigal. All predicted coding sequences were clustered with MMseqs2 (v.15.6f452) (Steinegger and Söding, 2017) at 95% nucleotide identity and 80% coverage (Cheng et al., 2024; Chibani et al., 2022; Yu et al., 2024) to generate a non-redundant gene catalog, from which representative sequences were extracted. This catalog served as the reference for abundance estimation. Paired-end reads were mapped to the catalog using CoverM (“tpm” mode) with stringent alignment thresholds (≥ 95% identity, ≥ 75% read coverage, and ≥ 50% gene coverage), and TPM values were calculated for each gene across samples. The final abundance of each functional gene (KO) was derived by summing the TPMs of all genes annotated to that KO.

For gene *aoxB* (K08356, arsenite oxidase large subunit AioA), contig-derived sequences annotated as *aoxB* by KofamScan were combined with reference sequences to reconstruct a maximum-likelihood phylogeny, in order to infer the potential taxonomic origin of sequences that were not assigned to MAGs. The reference set consisted of arsenite oxidase ARO sequences from the Greening Lab database of metabolic marker proteins (Greening et al., 2019), which were dereplicated using CD-HIT (v.4.8.1; parameters: -c 0.90, -n 5) (Fu et al., 2012) and further filtered to remove sequences that had been suppressed in NCBI (no longer annotated on any genome). This yielded 214 non-redundant reference sequences, all corresponding to RefSeq non-redundant protein accessions (WP_). AlphaFold 3 model (Abramson et al., 2024) was used to predict the three-dimensional structure of AioA subunit.

### 2.7 Community-level horizontal gene transfer identification

MetaCHIP (v.1.10.13) (Song et al., 2019) was used to infer HGT events from dereplicated MAGs, and genes involved in these events were functionally classified and annotated using KEGG.

For genes detected to have undergone horizontal gene transfer events, further phylogenetic analysis of their protein sequences was performed using RAxML (v.8.2.12) under the PROTGAMMALG model (Stamatakis, 2014). Maximum-likelihood trees were inferred with 1,000 rapid bootstrap replicates using the -f a option, and bootstrap support values were mapped onto the best-scoring tree.

## 3. Results

### 3.1 Source environment and physicochemical properties of the samples

The four sampling sites span distinct geomorphological and hydrological settings: S1 is a shallow site near the spring at the upstream margin of the upper lake; S2 is a deep-water site on the opposite side of the upper lake; S3 is a nearshore site in the downstream lake; and S4 is a deep-water site in the downstream lake (Fig. 1B). The water in the downstream lake was yellow and exhibited extensive salt precipitation resulting from intense evaporation. Together, these sites represent the major environmental niches along the Eboliang Hu saline lake transect.

Sediment and overlying water samples from the four sites showed a clear salinity gradient from S1 (1.97%) to S4 (6.84%) (Fig. 2A). All aqueous phases were weakly acidic (pH 6.37–6.60), indicating that the Eboliang Hu saline lakes deviate from the saline-alkaline conditions typical of most lakes in the Qaidam Basin (Stober et al., 2023; Wang et al., 2014). Ammonium-N concentrations increased with salinity and ranged from 25.2 to 45.6 mg L^-1^. Sulfate and sulfide concentrations varied markedly among sites, with the highest values observed at S3 (3,225 mg L^-1^ sulfate and 9.75 mg L^-1^ sulfide) and the lowest at S2 (1,050 mg L^-1^ sulfate and 3.75 mg L^-1^ sulfide). Nitrate- and nitrite-nitrogen concentrations were below 0.06 mg/L, as determined by the diazonium salt-based colorimetric assay (Supplementary methods and results 1).

**Figure 2.**
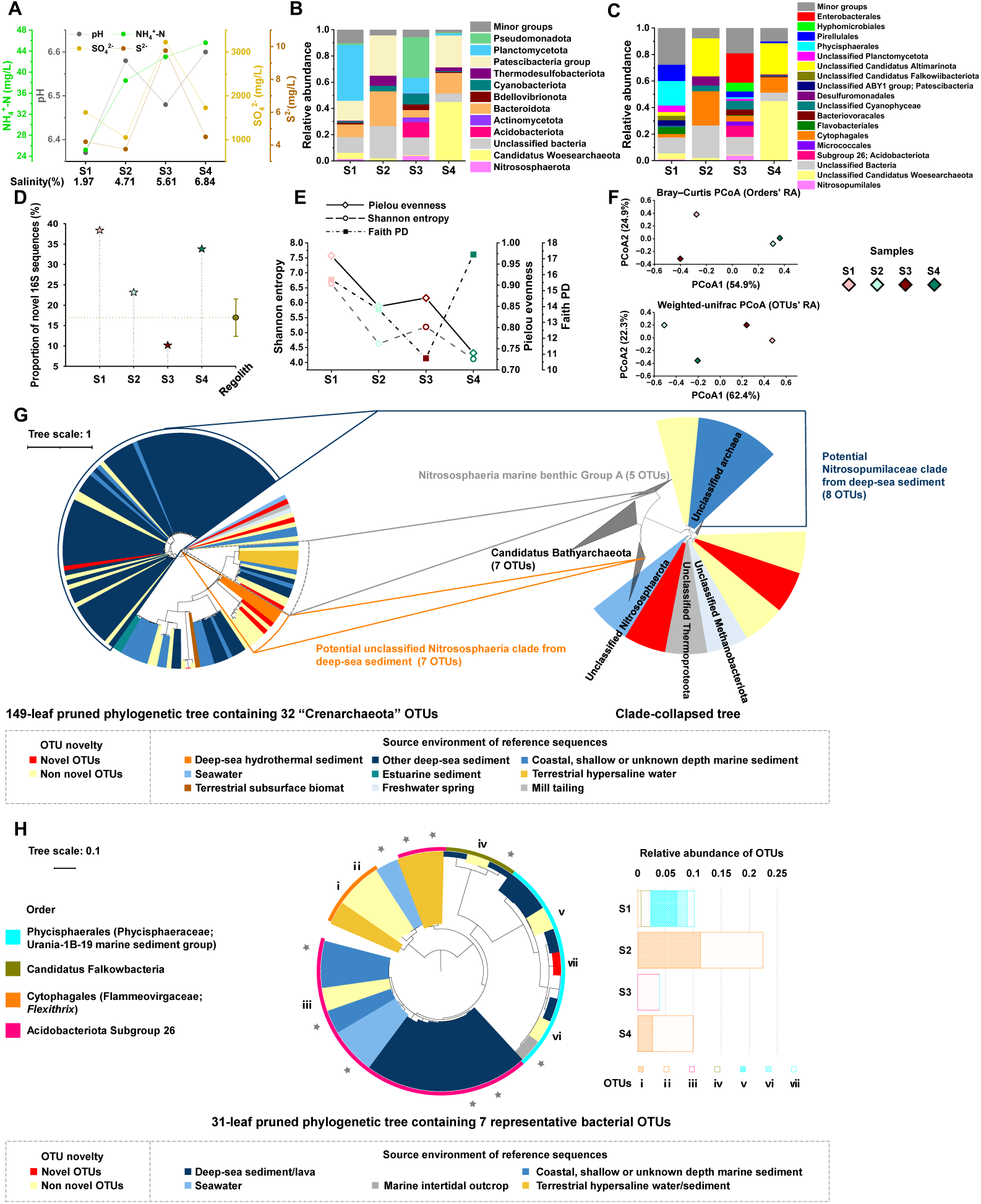
Ion chemistry of the water phase, prokaryotic community structure in sediments (based on 16S rRNA gene V4-region sequences), and phylogenetic analysis of representative sequences of OTUs. (A) Concentrations of key aqueous-phase parameters: salinity, pH, ammonium-N, sulfate and sulfide. (B) Relative abundances of dominant phyla (> 3% relative abundance in at least one sample) across samples (“Minor groups” denotes the combined category of OTUs other than the dominant phyla and bacterial OTUs not classified at the phylum level). (C) Relative abundances at the order level (Only dominant orders with relative abundance > 3% in any sample are shown. “Minor groups” denotes the combined category of OTUs other than the dominant orders and bacterial OTUs not classified at the phylum level). (D) Novelty of 16S rRNA sequences (the proportion of novel 16S sequences); “Regolith” sample data (NCBI BioSample ID “SAMN21365075”–“SAMN21365078”) of the Qaidam Basin are from Liu et al. (2022). (E) α-diversity metrics for each sample (Shannon entropy, Pielou’s evenness, and Faith’s phylogenetic diversity). (F) Principal coordinates analysis (PCoA) summarizing between-sample community similarity (“RA”: Relative abundance). (G) Maximum-likelihood phylogeny of representative OTUs assigned to the phylum “Crenarchaeota” and non-redundant reference V4 sequences from the SILVA 138 database; OTU novelty and the environmental origins of reference sequences are indicated by distinct colors (legend indicates color–environment mapping); the labels of the 5 reference sequences outside the clusters are based on NCBI taxonomy classification. (H) Maximum-likelihood phylogeny of representative bacterial OTUs and non-redundant reference V4 sequences from the SILVA 138 database and the NCBI core nucleotide database; the relative abundances of these OTUs are shown in the stacked bar plot; gray stars indicate reference sequences not included in the SILVA Ref database.

### 3.2 Microbial community composition

The prokaryotic community at site S1 was characterized by a pronounced dominance of Planctomycetota, which accounted for 42.5% of the total community (Fig. 2B). Members of the Patescibacteria (classified as the phylum level in the SILVA database) group, including multiple *Candidatus* lineages, represented 15.2%, while Bacteroidota comprised 9.7% of the community. The sediment communities at S2 and S4, representing the upstream and downstream deep-water sites, respectively, exhibited broadly similar phylum-level profiles, characterized by relatively high abundances of Patescibacteria and Bacteroidota and low proportions of Planctomycetota (< 1.5%). At S2, Thermodesulfobacteriota reached a relative abundance of 7.7%, the highest observed among all samples, whereas the archaeal lineage *Candidatus* Woesearchaeota was most abundant at S4, accounting for 45.0% of the community. In contrast, S3 displayed a distinct community structure, dominated by Pseudomonadota (31.0%) and Acidobacteriota (11.6%). Additional taxa with relative abundances exceeding 3% at this site included Planctomycetota (12.1%), Cyanobacteriota (8.0%), Bacteroidota (5.7%), Bdellovibrionota (4.2%), Actinomycetota (3.6%), and the archaeal phylum Nitrososphaerota (3.8%).

At a finer taxonomic resolution, Planctomycetota at site S1 were primarily affiliated with the orders Phycisphaerales and Pirellulales, accounting for 18.5% and 12.2% of the order-level community, respectively. The dominant Patescibacteria at this site were mainly assigned to the unclassified ABY1 group, *Candidatus* Falkowiibacteriota, and *Candidatus* Altimarinota, whereas Bacteroidota were predominantly represented by Flavobacteriales and Cytophagales. At sites S2 and S4, Patescibacteria were almost exclusively affiliated with *Candidatus* Altimarinota, and Bacteroidota were largely represented by Cytophagales. At S2, Desulfuromonadales accounted for 7.0% of the community. At S3, the dominant Proteobacteria were affiliated with Hyphomicrobiales and Enterobacterales. Similar to S2, Cyanobacteriota at S3 were mainly represented by taxa within the class Cyanophyceae that could not be confidently resolved at the order level. The relative abundances of the dominant phyla for S3 shown in Fig. 2B were mainly contributed by the orders Subgroup 26 (8.4%), Cytophagales (4.4%), Bacteriovoracales (4.2%), Micrococcales (3.2%), and Nitrosopumilales (3.8%), as illustrated in Fig. 2C.

At site S2, 24.5% of V4-region sequences could not be assigned to any known phylum, a higher proportion than in the other samples (Fig. 2B). However, BLASTn searches of all 16S rRNA sequences revealed that S1 and S4 harbored higher proportions of novel sequences, both exceeding 30% (Fig. 2D). For comparison, regolith surface metagenomes collected previously (Liu et al., 2022) from four sites in the basin (elevation: 2,797–3,151 m, see Supplementary Table S1) showed novelty level of 16.94 ± 4.59%.

Across the four samples, the relative patterns of Shannon entropy and Pielou’s evenness were consistent, with the highest values observed at S1 and the lowest at S4. In contrast, when phylogenetic relationships among representative OTUs were taken into account, another α-diversity metric—Faith’s phylogenetic diversity—was highest at S4 (Fig. 2E). PCoA analysis based on Bray–Curtis dissimilarities calculated from order-level relative abundances showed that S2 and S4 clustered closely together, consistent with their similar taxonomic composition at this level. In contrast, when phylogenetic relationships among OTUs were taken into account using weighted-unifrac distances, S2 and S4 no longer clustered together. Instead, the distance between S2 and S4 exceeded that observed between S1 and S3, indicating pronounced differences in phylogenetic community structure despite their taxonomic similarity (Fig. 2F).

Clustering of 18S rRNA V9-region sequences yielded 23 representative OTUs, among which nine dominant OTUs (relative abundance > 10%, whereas all remaining OTUs accounted for < 3% each) constituted the major components of the eukaryotic microbial communities. At site S1, the community was dominated by an unclassified lineage within the subclass Labyrinthulomycetes (40%) (OTU4: Labyrinthulomycetes X group), followed by Tulamoebidae (OTU5), Trypanosomatidae (OTU8), and an unclassified Choanoflagellatea lineage (OTU9: Choanoflagellatea XX group), each contributing 10%. At S2, only unclassified eukaryotic sequences were detected. At S4, one-third of the sequences were assigned to the unclassified Choanoflagellatea lineage, whereas fungal sequences accounted for 92.9% of the community at S3. Across sites S1, S2, and S4, unclassified eukaryotic sequences were primarily represented by OTU3, OTU6, and OTU7, although their relative distributions differed among samples. Phylogenetic relationships between the dominant OTUs and their closest reference rRNA gene sequences with species-level annotations are also shown in Fig. S2. Analyses based solely on the V9 region are subject to inherent limitations. When all 18S rRNA gene sequences were taxonomically annotated, additional eukaryotic lineages beyond the three dominant groups—Discoba, Stramenopiles, and Opisthokonta—were detected, including members of the classes Spirotrichea and Flabellinia, suggesting that these taxa may also inhabit this ecosystem (Fig. S2).

### 3.3 Marine affinity of species in communities

Phylogenetic analyses were performed to infer the environmental origins of taxa most closely related to the detected 16S rRNA gene sequences (see Supplementary Table S2). For representative V4 OTUs assigned to the phyla “Crenarchaeota” and “Nitrospirota”, phylogenetic trees were constructed together with non-redundant reference V4 sequences from the SILVA 138 database (7,190 and 1,204 sequences, respectively).

Among the Nitrospirota sequences that clustered with reference lineages and exceeded 70 bp in length, two OTUs were identified. One grouped with 3 terrestrial reference sequences originating from arid regions in Mexico (Corman et al., 2016; Pérez-Hernández et al., 2021), whereas the other clustered with reference sequences derived from marine sediments (Fig. S3).

In the 149-leaf pruned phylogenetic tree derived from an initial 7,224-leaf tree (Fig. 2G), 32 “Crenarchaeota” OTUs were resolved into distinct clades, each clustering with reference lineages from different environmental origins. Three Nitrososphaerota clades were resolved: (i) a putative deep-sea hydrothermal sediment-associated unclassified Nitrososphaeria lineage (seven OTUs, including two novel ones), (ii) the database-defined Nitrososphaeria marine benthic Group A (five OTUs, including one novel), and (iii) a putative deep-sea sediment-associated Nitrosopumilaceae lineage (eight OTUs, including one novel). In addition, three of the seven OTUs affiliated with *Candidatus* Bathyarchaeota also showed closer phylogenetic affinities to marine-derived reference sequences (see Supplementary Table S3).

Representative OTUs from other phyla also exhibited a tendency to cluster more closely with marine-derived lineages, including Planctomycetota, *Candidatus* Falkowiibacteriota, Bacteroidota, and Acidobacteriota (Supplementary methods and results 3, Fig S3-S5). Fig. 2H shows the phylogenetic relationships between selected high-abundance OTUs from the community and their closest reference sequences, with order-level taxonomic affiliations indicated. For example, three OTUs in the Phycisphaeraceae Urania-1B-19 marine sediment group (v, vi, and vii) each exceeded 1% relative abundance in S1. OTU v clustered with the deep-sea sequence KM454365 (Wang et al., 2019), whereas OTUs vi and vii formed another clade with marine-sediment sequences AY627529, JQ580169, and JX227329 (Acosta-González et al., 2013; Heijs et al., 2008; Wu et al., 2013).

These representative OTUs were also queried against the MAPseq/MicrobeAtlas reference framework (Rodrigues et al., 2017; Rodrigues et al., 2026), and their closest MAPseq matches were predominantly associated with marine habitats (Fig. S6). Taken together, the phylogenetic placement and habitat-association patterns suggest that these taxa may represent putative relic marine-associated lineages that persisted through long-term geographic isolation.

### 3.4 Characteristics of reconstructed metagenome-assembled genomes and their distribution

Following metagenomic assembly, binning, and dereplication, MAGs that failed quality assessment using GUNC were excluded. A total of 46 good-quality draft genomes (> 50% completeness and < 10% contamination) were retained (see Supplementary Table S4), including 13 high-quality MAGs (> 90% completeness and < 5% contamination). Clean reads from the four metagenomes showed high mapping rates to the recovered MAGs (52.0% for S1 and > 67% for the other three samples), indicating that the MAG collection provided good representation of the microbial communities.

The relative abundances of individual MAGs across samples, along with their phylogenetic placement in a maximum-likelihood tree constructed with reference genomes (see Supplementary Table S5), are shown in Fig. 3. Of the 46 recovered MAGs, 43 bacterial genomes were assigned to 13 known phyla, one bacterial MAG was classified only at the class level as Deltaproteobacteria with unresolved phylum-level affiliation, and two archaeal MAGs were affiliated with *Candidatus* Woesearchaeota. For MAGs reaching a relative abundance >5% in at least one sample (regarded as “dominant MAG”), the environmental sources of their closest reference genomes were annotated. The MAGs affiliated with Thermodesulfobacteriota exhibited notable genomic diversity, with four out of eight MAGs remaining unassigned to any known genus in the GTDB database. In samples S1, S2, and S4, the cumulative relative abundance of these eight MAGs exceeded that of any other phylum, reaching 19.90%, 25.41%, and 32.17%, respectively (Fig. S7). In contrast, the five Bacteroidota MAGs showed cumulative relative abundances of 5.46%, 10.44%, and 3.56%, and their estimated genome sizes were significantly larger than those of Thermodesulfobacteriota (two-tailed Mann–Whitney test, *p* = 0.011) (Fig. 3).

**Figure 3.**
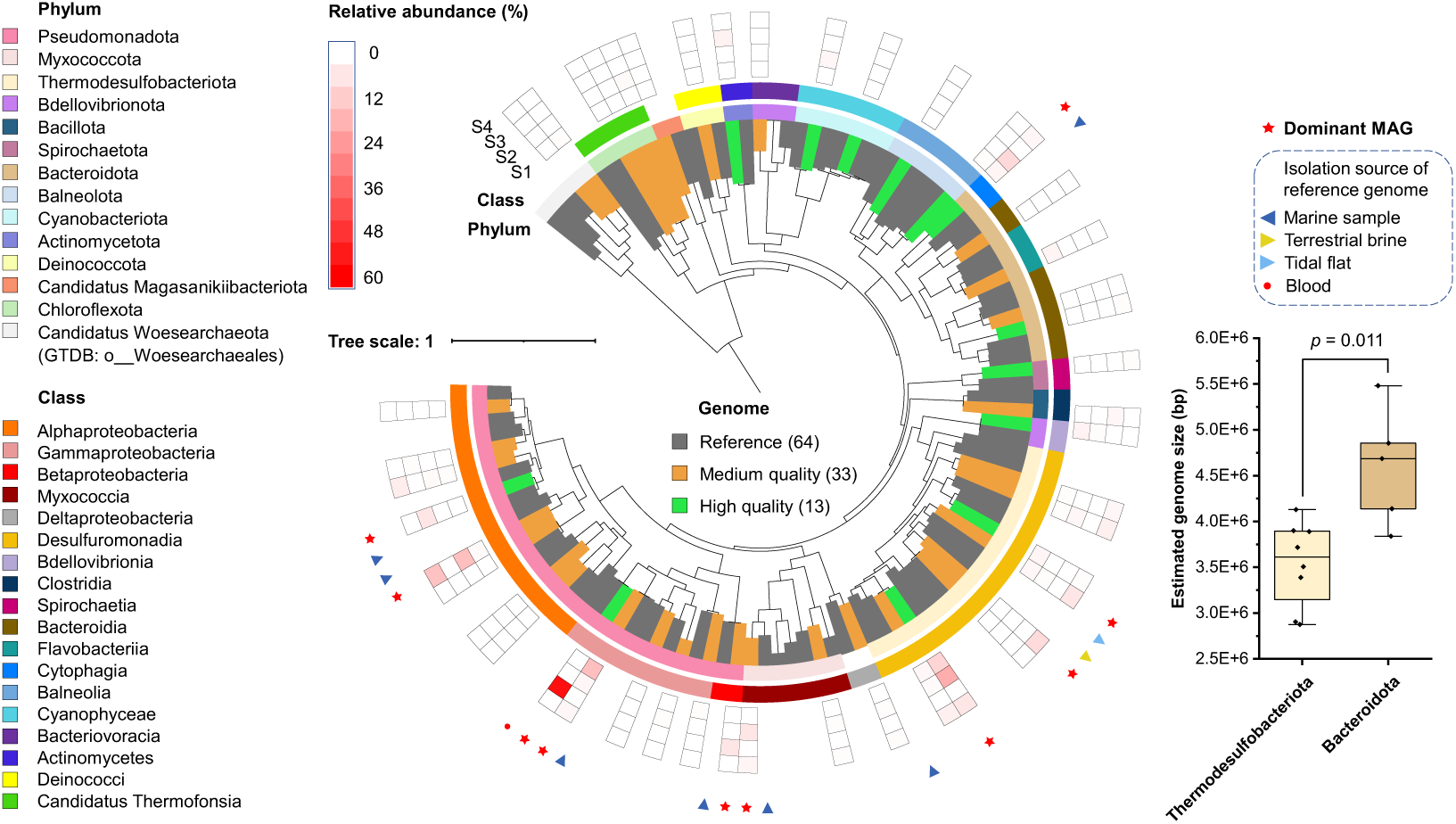
Maximum-likelihood phylogenomic tree of the 46 recovered MAGs and 64 reference genomes. The tree integrates 46 MAGs (high-quality MAGs shown in green; medium-quality MAGs shown in orange) and 64 reference genomes from the NCBI Genome database (gray). A heatmap adjacent to the tree displays the relative abundance of each MAG across samples S1-S4. The environmental origins of the closest reference genomes to dominant MAGs are annotated. A boxplot compares estimated genome sizes between MAGs affiliated with Thermodesulfobacteriota and Bacteroidota (estimated genome size = assembled MAG length / CheckM completeness).

### 3.5 Functional gene profile of microbiomes

Across the 46 MAGs, a total of 156,891 protein-coding genes were predicted, of which 42.6% were assigned to KO identifiers using KofamScan. Based on MAG relative abundances, Thermodesulfobacteriota, Pseudomonadota, and Bacteroidota were considered dominant phyla (Fig. S7). The proportions of genes annotated with KO identifiers differed significantly among these three MAG groups (Kruskal–Wallis test, *p* = 0.0015). Pairwise exact Mann–Whitney U tests with Holm correction showed a progressive decline in annotation coverage from Thermodesulfobacteriota to Pseudomonadota to Bacteroidota. The corresponding KO annotation rates were 51.2%, 47.0%, and 38.8%, respectively. Consistently, Bacteroidota exhibited a significantly lower proportion of COG-annotated proteins than Thermodesulfobacteriota (*p* = 4.57 × 10^-6^, two-tailed *t* test) and Pseudomonadota (*p* = 8.10 × 10^-6^, two-tailed *t* test) (Fig. 4A).

**Figure 4.**
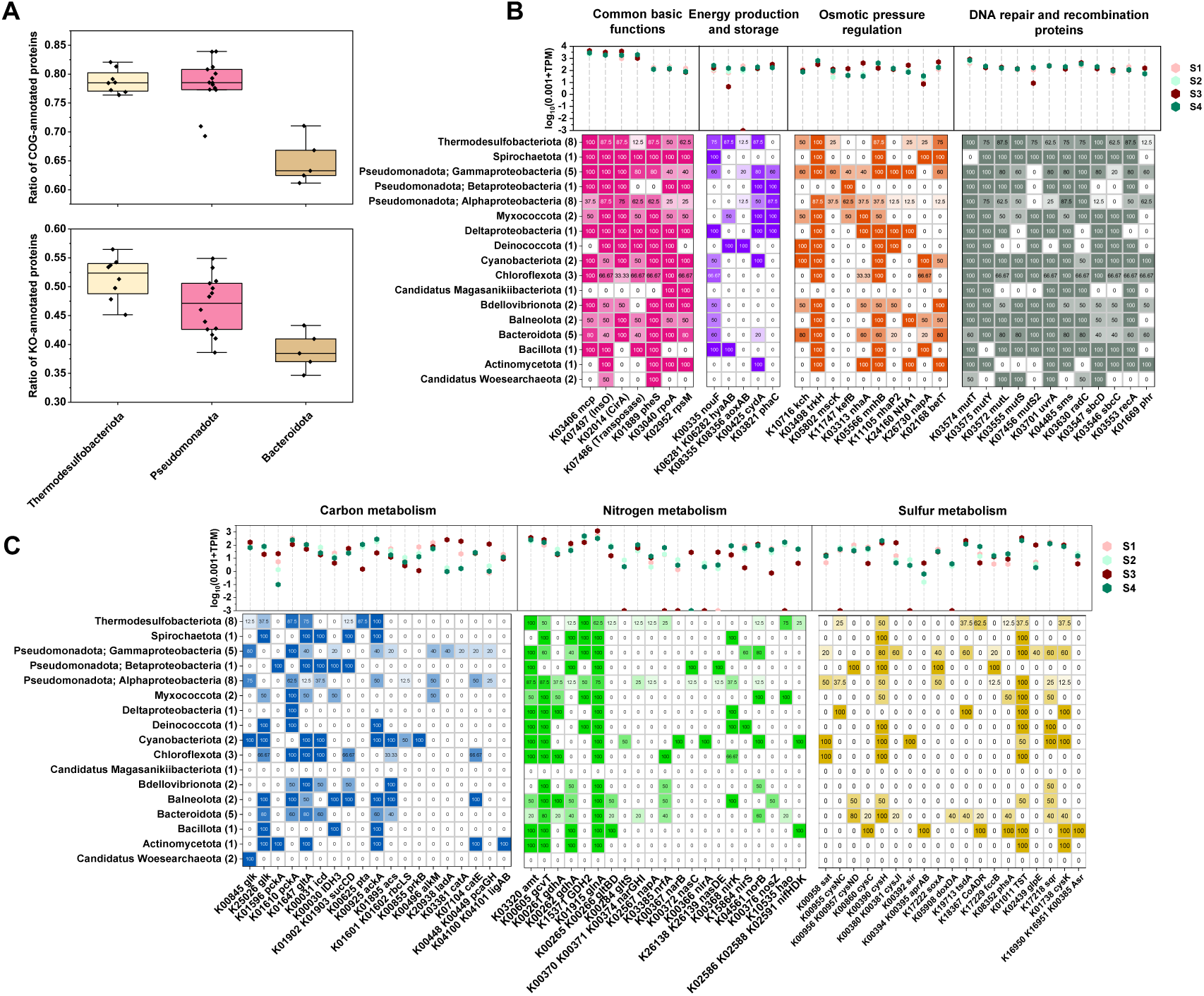
Functional gene profiles of MAGs. (A) Proportions of predicted proteins assigned to COG and KO annotations in MAGs grouped by phylum (Thermodesulfobacteriota, Pseudomonadota, and Bacteroidota). (B) Relative abundances (TPM) and prevalence of functional genes related to energy production and storage, osmotic-pressure regulation, and DNA repair/recombination across MAG lineages. Common reference genes shown for context include: K03406 (*mcp*, methyl-accepting chemotaxis protein), K00335 (*nuoF*, NADH dehydrogenase subunit), K00425 (*cydA*, cytochrome bd ubiquinol oxidase subunit I [EC:7.1.1.7]); single-copy marker genes (K01889, K03040, K02592); selected TCA-cycle markers (K01647, K00030, K00031); and K01915 (*glnA*, glutamine synthetase [EC:6.3.1.2]). (C) TPM level and prevalence of key genes involved in carbon, nitrogen, and sulfur cycling across MAG lineages.

The non-redundant protein-coding gene catalog constructed from assembled contigs comprised 937,632 gene sequences, of which 23.11% were assigned to KO identifiers. Universally conserved, single-copy marker genes “K01889 *pheS*, K03040 *rpoA*, K02592 *rpsM*” recommended by mOTUs1 (Sunagawa et al., 2013) were used as references to contextualize relative TPM levels of functional genes (Fig. 4B). In addition, several genes closely associated with core prokaryotic physiological and biochemical processes could be regarded as functional benchmarks, including *mcp* (K03406; methyl-accepting chemotaxis protein involved in bacterial chemotaxis), *nuoF* (K00335; NADH dehydrogenase subunit), *cydA* (K00425; cytochrome bd ubiquinol oxidase subunit I [EC:7.1.1.7]) for aerobic respiration, key genes for tricarboxylic acid (TCA) cycle (K01647 *gltA*, K00030 *IDH3*, K00031 *icd*), and *glnA* (K01915; glutamine synthetase [EC:6.3.1.2]).

Among all annotated genes, *mcp* exhibited the highest TPM values and occurred in multiple copies in 30 out of the 46 reconstructed MAGs. Additional genes with TPM values of approximately 1,000 or higher across all samples included K07497 (putative transposase; COG2801/InsO), K02014 (iron complex outer membrane receptor protein; COG1629/CirA), and K07486 (transposase; COG3547), suggesting that these genes are broadly distributed and frequently present in multiple copies. Among the four types of cytochrome oxidases associated with aerobic respiration, *cydA* showed the highest TPM levels. *glnA* ranked highest among genes related to coupled carbon and nitrogen cycling. Across all samples, some functional genes reached relative abundances comparable to or exceeding those of the reference genes (*pheS*, *rpoA*, *rpsM*; *nouF*, *cydA* involved in “Energy production and storage”; *gltA*, *IDH3*, *icd* involved in “Carbon metabolism”). These included genes encoding osmoregulation-related channel proteins, such as *kch* (K10716), *trkA* (K03499), *trkH* (K03498), *cvrA* (K11105), *NHA1* (K24160), and *betT* (K02168), as well as genes involved in DNA damage repair pathways (base mismatch repair, nucleotide excision repair, homologous recombination repair, and photoreactivation), including *mutT* (K03574), *mutY* (K03575), *mutL* (K03572), *mutS* (K03555), *uvrA* (K03701), *sms* (K04485), *sbcD* (K03547), *sbcC* (K03546), *recA* (K03553) and *phr* (K01669).

Hydrogen and arsenite were additional potential respiratory electron donors in the community. The TPM values of *hyaAB* (hydrogenase; EC:1.12.99.6) and *aoxAB* (arsenite oxidase; EC:1.20.2.1/1.20.9.1) ranged from 60.97 to 234.59 across S1, S2, and S4. *hya*AB was mainly affiliated with Thermodesulfobacteriota and peaked in S4, whereas *aoxAB* was mainly affiliated with Pseudomonadota and peaked in S1. The TPM values of *phaC*, which mediates the biosynthesis of polyhydroxyalkanoates (PHA), a class of linear polyesters used for carbon and energy storage (Muigano et al., 2025; Obruca et al., 2022; Zou et al., 2017), were generally higher than those of the single-copy marker genes across samples, and were mainly contributed by Pseudomonadota and Myxococcota genomes (Fig. 4B).

Fig. 4C summarizes the prevalence of key elemental cycling genes across different MAG lineages. Compared with Pseudomonadota and Bacteroidota, genomes affiliated with Thermodesulfobacteriota (all assigned to the class Desulfuromonadia) exhibited greater diversity in genes encoding glucose phosphorylation, including K00845, K25026, and K00886. Notably, the three most abundant Thermodesulfobacteriota MAGs all encoded a complete heme biosynthesis module (M00121, 10 genes), whereas this module was incomplete or absent in the remaining five genomes. Only a single genome, S4-bin9 (*Limnobacter profundi*), contained a complete TCA cycle, and only five MAGs encoded a complete module for the conversion of oxaloacetate to 2-oxoglutarate (M00010), consistent with the relatively low TPM values observed for K00030 and K00031. In Thermodesulfobacteriota genomes, the interconversion between acetate and acetyl-CoA was mediated by phosphate acetyltransferase (K00625) and acetate kinase (K00925), rather than by acetyl-CoA synthetase (K01895). In addition, key genes associated with alkane degradation (*alkM* or *ladA*) were identified in several Pseudomonadota MAGs. Genes related to ring-cleavage steps in aromatic compound degradation were present in MAGs from Pseudomonadota, Balneolota, Chloroflexota, and Actinomycetota (*catA*, *catE*, *pcaGH*, *ligAB*).

Three MAGs—two cyanobacterial genomes (S4-bin13, family Cymatolegaceae; S4-bin16, family Oscillatoriaceae) and one *Tranquillimonas* genome (S3-bin3)—were annotated with *rbcL* (K01601), a key gene of the Calvin–Benson–Bassham (CBB) cycle. Based on functional gene annotations from contigs, their genomes are also inferred to encode *rbcS* (K01602) and *prkB* (K00855). In addition, homologs of *rbcLS* and *prkB* most closely related to sequences from *Guyparkeria* or *Caenispirillum* were identified. BLASTp analyses indicated that the two annotated *acsB* genes (K14138) were affiliated with Desulfobulbales and Desulfobacterales, respectively. Although *acsA* (K00198) homologs related to these taxa were detected, the annotated *fhs* (K01938) sequences are more likely derived from Bacillota, Bacteroidota, or Chloroflexota.

Genes involved in nitrogen uptake and assimilation, including *amt* (K03320) and *gcvT* (K00605), were widely distributed across the MAGs, with *amt* occurring in multiple copies in nine genomes. The distribution of genes mediating ammonium assimilation into glutamate/glutamine differed among phyla. For example, Thermodesulfobacteriota genomes appeared to preferentially encode a GDH2-type glutamate dehydrogenase together with glutamine synthetase, rather than glutamate synthase. Consistently across all samples, the TPM values followed the order *GDH2* (K15371) > *gltD* (K00266) > *gltB* (K00265), with mean TPMs of 324.51, 117.66, and 18.67, respectively.

For dissimilatory nitrate reduction (module M00530), *narGHI* was more frequently detected in MAGs than *napAB*, and *nrfAH* was present instead of *nirBD*. One Bacteroidota genome (S4-bin17) encoded a complete M00530 module. Cyanobacteriota genomes (S4-bin13 and S4-bin16) contained the assimilatory nitrate reduction module (M00531; *narB* and *nirA*), whereas two Pseudomonadota genomes (S3-bin8 and S3-bin9) encoded a colinear *nasCDE* gene cluster within this module. A Deinococcota genome (S4-bin26) harbored both *nirK* and *nirS*. Considering the distribution of *norBC* and *nosZ*, including sequences detected on contigs not assigned to MAGs, *Pseudooceanicola* was identified as a potential complete denitrifier. A Balneolota genome (S4-bin30) encoded a clade II *nosZ*, while clade I *nosZ* genes affiliated with Gammaproteobacteria (*Marinobacter* and *Halomonas*) were detected on contigs. In addition, nitrogenase structural genes (*nifHDK*) were identified in genomes affiliated with Cyanobacteriota, Bacillota, and several Thermodesulfobacteriota MAGs.

With respect to sulfur cycling, none of the MAGs encoded a complete pathway for the reduction of sulfate to sulfide, and the SOX system was detected only in a subset of Pseudomonadota genomes. Enzymes such as CoA-dependent NAD(P)H sulfur oxidoreductase and sulfide–cytochrome c reductase were identified and may mediate the interconversion between elemental sulfur and sulfide. Taken together with the distribution of other sulfur-cycling genes, these results indicate that the microbial community harbors a versatile sulfur metabolic network centered on intermediate sulfur species, integrating sulfite and polysulfide reduction, sulfide oxidation, intracellular sulfur trafficking, and assimilatory sulfur incorporation.

### 3.6 Comparative genomic analysis of typical metagenome-assembled genomes

Dominant MAGs affiliated with the genera *Pseudooceanicola*, *Marinobacter*, and *Limnobacter*, as inferred by GTDB-Tk, exhibited their highest average nucleotide identity (ANI) values to reference genomes derived from cultured strains isolated from marine environments (marked in Fig. 3). Among the high-quality MAGs, closest reference genomes related to members of the order Bdellovibrionales and the genus *Gracilimonas* were likewise derived from deep-sea environments (GCA_022450595.1 and GCF_002911695.1, see Supplementary Table S5).

The *Oceanicaulis*-affiliated MAG S2-bin15, with 0% contamination, represents a putative novel species whose GTDB-Tk taxonomic placement is fully defined by topology. It accounted for more than 1.60% relative abundance in samples S1, S2, and S4. A maximum-likelihood phylogeny was reconstructed using this MAG together with 14 *Oceanicaulis* reference genomes in RefSeq (including four isolated strains; strain names also indicated in Fig. 5A), as well as the type strains *Glycocaulis abyssi* LMG 27140 and *Alkalicaulis satelles* G-192 within the family Maricaulaceae. S2-bin15 had a larger estimated genome size (higher than the 2.9–3.3 Mbp range of the 14 *Oceanicaulis* references) and a higher GC content (67.7% versus 61.5– 64.0%), suggesting that it may even represent a novel genus.

**Figure 5.**
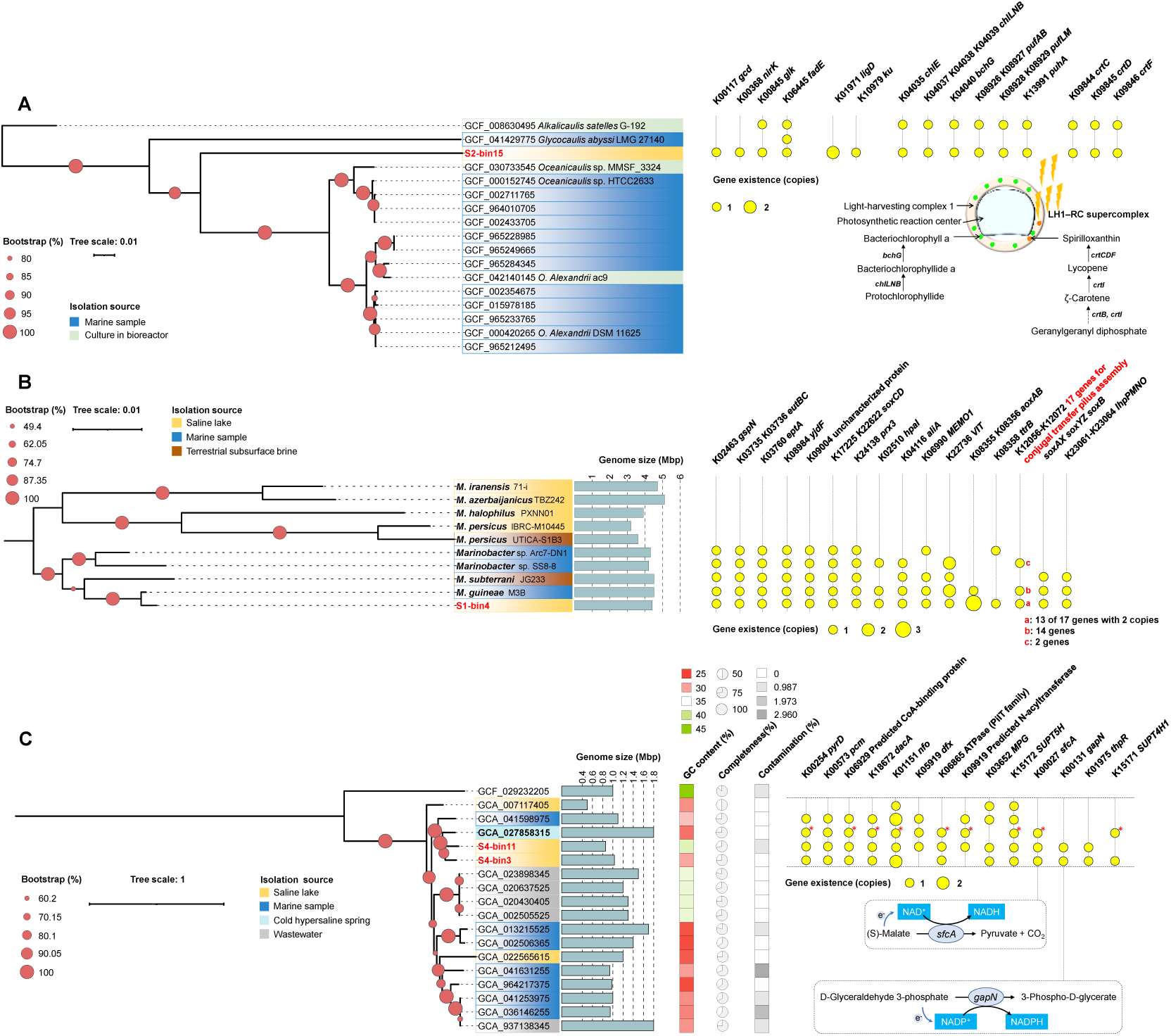
Phylogenetic positions and comparative genomic features of representative MAGs. (A) Species-specific genes and copy numbers in the putative novel *Oceanicaulis* MAG S2-bin15, together with a schematic representation of the proposed LH1–RC supercomplex. (B) Genome size and lineage-specific gene copy numbers of the *Marinobacter guineae* MAG S1-bin4 (showing marine affinity) and reference genomes; additional annotations are provided for the 17 conjugal transfer pilus assembly proteins (K12056–K12072). (C) Comparative genomic features of the two *Candidatus* Woesearchaeota MAGs, S4-bin3 and S4-bin11, and reference genomes; lineage-specific genes and copy numbers in the putative novel branch are indicated, and red “*” marks cases in which the closest homologous protein sequence was derived from archaeon isolate GH56 from the cold hypersaline spring; a schematic of the oxidation of organic compounds is also shown.

Interestingly, functional annotation suggested that some genes uniquely detected in S2-bin15 within “*Oceanicaulis* clade” were exactly also present in *Alkalicaulis satelles* G-192, despite the latter being more distantly related according to single-copy marker genes. For genes encoding quinoprotein glucose dehydrogenase (K00117), glucokinase (K00845), acyl-CoA dehydrogenase (K06445), and the LigD (K01971) and Ku (K10979) proteins involved in the bacterial non-homologous end-joining pathway (Doherty et al., 2001; Weller et al., 2002), BLASTp analysis indicated that their homologs also occurred in several other saline-lake-derived *Oceanicaulis* MAGs, although these genomes are not currently represented in RefSeq. By contrast, no *nirK* homolog from this genus was found in the NR database; the closest matches (also highest percent identity exhibited) were sequences from *Jannaschia formosa* and *Pararhodobacter marinus*, both isolated from marine sediments (Zhang et al., 2019a; Zhang et al., 2019b). S2-bin15 also showed potential for anoxygenic photosynthesis. Genes encoding the reaction center and light-harvesting complex 1 (*pufAB*, *pufLM*, and *puhA*) were also present in other saline-lake-derived *Oceanicaulis* MAGs, and their closest homologs in cultured strains were from *Apolloniradiicaulis salifontis* MS644^T^, isolated from a saline spring near Lake Winnipegosis, Canada, which has been considered a candidate site for Mars-analog life (Messner et al., 2026). The same pattern was observed for genes encoding enzymes involved in bacteriochlorophyll a biosynthesis from protochlorophyllide (*chlE*, *chlLNB*, and *bchG*), as well as genes encoding enzymes responsible for converting lycopene to spirilloxanthin (*crtCDF*). The further conversion of precursor carotenoids to spirilloxanthin may enhance photoprotection under conditions of high light and coexisting oxygen (Niedzwiedzki et al., 2015; Slouf et al., 2012).

The MAG S1-bin4 accounted for more than 1.87% relative abundance in samples S1, S2, and S4. In the maximum-likelihood phylogenomic tree, this genome formed a distinct clade with four *Marinobacter* genomes—*M. guineae* M3B, *M. subterrani* JG233, *Marinobacter* sp. SS8-8, and *Marinobacter* sp. Arc7-DN1—which also represented its closest relatives based on ANI values (97.53, 84.95, 86.70, and 88.19%, respectively). These reference genomes were isolated from marine sediments of the South Shetland Islands (Antarctica), an iron mine borehole in the United States, deep-sea hydrothermal sulfide deposits in the Indian Ocean, and Arctic Ocean sediments at a depth of 580 m, respectively. This lineage therefore appears to represent taxa adapted to subsurface environments, particularly marine subsurface systems, and is distinct from *Marinobacter* strains isolated from continental saline lakes (Fig. 5B).

At the functional gene level, this lineage also exhibited a distinct metabolic profile, including genes encoding secretion pathway protein N (K02463), ethanolamine ammonia-lyase (K03735, K03736), lipid A ethanolaminephosphotransferase (K03760), S-disulfanyl-L-cysteine oxidoreductase (K17225, K22622), glutaredoxin/glutathione-dependent peroxiredoxin (K24138), as well as several proteins of unknown function (K08984, K09004). Additional genes were present in this lineage in a non-universal manner, yet were shared by both *M. guineae* genomes (Fig. 5B). Moreover, the *M. guineae* MAG identified in this study encoded three copies of *aoxAB* (K08355, K08356) and duplicated sets (13 out of 17) of genes encoding conjugal transfer pilus assembly proteins.

*Candidatus* Woesearchaeota was the most abundant archaeal lineage overall, except in S3 where it was less abundant than Nitrososphaerota (Fig. 2B). The highly reduced genomes S4-bin3 (1.03 Mbp) and S4-bin11 (0.86 Mbp) encoded few genes related to major biogeochemical cycling processes in this ecosystem (Fig. 4C). GTDB-Tk assigned both MAGs to the UBA583 group, which is classified at the family level in GTDB. A total of 15 (19 genomes from this family excluding 4 MAGs annotated as bacterial in the NCBI Genome database) MAGs and *Candidatus Nanohalovita haloferacivicina* BNXNv (GCF_029232205) as an outgroup, were used to reconstruct the phylogeny. This family was not included in previous comparative genomic analyses of Woesearchaeota (Huang et al., 2021).

Overall, the environmental origin of the MAGs did not appear to strongly determine their phylogenetic distance. The two MAGs from this study clustered with *Candidatus* Woesearchaeota archaeon isolate GH56 (GCA_027858315), which was isolated from anaerobic sediment in a high-Arctic cold saline sulfur spring (Magnuson et al., 2023) and may represent a high-salinity clade (higher than the average salinity of the ocean), although this genome is smaller and has a higher GC content. They were also closely related to a MAG from coastal bare tidal sandflat sediment (GCA_041598975) (Fig. 5C).

Fig. 5C highlights genes present in this clade but absent from other major branches, which are composed mainly of marine- or wastewater-derived genomes; copy numbers of these genes in GCA_041598975 and in GCA_007117405, a MAG from the Kulunda Steppe saline lake, are also indicated. The protein sequences encoded by these genes in S4-bin3 and S4-bin11 were most closely related to homologs from isolate GH56 in many cases (NR database used). In particular, for *nfo* (K01151; deoxyribonuclease IV), which participates in the base excision repair module, S4-bin3 contained two copies: one, like the single copy in S4-bin11, was most closely related to the isolate GH56 sequence, whereas the other copy was closest to a homolog encoded by a *Candidatus* Peregrinibacteria bacterium MAG from hypoxic seawater (MBT4055861.1). For *dfx* (K05919; superoxide reductase/desulfoferrodoxin), which anaerobic microorganisms use for oxygen detoxification without superoxide dismutase (Jenney et al., 1999), the protein sequence of S4-bin11 (126 aa) showed its highest-identity NR match to the corresponding protein (WP_205242075.1, 124 aa, 70.16% identity) from *Desulfobulbus alkaliphilus*, isolated from anoxic sediments of the Kulunda Steppe saline lake (Sorokin et al., 2012). However, in a phylogenetic tree reconstructed from 274 reference sequences, it tended to cluster with sequences from Methanobacteriati (Supplementary methods and results 4, Fig. S8).

Genes unique to the high-salinity clade included a predicted ATPase containing an N-terminal PIN domain (K06865; COG1855), *sfcA* (K00027; malate dehydrogenase), *gapN* (K00131; glyceraldehyde-3-phosphate dehydrogenase), *thpR* (K01975; RNA 2′,3′-cyclic 3′-phosphodiesterase), and *SUPT4H1* (K15171; transcription elongation factor SPT4), which are implicated in partial oxidation of low-molecular-weight organic compounds, production of reducing equivalents (NADH/NADPH), tRNA splicing, and archaeal transcriptional regulation complexes. These genes broaden the metabolic complexity of the Woesearchaeota UBA583 group, although homologs are also frequently detected in groundwater-derived genomes outside this group. For COG3146, a predicted N-acyltransferase (K09919) prevalent in bacteria, the S4-bin11 homolog exhibited > 40% amino acid identity exclusively to the peptidoglycan biosynthesis protein of isolate GH56 (47.04%), with only four archaeal sequences (372–384 aa) sharing > 20% identity.

### 3.7 Potential horizontal gene transfer events in communities

In this extreme and isolated environment, survival-critical genes may be acquired via HGT, facilitating local adaptation. MetaCHIP identified 94 inter-MAG HGT events. Fig. 6A summarizes gene flow among different classes, with donor and recipient lineages connected by bands whose widths indicate the number of inferred HGT events and whose colors correspond to the donors. The most prominent gene donors were the highly diverse Thermodesulfobacteriota and Proteobacteriota. Fig. 6B shows the functional classification of these transferred genes: nearly half of the encoded enzymes were associated with secondary metabolite biosynthesis. Desulfuromonadia was the class contributing the largest number of genes encoding enzymes or transporters, whereas Gammaproteobacteria was the major donor of ribosomal protein genes. In addition, 19.1% of the transferred genes remained poorly characterized (no KO assigned) and were annotated to functional categories including oxidoreductases, membrane proteins, response regulators, and others.

**Figure 6.**
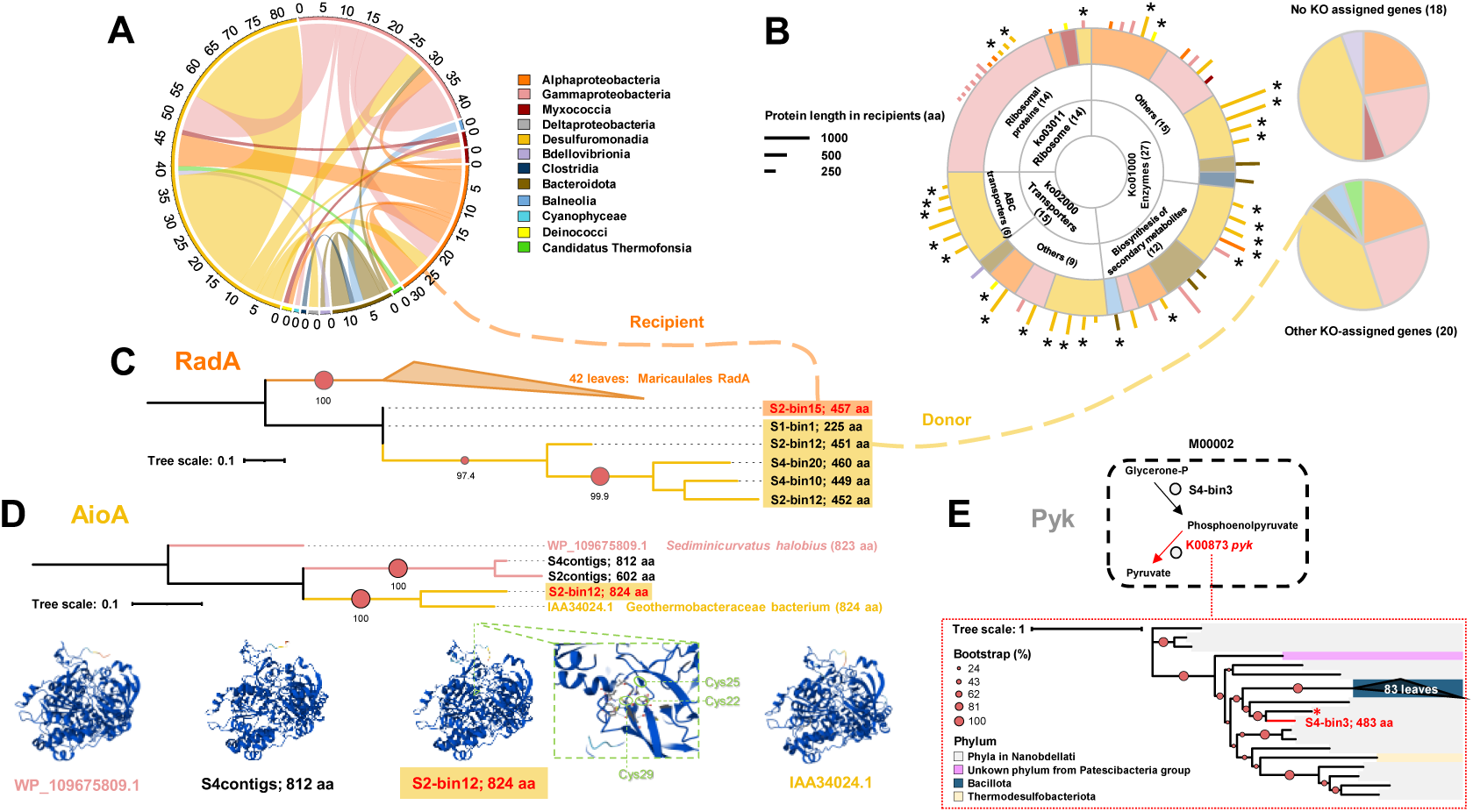
Potential HGT events in microbial communities. (A) MetaCHIP-inferred gene flow among MAGs at the class level. Bands connect donors and recipients; band width is proportional to the number of HGT events, and band color indicates the donor class. (B) KEGG-based functional classification of the putatively transferred genes. KO-assigned genes were primarily associated with three functional categories: enzymes, transporters, and ribosomal proteins. In the sunburst plot, sector size represents the number of genes assigned to each functional category, inner-sector colors indicate inferred donor classes, and the external bars show the protein lengths of the corresponding genes in recipient MAGs. Bar colors denote recipient classes. “*” marks high-similarity transfers (donor–recipient homologues sharing >80% amino acid identity). The two pie charts summarize donor-class contributions for other KO-assigned genes and KO-unassigned genes, respectively. (C) Maximum-likelihood phylogeny of RadA supporting the putative acquisition of a *radA* homologue by S2-bin15 from a Desulfuromonadia-related donor. (D) Example of a putative HGT event without an unambiguous MAG-resolved donor. The pruned maximum-likelihood phylogeny of AioA suggests that S2-bin12 may have acquired an *aoxB* homologue from a Gammaproteobacteria-related lineage. The tree includes the S2-bin12 sequence, two unbinned contig-derived homologues, and closely related reference sequences from the NR database. AlphaFold 3 structural models are shown for four AioA proteins retaining both the [3Fe–4S] cluster and the molybdopterin oxidoreductase domain. The conserved [3Fe–4S] cluster-coordinating motif, Cys-X2-Cys-X3-Cys, is highlighted in the S2-bin12 AioA model. (E) Maximum-likelihood phylogeny of S4-bin3 *pyk*, which retains a complete M00002 module (glycerone-P to pyruvate) and may have acquired *pyk* via HGT from Bacillota (the reference sequence marked with red “*” is from archaeon isolate GH56).

Some genes involved in DNA damage repair may also have been transferred, including the detected DNA repair protein RadA/Sms (K04485). Forty-two alphaproteobacterial *radA* sequences similar to the homolog from S2-bin15, (26 identified by BLASTp against the NR database, the other 16 *radA* from reference genomes shown in Fig. 5B), together with *radA* sequences from five Desulfuromonadia MAGs recovered, were used to reconstruct the phylogenetic tree. As shown in Fig. 6C, the *radA* sequence from S2-bin15 clustered with Desulfuromonadia *radA* sequences rather than with those from Alphaproteobacteria.

Of course, MetaCHIP could not capture all HGT events, because the true donors may not be included in the MAG set. Phylogenetic analysis of *aoxB* suggested that the contig-derived *aoxB* sequences were mainly affiliated with Alphaproteobacteria and Gammaproteobacteria. In particular, *aoxB* from the Desulfuromonadia MAG S2-bin12 appeared to have been acquired from Gammaproteobacteria or Betaproteobacteria via HGT, as indicated by both the tree topology and BLASTp searches. This interpretation is further supported by the fact that homologs of this gene are exceptionally rare in known Thermodesulfobacteriota genomes (Supplementary methods and results 5, Fig. S9). The closest sequence to this *aoxB* was IAA34024.1 from a Geothermobacteraceae bacterium MAG recovered from an iron-rich hot spring, an environment that has been proposed as an analog of the pre-Cambrian surface system (Li-Hau et al., 2025). Fig. 6D shows a pruned maximum-likelihood tree including this specific clade as well as two contig-derived *aoxB* sequences that were not binned into MAGs. According to protein structure prediction (pTM score = 0.97, indicating high confidence in the overall model), identified *aoxB* from the Desulfuromonadia genomes together with its close contig-derived homolog, encoded AioA that were structurally similar to the corresponding subunit from Proteobacteria, represented by WP_109675809.1 from *Sediminicurvatus halobius*. The [3Fe–4S] cluster is located in domain I and is coordinated by a conserved cysteine-rich motif (CHFCIVGC for both *Sediminicurvatus halobius* and S2-bin12), three cysteines highlighted in Fig. 6D.

Another representative case was pyruvate kinase (*pyk*, K00873) in the *Candidatus* Woesearchaeota MAG. S4-bin3 retained a complete “glycolysis core module involving three-carbon compounds” (M00002). In the phylogenetic analysis, *pyk* clustered with sequences from two Nanobdellati archaeal MAGs (MDA3856203.1 and MDH3353556.1) and 83 pyruvate kinase homologs of Bacillota (bootstrap = 68%), most of which were from cultured strains, rather than with the other Nanobdellati-dominated clades (Fig. 6E). This pattern indicates potential gene transfer from Bacillota to *Candidatus* Woesearchaeota.

### 3.8 Characteristic metabolic patterns of species in communities

The ecological basis underlying the dominance of these MAGs can be inferred from their relative enrichment in four functional gene categories—high growth yield, resource acquisition, stress tolerance, and osmotic pressure regulation (Supplementary methods and results 6; Fig. S10)—and from their distinct metabolic features relative to closest phylogenetic references, particularly regarding substrate utilization and element cycling.

Rather than exhibiting a uniform adaptive strategy, dominant MAGs displayed complementary functional characteristics that may collectively support microbial persistence under the extreme conditions of Eboliang Hu saline lake system. Several abundant genomes showed enhanced capacities for resource acquisition through diverse organic matter utilization pathways. For example, multiple Pseudomonadota MAGs (*M. guineae* S1-bin4, *Pseudooceanicola* S3-bin8, and *Pseudomonas alloputida* S3-bin2) encoded expanded pathways involved in aromatic compound degradation, suggesting a potential role in exploiting chemically diverse organic substrates. In contrast, Thermodesulfobacteriota MAG S1-bin1 exhibited strong adaptations related to nitrogen acquisition (Fig. S16) and osmotic regulation (Fig. S10), including expanded ammonium assimilation capacities and abundant genes associated with salt stress tolerance.

Comparisons with closely related reference genomes further revealed lineage-specific functional innovations. Several dominant MAGs possessed additional metabolic modules, increased copy numbers of key genes, or unique regulatory and stress-response capacities absent from their nearest relatives. For example, compared with 14 larger reference Flammeovirgaceae genomes (Fig. S24), the Flammeovirgaceae MAG S2-bin24 additionally encoded a tryptophan 2-monooxygenase (implicated in L-tryptophan oxidation), a multicomponent Na^+^/H^+^ antiporter complex (mnhA–G), and a suite of CRISPR-associated proteins corresponding to Type I (K19114, K19115, K19116) and Type III (K07016, K09002, K19138, K19139, K19140) CRISPR–Cas systems (Fig. S17). These genome-specific features suggest that local environmental pressures have shaped distinct adaptive trajectories after lineage diversification.

Therefore, community dominance in Eboliang Hu ecosystem is likely maintained through a combination of ecological niche differentiation and evolutionary adaptation (for detailed comparative analyses of individual dominant MAGs and their associated genomic traits, see Supplementary methods and results 6).

## 4. Discussion

### 4.1 The unique isolated aquatic environment preserves a microbial community that exhibits partial marine affinity

The microbial community in this unique aquatic ecosystem may be jointly structured by osmotic stress, strong UV radiation, reduced nitrogen availability, sulfur transformations, and limited input of terrestrial organic carbon, resembling an “island in the desert.” This setting differs from most alkaline saline lakes on the Tibetan Plateau (Cheng et al., 2024) and provides a useful natural system for examining how microbial communities persist under combined salinity, desiccation, UV exposure, and nutrient limitation. The sediment community of Eboliang Hu saline lakes in this study, by comparison, appears more similar to a typical saline-lake sediment community because of the enrichment of Thermodesulfobacteriota (Fig. S7). Nevertheless, at higher taxonomic resolution, even S2 and S4 which shared similar dominant phyla showed pronounced small-scale heterogeneity (Fig. 2F), underscoring the distinctiveness of this community.

At the 16S rRNA gene level, Planctomycetota, Patescibacteria, and *Candidatus* Woesearchaeota each became strongly enriched at particular sampling sites (>30% relative abundance), yet no correspondingly diverse MAGs were recovered for these lineages in this study. These Patescibacteria groups may preferentially inhabit oligotrophic groundwater environments (Herrmann et al., 2019) and often engage in symbiotic interactions with other bacteria (Chaudhari et al., 2021; Nakajima et al., 2025). Among Planctomycetota, Pirellulales are known to possess one of the highest encoded hydrolytic potentials (Klimek et al., 2024), and their relatively high abundance at two sampling sites may indicate greater polysaccharide availability. Considering the marine affinity of the dominant Phycisphaerales OTUs (Fig. 2H), their capacity to degrade complex organic compounds (Lenferink et al., 2024) may have been an important factor supporting their long-term persistence in this ecosystem.

By broadly comparing gene and protein sequences against public databases, we revealed both the novelty of the community members and a phylogenetic pattern in which some taxa showed marine affinity. This pattern was evident across multiple taxonomic and functional levels, including archaeal and bacterial 16S gene sequences (Fig. 2G, Fig. 2H), eukaryotic 18S gene sequences (Supplementary methods and results 12, Fig. S2), diverse bacterial MAGs from different phyla, and protein sequences encoded by functionally relevant genes potentially linked to environmental adaptation (Supplementary methods and results 7, 10). However, it should be noted that some of the closest reference sequences were derived from other habitats, such as groundwater, deserts, and other saline lakes, highlighting the complex sources of the Eboliang Hu lake microbiome. The evolutionary history of species entering this environment likely differs substantially in duration and trajectory, and dispersal mediated by water birds or groundwater infiltration may have contributed to colonization even in this type of isolated, island-like extreme ecosystem (Schiwitza et al., 2021). Notably, compared with several other typical hypersaline inland lakes, the eukaryotic community composition suggests that the degree of isolation in the Eboliang Hu lake may be more extreme, with no enrichment of brine shrimp (Gao et al., 2026) or eukaryotic algae (Al-Daghistani et al., 2024; Casamayor et al., 2013; Wang et al., 2014). Molecular clock analyses yield a divergence time of 10.5 million years ago (mya) for the two *Marinobacter guineae* genomes (Supplementary methods and results 11, Fig. S26), which is comparable to the ∼11.0 mya divergence inferred previously for *Oceanicaulis* (Marin et al., 2017). These estimates together indicate that the relevant *Marinobacter*- and *Oceanicaulis*-related lineages (Fig. 5) diverged within a ∼10–11 mya window, temporally overlapping with the late-Miocene uplift of the western Qaidam Basin (Miao et al., 2022). Such a correspondence is consistent with a scenario in which regional uplift and basin reorganization promoted the isolation and subsequent diversification of these lineages.

Future, more extensive hydrochemical surveys and microbial sampling around the Eboliang area may help further clarify the environmental factors shaping the microbial diversity observed in this study, as well as the extent of marine affinity among community members. At the same time, such efforts should be carefully designed to minimize disturbance to active indigenous microbial communities and to prevent degradation of microbial fossils or ancient DNA preserved in the sediments.

### 4.2 The genomic functional profile indicates the flexible and cooperative metabolic patterns of the community

The functional profile of Eboliang Hu microbiome points to a biogeochemical network dominated by metabolic flexibility rather than by a single complete canonical pathway (Fig. 7). The metabolic pathways encoded by the dominant MAGs appear to cover the core material cycling processes of the community, whereas the less abundant genomes may still play important roles in energy input and inorganic carbon fixation within the system.

**Figure 7.**
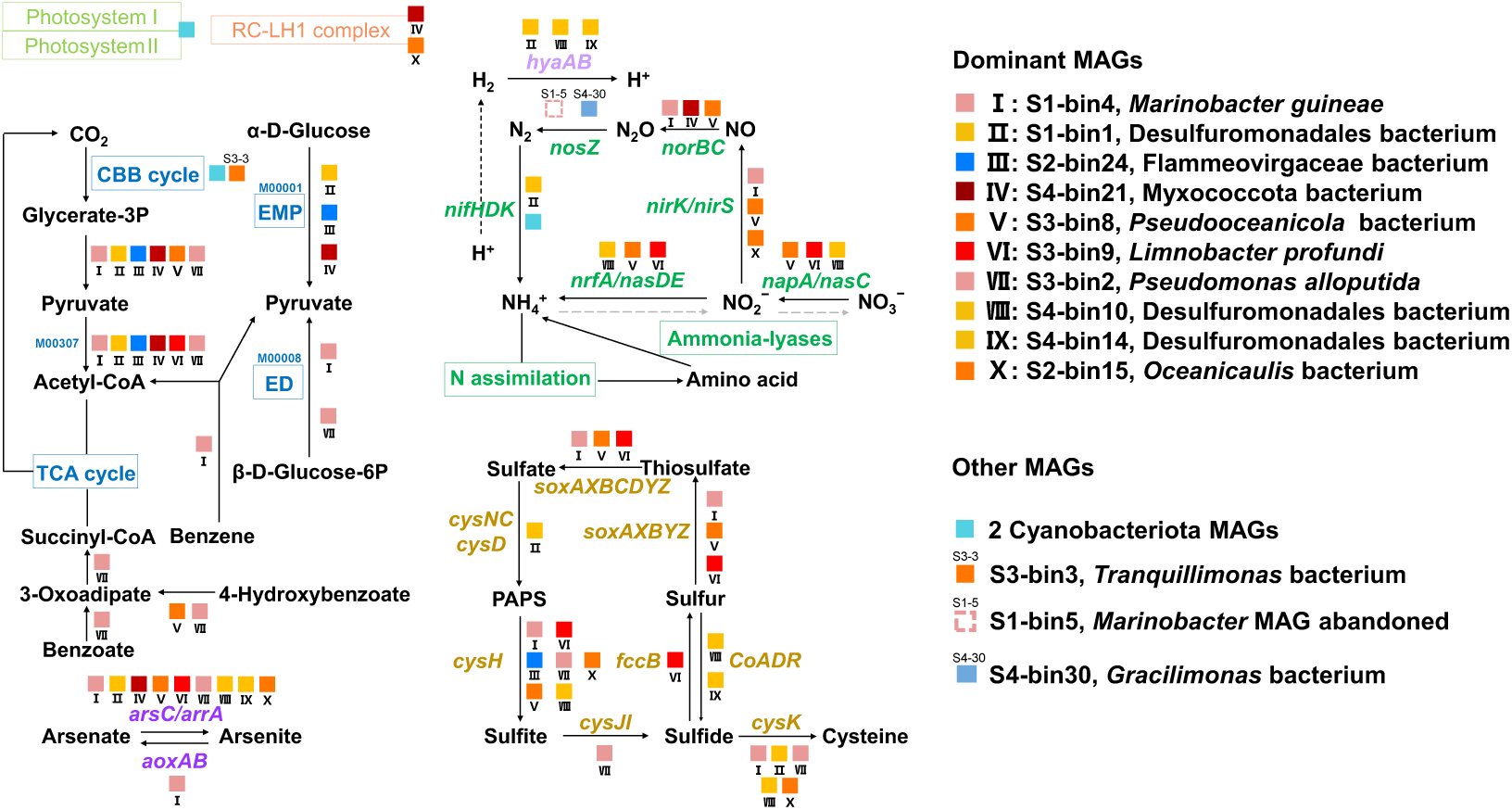
Schematic representation of major MAGs implicated in phototrophic systems and carbon, nitrogen, sulfur, arsenic, and hydrogen metabolism within the community. No MAGs contained confidently annotated ammonia monooxygenase or nitrite oxidoreductase genes; thus, the corresponding transformation steps are depicted as gray dashed lines. The S1-bin5 *Marinobacter* MAG was excluded after failing GUNC quality assessment but supported *nosZ* presence in this genus.

Carbon metabolism appeared to rely on partial and lineage-specific routes, including sugar catabolism, incomplete or selectively retained TCA-cycle modules, PHA biosynthesis, and degradation of aromatic or alkane-derived substrates (Fig. 4C). Collectively, these features suggest that the dominant lineages are capable of utilizing chemically diverse carbon substrates. Such versatility may be advantageous in environments where carbon availability fluctuates over space and time, enabling microorganisms to exploit transient resource inputs and maintain cellular carbon reserves through PHA accumulation (Mason-Jones et al., 2023).

In particular, the complex nitrogen and sulfur cycles likely require cooperation among multiple taxa. Nitrogen cycling was biased toward assimilation and reductive transformations. The relative high ammonium concentrations and the near absence of nitrate and nitrite in the water phase are consistent with the genomic enrichment of ammonium transporters and glutamate/glutamine biosynthesis routes. The higher abundance of *GDH2* relative to *gltBD* suggests that direct ammonium incorporation via glutamate dehydrogenase may be particularly important under the observed geochemical conditions. At the same time, nitrate reduction, nitrite reduction, denitrification-related steps and nitrogen fixation are distributed across different lineages rather than concentrated in a single dominant taxon. Although OTUs assigned to Nitrososphaerota and Nitrospirota were detected, no corresponding MAGs were recovered, and no high-confidence *amoA* or *nxrA* genes were identified in the contigs. This likely reflects the very low abundance of these taxa, which hindered their recovery. Sulfur metabolism also appeared modular. The absence of a complete sulfate-to-sulfide reduction pathway in recovered MAGs argues against canonical dissimilatory sulfate reduction as the dominant sulfur process captured by the MAG set. Instead, the community-level sulfur network centered on intermediate sulfur species may be particularly relevant in shallow saline sediments where redox gradients, sulfide availability and organic-matter inputs fluctuate over small spatial scales (Farias et al., 2017; Fernandez et al., 2016).

Hydrogen oxidation has been highlighted as an important contributor to primary production in desert soils, and it can be further stimulated by hydration events (Jordaan et al., 2020; Liu et al., 2022). In this context, the electron-acceptor preferences of the novel Thermodesulfobacteriota genomes recovered from these sediments represent an interesting question that merits further investigation using additional approaches. Their cytochrome bd ubiquinol oxidase (Fig. 4B) may help alleviate oxygen stress (Ramel et al., 2013), whereas under anoxic conditions they appear to invest more broadly in arsenate reduction (Supplementary methods and results 13, Fig. S27), with TPM values even exceeding those of genes related to nitrate or nitrite reduction. Such flexibility in the use of different electron donors and acceptors may help metabolic generalists expand their ecological niche (Chen et al., 2021), as exemplified by S1-bin1, which exceeded 5% relative abundance in three samples. Under the weakly acidic pH range observed here (Fig. 2A), the thermodynamic ranking of electron acceptors can be summarized as nitrate > nitrite > arsenate >> sulfate, although physiological preferences do not necessarily follow this order exactly (Rauschenbach et al., 2012); nevertheless, arsenate respiration may represent an important strategy. Notably, all four MAGs encoding *aoxAB* also encoded arsenate reductase genes, which may likewise reflect redox flexibility in the sediments.

The overall fraction of protein-coding genes assigned KO identifiers was below one half for the MAGs, and it was even lower at the contig level. Compared with Thermodesulfobacteriota, Bacteroidota showed larger genome sizes (Fig. 3) but lower annotation rates for protein-coding genes (Fig. 4A), largely because of genome novelty. Indeed, four MAGs, including the dominant S2-bin24, had no GTDB reference genome with ANI >80%, which likely limited the availability of homologous references for annotation. Future integration of protein language models, structural prediction, and experimental validation will be essential for resolving the ecological roles of these uncharacterized genes and for expanding our understanding of microbial extremozymes (Chasapi et al., 2026; Jiang et al., 2026; Szydlowski et al., 2024).

### 4.3 Co-evolutionary history and HGT events shape environmentally adaptive genome functions

The recent metagenomic surveys of high-salinity soils in the Qaidam Basin Mars-analog desert did not recover any *Candidatus* Woesearchaeota MAGs (Liu et al., 2025), whereas these genomes have been detected in lake sediments from other regions of the Qinghai–Tibet Plateau (Zhang et al., 2025), suggesting some environmental preference. A close phylogenetic relative of the two MAGs recovered in this study was from Kulunda saline-lake sediment metagenomes, where *Phormidium* strains and diverse Thermodesulfobacteriota are also present (Vavourakis et al., 2018). In addition, phylogenetic analysis of S4-bin13 placed it within a “*Dactylococcopsis* + *Halothece*” clade (Supplementary Methods and Results 8, Fig. S23), whose other six MAGs were all recovered from sediments or microbial mats in hypersaline brines from continental arid regions (Gutiérrez-Preciado et al., 2024; Gutiérrez-Preciado et al., 2018; Kurth et al., 2021; McGonigle et al., 2019; Saghaï et al., 2017). Despite differences in pH, salinity, and moisture regime, these samples showed broadly similar community compositions, with *Candidatus* Woesearchaeota, Thermodesulfobacteriota, Chloroflexota, *Candidatus* Magasanikiibacteriota, Spirochaetota, Deinococcota, and Bradymonadales (Deltaproteobacteria) recurrently detected (though not necessarily all in every sample).

Collectively, these patterns suggest that similar assemblages recur in comparable extreme environments; however, no two geographically distant sites show the highest phylogenetic similarity across multiple phyla simultaneously. This indicates a long shared evolutionary history, followed by geographic isolation at different time points and subsequent maintenance of functionally coupled associations under environmental selection. Nevertheless, the low biomass typical of the drier shallow-water sites makes it especially difficult to extract sufficient nucleic acids, and the scarcity of multi-omics datasets currently limits our ability to resolve the interspecific interaction mechanisms within such putative symbiotic consortia based on genome annotation alone.

Under this co-occurrence pattern, some functional genes may be particularly important, including those shown in Fig. 4B to have higher TPM values than the corresponding reference genes. These genes are involved in chemotaxis (for microscale resource tracking), iron uptake, transposase activity, and strategies that directly help microorganisms resist high osmotic pressure and UV radiation, indicating that the community has substantial capacity to withstand selective pressures and that its genomes are highly plastic (Fambrini et al., 2020). HGT events promoted extensive interspecific gene flow, such as exchange of transporter genes within Desulfuromonadia, whereas the secondary-metabolite biosynthetic capacity of Desulfuromonadia was additionally acquired by Proteobacteria members (Fig. 6B). HGT is considered an important mechanism for rapidly acquiring adaptive functions in extreme environments. Compared to relying on de novo mutations, directly obtaining genes that are already suitable for the environment is faster and more cost-effective (Zhaxybayeva and Nesbo, 2026).

Comparative genomic analyses revealed both functional similarity and divergence among closely related genomes inhabiting high-salinity environments, and the latter may reflect the stronger selective pressures imposed by extreme habitats such as the one studied here. Taking *Candidatus* Woesearchaeota and Cyanophyceae as examples, genomes within the high-salinity clade retained several features absent from their closest relatives (Fig. 5C; Supplementary methods and results 8, Fig. S23); if these traits are interpreted as vertically inherited from a common ancestor, they would have been differentially retained or lost during evolution at higher taxonomic levels. A recent study of archaeal co-evolution in halophiles suggested that genome-reduced halophilic archaea may use sister taxa as hosts and acquire essential survival genes via HGT, but no clear signal of dependence on other archaea was detected for *Candidatus* Woesearchaeota (Hamm et al., 2026). This may be because they are not restricted to single, extreme salt or thermal environments (Huang et al., 2021), and environmental selection did not force them to repeatedly acquire a complete suite of adaptive genes from a specific archaeal host. Nevertheless, even *kefB* that helps extremophilic halophilic archaea maintain osmotic balance (Hamm et al., 2026), was absent from the two size-reduced *Candidatus* Woesearchaeota MAGs recovered here (Fig. 4B), strongly suggesting host dependence; we also identified a *pyk* homolog that may have been acquired from Bacillota (Fig. 6E). Future expansion of the high-salinity clade dataset will help determine whether Bacillota serve as specific hosts for *Candidatus* Woesearchaeota under particular environmental conditions.

Two other illustrative cases are the *Marinobacter* and *Oceanicaulis* MAGs, which remain most closely related to present-day marine genomes. For *Marinobacter*, S1-bin4 may have retained marine-lineage genes that were common at the divergence node because they conferred adaptive advantages, such as *aoxAB*, which may expand the ecological niche by supporting energy generation (Fig. 5B; for other genes, they may help with substrate acquisition, transport or/and resistance to environmental stress, see Supplementary Table S8). In contrast, S2-bin15 differs substantially from extant marine *Oceanicaulis* lineages in functional gene profile and appears to have adopted a distinct lifestyle; its superior DNA repair capacity and the emergence of “saline-lake-specific” genes might have resulted from early HGT events (Fig. 5A).

Arsenic concentrations in the ancient crust were likely much higher than they are today, and arsenite has been proposed as a candidate electron donor for the chemolithotrophic growth of early life (Rascovan et al., 2016). Consequently, enrichment and microbial utilization of arsenic in ancient marine sediments may have been widespread. In addition, arsenic methylation products have been suggested to contribute to pressure tolerance in deep-sea organisms (Hoffmann et al., 2018; Li et al., 2024), potentially favoring the long-term maintenance of arsenic-related metabolisms. These factors may have facilitated the persistence of arsenic-metabolism genes in the ancestors of the sediment microbiome studied here before regional uplift. Arsenic concentrations in high-altitude saline lakes can also be elevated through the gradual release of trace elements from minerals during weathering (Kurth et al., 2017), and enrichment of arsenic-metabolism genes has been reported in comparable low-oxygen saline habitats (Sancho-Tomás et al., 2018; Veloso et al., 2023). Retention of *aoxAB* may therefore represent an important strategy for adapting to this environmental transition (though without quantitative measurements of arsenic species in the current sediments), particularly in *Marinobacter*, *Roseovarius*, and Trueperaceae observed here (Fig. S9). Notably, *aoxB* is extremely rare in Desulfuromonadia. Combined evidence from phylogenetic analyses and protein structure prediction suggests that this lineage MAGs could acquire a functional version of this enzyme from Proteobacteria through HGT events to broaden electron-donor use (Fig. 6D).

Comparative analyses between the dominant MAGs and their closest reference genomes revealed not only additional genes related to substrate utilization and DNA repair, but also, in particular for Cyanophyceae and Flammeovirgaceae, the expanded CRISPR-Cas systems (Supplementary methods and results 8, 9; Fig. S17), likely reflecting strong selection imposed by abundant viruses and mobile genetic elements (Bhattarai et al., 2021; Meaden et al., 2022; Roberts et al., 2025). These lineage-specific advantages highlighted by cross-genome comparisons may help generate hypotheses about their symbiotic modes; for example, salt-tolerant Cyanophyceae might function not only as primary producers but also as contributors to stress resistance (S strategy), whereas Spirochaetota may act as providers of metabolic substrates for their partners (A strategy) (Fig. S10).

### 4.4 Implications for extraterrestrial biology

The specific microbial assemblages preserved in the Eboliang Hu saline lake sediments and analogous hypersaline environments provide compelling terrestrial models for refining astrobiological exploration strategies, particularly for targeting past or extant life in Martian brines. Importantly, the mechanism by which long-term localized hyperaridification shapes the genomic plasticity of these microbial communities presents an evolutionary trajectory highly analogous to that of Mars. The red planet transitioned from a volatile-rich, actively hydrologic system during the Noachian epoch, through a period marked by localized evaporitic basins in the Hesperian, and ultimately into its current hyperarid, frozen Amazonian state (Carr and Head, 2010).

Recent Martian geomorphological and mineralogical surveys have further solidified these analog links. The continuous mapping of paleoshorelines and scarp-fronted deltaic deposits across the northern lowlands and within Valles Marineris provides robust lines of evidence for an ancient, planet-wide ocean system (Li et al., 2025a), which would have left extensive sedimentary records. Notably, NASA’s Curiosity rover recently identified a diverse array of complex, preserved organic compounds—including benzothiophene and nitrogen-bearing structures reminiscent of nucleotide precursors—entombed within 3.5-billion-year-old clay-rich lacustrine mudstones at Gale Crater (Williams et al., 2026). This directly echoes the biosignature preservation dynamics observed in terrestrial distal alluvial fans and aqueous deposits experiencing limited surface degradation under severe desiccation in the Qaidam Basin (Chen et al., 2022).

Furthermore, the genomic enrichment of ancient chemolithotrophic pathways observed in this study, specifically hydrogen oxidation and arsenate/arsenite utilization, aligns with the putative chemical energetic landscape of early Mars (McCollom et al., 2022; Tarnas et al., 2018). Although in situ data on the availability of Martian arsenic species remain sparse, arsenic-driven metabolisms have long been proposed as potential energetic pathways for primitive extraterrestrial ecosystems (Oremland and Stolz, 2003). In such anoxic, hypersaline, and potentially heavy-metal-enriched planetary subsurfaces, life may favor the highly flexible, modular, and symbiotic energy-cooperative networks demonstrated by the Eboliang Hu microbiome. Consequently, the recurrent microbial consortia identified here—comprising Thermodesulfobacteriota, Pseudomonadota, and genome-reduced *Candidatus* Woesearchaeota—represent ideal candidates not only for validating computational models of planetary habitability but also as foundational chassis for synthetic ecology and bioreactor engineering in future extraterrestrial colonization and in-situ resource utilization (ISRU) frameworks (Starr and Muscatello, 2020).

## 5. Conclusion

Eboliang Hu saline lake hosts a diverse and spatially heterogeneous microbial community shaped by extreme salinity, nutrient limitation, and long-term geographic isolation. Genome-resolved metagenomics recovered 46 good-quality MAGs spanning major bacterial and archaeal lineages, with Thermodesulfobacteriota and Pseudomonadota contributing substantially to community abundance and functional potential. Across sites, microbial assemblages exhibit strong microhabitat differentiation despite shared dominant phyla, indicating fine-scale environmental filtering. Functional reconstruction reveals a distributed metabolic architecture in which no single lineage encodes complete biogeochemical pathways. Instead, carbon, nitrogen, and sulfur cycling are partitioned across taxa, relying on complementary and partial pathways. Key processes include versatile carbon substrate utilization, nitrogen assimilation centered on ammonium, and sulfur transformations mediated by intermediate oxidation states. Genes associated with stress tolerance, DNA repair, and osmotic regulation are widely distributed, consistent with adaptation to hypersaline and UV-exposed conditions. Phylogenetic analyses consistently indicate that multiple dominant taxa are closely related to marine- and subsurface-associated lineages, suggesting long-term biogeographic connectivity rather than purely local diversification. Divergence time estimates for *Marinobacter*- and *Oceanicaulis*-related clades coincide with late-Miocene tectonic uplift of the western Qaidam Basin, supporting a scenario in which basin reorganization contributed to lineage isolation and subsequent evolutionary divergence. HGT further contributes to functional innovation, particularly in genes related to energy metabolism and environmental adaptation, reinforcing a model of ongoing co-evolution among coexisting lineages. Future work integrating time-resolved geochemistry and broader spatial sampling will be essential to disentangle the relative roles of historical contingency and contemporary selection in shaping this extreme ecosystem.

## Supporting information

Supplementary methods, results, tables, figures; files of main figures

## Supplementary information

E-supplementary data for this work can be found in e-version of this paper online.

## References

Abramson, J., Adler, J., Dunger, J., Evans, R., Green, T., Pritzel, A., Ronneberger, O., Willmore, L., Ballard, A., Bambrick, J., Bodenstein, S., Evans, D., Hung, C., O’Neill, M., Reiman, D., Tunyasuvunakool, K., Wu, Z., Zemgulyte, A., Arvaniti, E., Beattie, C., Bertolli, O., Bridgland, A., Cherepanov, A., Congreve, M., Cowen-Rivers, A., Cowie, A., Figurnov, M., Fuchs, F., Gladman, H., Jain, R., Khan, Y., Low, C., Perlin, K., Potapenko, A., Savy, P., Singh, S., Stecula, A., Thillaisundaram, A., Tong, C., Yakneen, S., Zhong, E., Zielinski, M., Zídek, A., Bapst, V., Kohli, P., Jaderberg, M., Hassabis, D. and Jumper, J. 2024. Accurate structure prediction of biomolecular interactions with AlphaFold 3. Nature 630(8016), 493–500.

Acosta-González, A., Rosselló-Móra, R. and Marqués, S. 2013. Characterization of the anaerobic microbial community in oil-polluted subtidal sediments: aromatic biodegradation potential after the Prestige oil spill. Environmental Microbiology 15(1), 77–92.

Al-Daghistani, H.I., Zein, S. and Abbas, M.A. 2024. Microbial communities in the Dead Sea and their potential biotechnological applications. Communicative & Integrative Biology 17(1), 2369782.

Aramaki, T., Blanc-Mathieu, R., Endo, H., Ohkubo, K., Kanehisa, M., Goto, S. and Ogata, H. 2020. KofamKOALA: KEGG Ortholog assignment based on profile HMM and adaptive score threshold. Bioinformatics 36(7), 2251–2252.

Aroney, S., Newell, R., Nissen, J., Camargo, A., Tyson, G. and Woodcroft, B. 2025. CoverM: read alignment statistics for metagenomics. Bioinformatics 41(4), btaf147.

Bankevich, A., Nurk, S., Antipov, D., Gurevich, A., Dvorkin, M., Kulikov, A., Lesin, V., Nikolenko, S., Pham, S., Prjibelski, A., Pyshkin, A., Sirotkin, A., Vyahhi, N., Tesler, G., Alekseyev, M. and Pevzner, P. 2012. SPAdes: a new genome assembly algorithm and its applications to single-cell sequencing. Journal of Computational Biology 19(5), 455–477.

Bhattarai, B., Bhattacharjee, A., Coutinho, F. and Goel, R. 2021. Viruses and their interactions with bacteria and archaea of hypersaline Great Salt Lake. Frontiers in Microbiology 12, 701414.

Bolyen, E., Rideout, J., Dillon, M., Bokulich, N., Abnet, C., Al-Ghalith, G., Alexander, H., Alm, E., Arumugam, M., Asnicar, F., Bai, Y., Bisanz, J., Bittinger, K., Brejnrod, A., Brislawn, C., Brown, C., Callahan, B., Caraballo-Rodríguez, A., Chase, J., Cope, E., Da Silva, R., Diener, C., Dorrestein, P., Douglas, G., Durall, D., Duvallet, C., Edwardson, C., Ernst, M., Estaki, M., Fouquier, J., Gauglitz, J., Gibbons, S., Gibson, D., Gonzalez, A., Gorlick, K., Guo, J., Hillmann, B., Holmes, S., Holste, H., Huttenhower, C., Huttley, G., Janssen, S., Jarmusch, A., Jiang, L., Kaehler, B., Bin Kang, K., Keefe, C., Keim, P., Kelley, S., Knights, D., Koester, I., Kosciolek, T., Kreps, J., Langille, M., Lee, J., Ley, R., Liu, Y., Loftfield, E., Lozupone, C., Maher, M., Marotz, C., Martin, B., McDonald, D., McIver, L., Melnik, A., Metcalf, J., Morgan, S., Morton, J., Naimey, A., Navas Molina, J., Nothias, L., Orchanian, S., Pearson, T., Peoples, S., Petras, D., Preuss, M., Pruesse, E., Rasmussen, L., Rivers, A., Robeson, M., Rosenthal, P., Segata, N., Shaffer, M., Shiffer, A., Sinha, R., Song, S., Spear, J., Swafford, A., Thompson, L., Torres, P., Trinh, P., Tripathi, A., Turnbaugh, P., Ul-Hasan, S., vander Hooft, J., Vargas, F., Vázquez-Baeza, Y., Vogtmann, E., von Hippel, M., Walters, W., Wan, Y., Wang, M., Warren, J., Weber, K., Williamson, C., Willis, A., Xu, Z., Zaneveld, J., Zhang, Y., Zhu, Q., Knight, R. and Caporaso, J. 2019. Reproducible, interactive, scalable and extensible microbiome data science using QIIME 2. Nature Biotechnology 37(8), 852–857.

Capella-Gutiérrez, S., Silla-Martínez, J. and Gabaldón, T. 2009. trimAl: a tool for automated alignment trimming in large-scale phylogenetic analyses. Bioinformatics 25(15), 1972–1973.

Carr, M. and Head, J. 2010. Geologic history of Mars. Earth and Planetary Science Letters 294(3-4), 185–203.

Casamayor, E., Triadó-Margarit, X. and Castañeda, C. 2013. Microbial biodiversity in saline shallow lakes of the Monegros Desert, Spain. Fems Microbiology Ecology 85(3), 503–518.

Chasapi, M., Kontis, N., Lehmann, R., Tasneem, R., Patel, N., Khan, S., de Morentin, X., Chasapi, I., Aplakidou, E., Galaras, A., Aldakheel, L., Su, M., Baltoumas, F., Venkateswaran, K., Lagani, V., Gómez-Cabrero, D., Tegnér, J., Pavlopoulos, G. and Rosado, A. 2026. Decoding extremophiles: insights from bioinformatics, machine learning, and data-driven approaches. Briefings in Bioinformatics 27(3), bbag236.

Chaudhari, N., Overholt, W., Figueroa-Gonzalez, P., Taubert, M., Bornemann, T., Probst, A., Hölzer, M., Marz, M. and Küsel, K. 2021. The economical lifestyle of CPR bacteria in groundwater allows little preference for environmental drivers. Environmental Microbiome 16(1), 24.

Chaumeil, P., Mussig, A., Hugenholtz, P. and Parks, D. 2020. GTDB-Tk: a toolkit to classify genomes with the Genome Taxonomy Database. Bioinformatics 36(6), 1925–1927.

Chen, Y., Leung, P., Wood, J., Bay, S., Hugenholtz, P., Kessler, A., Shelley, G., Waite, D., Franks, A., Cook, P. and Greening, C. 2021. Metabolic flexibility allows bacterial habitat generalists to become dominant in a frequently disturbed ecosystem. ISME J 15(10), 2986–3004.

Chen, Y., Shen, J., Liu, L., Sun, Y., Pan, Y. and Lin, W. 2022. Preservation of organic matter in aqueous deposits and soils across the Mars-analog qaidam basin, NW China: implications for biosignature detection on Mars. Journal of Geophysical Research-Planets 127(12), e2022JE007418.

Cheng, M.Y., Luo, S., Zhang, P., Xiong, G.Z., Chen, K., Jiang, C.Q., Yang, F.D., Huang, H.H., Yang, P.S., Liu, G.X., Zhang, Y.H., Ba, S., Yin, P., Xiong, J., Miao, W. and Ning, K. 2024. A genome and gene catalog of the aquatic microbiomes of the Tibetan Plateau. Nature Communications 15(1), 1438.

Chibani, C., Mahnert, A., Borrel, G., Almeida, A., Werner, A., Brugère, J., Gribaldo, S., Finn, R., Schmitz, R. and Moissl-Eichinger, C. 2022. A catalogue of 1,167 genomes from the human gut archaeome. Nature Microbiology 7(1), 48–61.

Corman, J., Poret-Peterson, A., Uchitel, A. and Elser, J. 2016. Interaction between lithification and resource availability in the microbialites of Rio Mesquites, Cuatro Cienegas, Mexico. Geobiology 14(2), 176–189.

Doherty, A., Jackson, S. and Weller, G. 2001. Identification of bacterial homologues of the Ku DNA repair proteins. FEBS Letters 500(3), 186–188.

Fambrini, M., Usai, G., Vangelisti, A., Mascagni, F. and Pugliesi, C. 2020. The plastic genome: The impact of transposable elements on gene functionality and genomic structural variations. Genesis 58(12), e23399.

Farias, M., Rasuk, M., Gallagher, K., Contreras, M., Kurth, D., Fernandez, A., Poiré, D., Novoa, F. and Visscher, P. 2017. Prokaryotic diversity and biogeochemical characteristics of benthic microbial ecosystems at La Brava, a hypersaline lake at Salar de Atacama, Chile. PLoS One 12(11), e0186867.

Fernandez, A., Rasuk, M., Visscher, P., Contreras, M., Novoa, F., Poire, D., Patterson, M., Ventosa, A. and Farias, M. 2016. Microbial diversity in sediment ecosystems (evaporites domes, microbial mats, and crusts) of hypersaline Laguna Tebenquiche, Salar de Atacama, Chile. Frontiers in Microbiology 7, 215580.

Fu, L., Niu, B., Zhu, Z., Wu, S. and Li, W. 2012. CD-HIT: accelerated for clustering the next-generation sequencing data. Bioinformatics 28(23), 3150–3152.

Gao, L., Fang, B., Yang, J., Lian, Z., Chen, Y., Mohamad, O., Xu, Q., Liu, Y., Wu, D., Yuan, Y., Abdugheni, R., Li, M., Wang, P., Ortúzar, M., Li, X., Huang, J., Liu, L., Jiang, H., Shu, W., Hedlund, B., Li, W. and Jiao, J. 2026. Microbial decomposer diversity and metabolic function during the decomposition of brine shrimp carcasses in a saline lake. Microbiome 14(1), 141.

Greening, C., Geier, R., Wang, C., Woods, L., Morales, S., McDonald, M., Rushton-Green, R., Morgan, X., Koike, S., Leahy, S., Kelly, W., Cann, I., Attwood, G., Cook, G. and Mackie, R. 2019. Diverse hydrogen production and consumption pathways influence methane production in ruminants. ISME J 13(10), 2617–2632.

Guillou, L., Bachar, D., Audic, S., Bass, D., Berney, C., Bittner, L., Boutte, C., Burgaud, G., de Vargas, C., Decelle, J., del Campo, J., Dolan, J., Dunthorn, M., Edvardsen, B., Holzmann, M., Kooistra, W., Lara, E., Le Bescot, N., Logares, R., Mahé, F., Massana, R., Montresor, M., Morard, R., Not, F., Pawlowski, J., Probert, I., Sauvadet, A., Siano, R., Stoeck, T., Vaulot, D., Zimmermann, P. and Christen, R. 2013. The Protist Ribosomal Reference database (PR2): a catalog of unicellular eukaryote small sub-unit rRNA sequences with curated taxonomy. Nucleic Acids Research 41(D1), D597–D604.

Gutiérrez-Preciado, A., Dede, B., Baker, B., Eme, L., Moreira, D. and López-García, P. 2024. Extremely acidic proteomes and metabolic flexibility in bacteria and highly diversified archaea thriving in geothermal chaotropic brines. Nature Ecology & Evolution 8(10), 1856–1869.

Gutiérrez-Preciado, A., Saghaï, A., Moreira, D., Zivanovic, Y., Deschamps, P. and López-García, P. 2018. Functional shifts in microbial mats recapitulate early Earth metabolic transitions. Nature Ecology & Evolution 2(11), 1700–1708.

Hamm, J., Dombrowski, N., Valentin-Alvarado, L., Greening, C., Williams, T. and Spang, A. 2026. New lineages provide insights into the convergent evolution of extreme salt adaptation within symbiotic Archaea. Molecular Biology and Evolution 43(5), msag091.

He, Y.H., Baltar, F. and Wang, Y. 2025. Seasonal variability in community structure and metabolism of active deep-sea microorganisms. ISME J 19(1), wraf214.

Heijs, S., Laverman, A., Forney, L., Hardoim, P. and van Elsas, J. 2008. Comparison of deep-sea sediment microbial communities in the Eastern Mediterranean. FEMS Microbiology Ecology 64(3), 362–377.

Herrmann, M., Wegner, C., Taubert, M., Geesink, P., Lehmann, K., Yan, L., Lehmann, R., Totsche, K. and Küsel, K. 2019. Predominance of Cand. Patescibacteria in groundwater is caused by their preferential mobilization from soils and flourishing under oligotrophic conditions. Frontiers in Microbiology 10, 1407.

Hoffmann, T., Warmbold, B., Smits, S., Tschapek, B., Ronzheimer, S., Bashir, A., Chen, C., Rolbetzki, A., Pittelkow, M., Jebbar, M., Seubert, A., Schmitt, L. and Bremer, E. 2018. Arsenobetaine: an ecophysiologically important organoarsenical confers cytoprotection against osmotic stress and growth temperature extremes. Environmental Microbiology 20(1), 305–323.

Huang, W., Liu, Y., Zhang, X., Zhang, C., Zou, D., Zheng, S., Xu, W., Luo, Z., Liu, F. and Li, M. 2021. Comparative genomic analysis reveals metabolic flexibility of Woesearchaeota. Nature Communications 12(1), 5281.

Huang, Y., Gilna, P. and Li, W. 2009. Identification of ribosomal RNA genes in metagenomic fragments. Bioinformatics 25(10), 1338–1340.

Hyatt, D., Chen, G., LoCascio, P., Land, M., Larimer, F. and Hauser, L. 2010. Prodigal: prokaryotic gene recognition and translation initiation site identification. BMC Bioinformatics 11, 119.

Jenney, F.J., Verhagen, M., Cui, X. and Adams, M. 1999. Anaerobic microbes: Oxygen detoxification without superoxide dismutase. Science 286(5438), 306–309.

Jiang, P., Liang, Z., Kovacevic, V., Shi, J., Milicevic, N., Wang, F., Liu, L., Liu, Y., Jiang, Y., Han, M., Lin, X., Petronic, C., Stanojevic, N., Wang, L., Wang, S., Cheng, H., Li, J., Chen, R., Zhang, Y., Li, Y., Li, J., Fang, X., Yue, Z., Xue, C., Yin, P. and Chen, H. 2026. The Extreme Environment Microbiome Catalog (EEMC): a global resource for microbial diversity and antimicrobial discovery. Nature Communications 17(1), 4791.

Jordaan, K., Lappan, R., Dong, X., Aitkenhead, I., Bay, S., Chiri, E., Wieler, N., Meredith, L., Cowan, D., Chown, S. and Greening, C. 2020. Hydrogen-oxidizing bacteria are abundant in desert soils and strongly stimulated by hydration. Msystems 5(6), e01131–01120.

Junier, T. and Zdobnov, E. 2010. The Newick utilities: high-throughput phylogenetic tree processing in the Unix shell. Bioinformatics 26(13), 1669–1670.

Katoh, K. and Standley, D. 2013. MAFFT multiple sequence alignment software version 7: improvements in performance and usability. Molecular Biology and Evolution 30(4), 772–780.

Klimek, D., Herold, M. and Calusinska, M. 2024. Comparative genomic analysis of Planctomycetota potential for polysaccharide degradation identifies biotechnologically relevant microbes. BMC Genomics 25(1), 523.

Kong, F.J., Zheng, M.P., Hu, B., Wang, A.A., Ma, N. and Sobron, P. 2018. Dalangtan saline playa in a hyperarid region on Tibet Plateau: I. Evolution and environments. Astrobiology 18(10), 1243–1253.

Kurth, D., Amadio, A., Ordoñez, O., Albarracín, V., Gärtner, W. and Farías, M. 2017. Arsenic metabolism in high altitude modern stromatolites revealed by metagenomic analysis. Scientific Reports 7, 1024.

Kurth, D., Elias, D., Rasuk, M., Contreras, M. and Farías, M. 2021. Carbon fixation and rhodopsin systems in microbial mats from hypersaline lakes Brava and Tebenquiche, Salar de Atacama, Chile. Plos One 16(2), e0246656.

Lenferink, W., van Alen, T., Jetten, M., op den Camp, H., van Kessel, M. and Luecker, S. 2024. Genomic analysis of the class Phycisphaerae reveals a versatile group of complex carbon-degrading bacteria. Antonie Van Leeuwenhoek International Journal of General and Molecular Microbiology 117(1), 104.

Li-Hau, F., Nakagawa, M., Kakegawa, T., Ward, L., Ueno, Y. and McGlynn, S. 2025. Metabolic potential and microbial diversity of late Archean to early Proterozoic ocean analog hot springs of Japan. Microbes and Environments 40(3), ME24067.

Li, D., Luo, R., Liu, C., Leung, C., Ting, H., Sadakane, K., Yamashita, H. and Lam, T. 2016. MEGAHIT v1.0: A fast and scalable metagenome assembler driven by advanced methodologies and community practices. Methods 102, 3–11.

Li, J., Liu, H., Meng, X., Duan, D., Lu, H., Zhang, J., Zhang, F., Elsworth, D., Cardenas, B., Manga, M., Zhou, B. and Fang, G. 2025a. Ancient ocean coastal deposits imaged on Mars. Proceedings of the National Academy of Sciences of the United States of America 122(9), e2422213122.

Li, Z., He, Y., Zhang, H., Qian, H. and Wang, Y. 2024. Biotransformations of arsenic in marine sediments across marginal slope to hadal zone. Journal of Hazardous Materials 480, 135955.

Li, Z., Wei, T., He, L., Qian, H., Zhu, Y. and Wang, Y. 2025b. Genomic potential for mercury biotransformation in marine sediments across marginal slope to hadal zone. Nature Communications 16(1), 8655.

Liu, L., Liu, H.Y., Zhang, W.S., Chen, Y., Shen, J.X., Li, Y.L., Pan, Y.X. and Lin, W. 2022. Microbial diversity and adaptive strategies in the Mars-like Qaidam Basin, North Tibetan Plateau, China. Environmental Microbiology Reports 14(6), 873–885.

Liu, L., Wang, Z., Zhang, W.S. and Lin, W. 2025. Recovery of 1,773 microbial genomes and 2,060 viral genomes from the Mars-analog Qaidam Basin desert. Scientific Data 12(1), 1795.

Magnuson, E., Altshuler, I., Freyria, N., Leveille, R. and Whyte, L. 2023. Sulfur-cycling chemolithoautotrophic microbial community dominates a cold, anoxic, hypersaline Arctic spring. Microbiome 11(1), 203.

Marin, J., Battistuzzi, F., Brown, A. and Hedges, S. 2017. The timetree of prokaryotes: new insights into their evolution and speciation. Molecular Biology and Evolution 34(2), 437–446.

Mason-Jones, K., Breidenbach, A., Dyckmans, J., Banfield, C. and Dippold, M. 2023. Intracellular carbon storage by microorganisms is an overlooked pathway of biomass growth. Nature Communications 14(1), 2240.

McCollom, T., Klein, F. and Ramba, M. 2022. Hydrogen generation from serpentinization of iron-rich olivine on Mars, icy moons, and other planetary bodies. Icarus 372, 114754.

McGonigle, J., Bernau, J., Bowen, B. and Brazelton, W. 2019. Robust archaeal and bacterial communities inhabit shallow subsurface sediments of the Bonneville Salt Flats. MSphere 4(4), e00378–00319.

Meaden, S., Biswas, A., Arkhipova, K., Morales, S., Dutilh, B., Westra, E. and Fineran, P. 2022. High viral abundance and low diversity are associated with increased CRISPR-Cas prevalence across microbial ecosystems. Current Biology 32(1), 220–227.

Messner, K., Pereira, C., Kyndt, J., Palmer, M. and Yurkov, V. 2026. Apolloniradiicaulis salifontis gen. nov., sp. nov., a New Prosthecate Aerobic Anoxygenic Phototroph Isolated from Lake Winnipegosis Region Salt Springs. Microorganisms 14(3), 525.

Miao, Y., Fang, X., Sun, J., Xiao, W., Yang, Y., Wang, X., Farnsworth, A., Huang, K., Ren, Y., Wu, F., Qiao, Q., Zhang, W., Meng, Q., Yan, X., Zheng, Z., Song, C. and Utescher, T. 2022. A new biologic paleoaltimetry indicating Late Miocene rapid uplift of northern Tibet Plateau. Science 378(6624), 1074–1078.

Minh, B., Schmidt, H., Chernomor, O., Schrempf, D., Woodhams, M., von Haeseler, A. and Lanfear, R. 2020. IQ-TREE 2: new models and efficient methods for phylogenetic inference in the genomic era. Molecular Biology and Evolution 37(5), 1530–1534.

Muigano, M., Mauti, G., Anami, S. and Onguso, J. 2025. Advances and challenges in polyhydroxyalkanoates (PHA) production using Halomonas species: A review. International Journal of Biological Macromolecules 309, 142850.

Nakajima, M., Nakai, R., Hirakata, Y., Kubota, K., Satoh, H., Nobu, M., Narihiro, T. and Kuroda, K. 2025. Minisyncoccus archaeiphilus gen. nov., sp. nov., a mesophilic, obligate parasitic bacterium and proposal of Minisyncoccaceae fam. nov., Minisyncoccales ord. nov., Minisyncoccia class. nov. and Minisyncoccota phyl. nov. formerly referred to as Candidatus Patescibacteria or candidate phyla radiation. International Journal of Systematic and Evolutionary Microbiology 75(2), 006668.

Niedzwiedzki, D., Dilbeck, P., Tang, Q., Mothersole, D., Martin, E., Bocian, D., Holten, D. and Hunter, C. 2015. Functional characteristics of spirilloxanthin and keto-bearing analogues in light-harvesting LH2 complexes from Rhodobacter sphaeroides with a genetically modified carotenoid synthesis pathway. Biochimica Et Biophysica Acta-Bioenergetics 1847(6-7), 640–655.

Obruca, S., Dvorák, P., Sedlácek, P., Koller, M., Sedlár, K., Pernicová, I. and Safránek, D. 2022. Polyhydroxyalkanoates synthesis by halophiles and thermophiles: towards sustainable production of microbial bioplastics. Biotechnology Advances 58, 107906.

Olm, M.R., Brown, C.T., Brooks, B. and Banfield, J.F. 2017. dRep: a tool for fast and accurate genomic comparisons that enables improved genome recovery from metagenomes through de-replication. ISME J 11(12), 2864–2868.

Orakov, A., Fullam, A., Coelho, L., Khedkar, S., Szklarczyk, D., Mende, D., Schmidt, T. and Bork, P. 2021. GUNC: detection of chimerism and contamination in prokaryotic genomes. Genome Biology 22(1), 178.

Oremland, R. and Stolz, J. 2003. The ecology of arsenic. Science 300(5621), 939–944.

Parks, D., Chuvochina, M., Rinke, C., Mussig, A., Chaumeil, P. and Hugenholtz, P. 2022. GTDB: an ongoing census of bacterial and archaeal diversity through a phylogenetically consistent, rank normalized and complete genome-based taxonomy. Nucleic Acids Research 50(D1), D785–D794.

Parks, D., Imelfort, M., Skennerton, C., Hugenholtz, P. and Tyson, G. 2015. CheckM: assessing the quality of microbial genomes recovered from isolates, single cells, and metagenomes. Genome Research 25(7), 1043–1055.

Pérez-Hernández, V., Hernández-Guzmán, M., Luna-Guido, M., Navarro-Noya, Y., Romero-Tepal, E. and Dendooven, L. 2021. Bacterial communities in alkaline saline soils amended with young maize plants or its (hemi) cellulose fraction. Microorganisms 9(6), 1297.

Quast, C., Pruesse, E., Yilmaz, P., Gerken, J., Schweer, T., Yarza, P., Peplies, J. and Glöckner, F. 2013. The SILVA ribosomal RNA gene database project: improved data processing and web-based tools. Nucleic Acids Research 41(D1), D590–D596.

Ramel, F., Amrani, A., Pieulle, L., Lamrabet, O., Voordouw, G., Seddiki, N., Brèthes, D., Company, M., Dolla, A. and Brasseur, G. 2013. Membrane-bound oxygen reductases of the anaerobic sulfate-reducing Desulfovibrio vulgaris Hildenborough: roles in oxygen defence and electron link with periplasmic hydrogen oxidation. Microbiology-Sgm 159, 2663–2673.

Rascovan, N., Maldonado, J., Vazquez, M. and Farías, M. 2016. Metagenomic study of red biofilms from Diamante Lake reveals ancient arsenic bioenergetics in haloarchaea. ISME J 10(2), 299–309.

Rauschenbach, I., Bini, E., Häggblom, M. and Yee, N. 2012. Physiological response of Desulfurispirillum indicum S5 to arsenate and nitrate as terminal electron acceptors. FEMS Microbiology Ecology 81(1), 156–162.

Roberts, A., Adler, B., Cress, B., Doudna, J. and Barrangou, R. 2025. Phage-based delivery of CRISPR-associated transposases for targeted bacterial editing. Proceedings of the National Academy of Sciences of the United States of America 122(30), e2504853122.

Rodrigues, J., Schmidt, T., Tackmann, J. and von Mering, C. 2017. MAPseq: highly efficient k-mer search with confidence estimates, for rRNA sequence analysis. Bioinformatics 33(23), 3808–3810.

Rodrigues, J., Tackmann, J., Malfertheiner, L., Patsch, D., Perez-Molphe-Montoya, E., Näpflin, N., Gaio, D., Rot, G., Danaila, M., Peluso, M., Dmitrijeva, M., Schmidt, T. and von Mering, C. 2026. The MicrobeAtlas database: Global trends and insights into Earth’s microbial ecosystems. Cell 189(7), 2092–2107.

Saghaï, A., Gutierrez-Preciado, A., Deschamps, P., Moreira, D., Bertolino, P., Ragon, M. and López-Garcïa, P. 2017. Unveiling microbial interactions in stratified mat communities from a warm saline shallow pond. Environmental Microbiology 19(6), 2405–2421.

Sancho-Tomás, M., Somogyi, A., Medjoubi, K., Bergamaschi, A., Visscher, P., Van Driessche, A., Gérard, E., Farias, M., Contreras, M. and Philippot, P. 2018. Distribution, redox state and (bio)geochemical implications of arsenic in present day microbialites of Laguna Brava, Salar de Atacama. Chemical Geology 490, 13–21.

Schiwitza, S., Gutsche, L., Freches, E., Arndt, H. and Nitsche, F. 2021. Extended divergence estimates and species descriptions of new craspedid choanoflagellates from the Atacama Desert, Northern Chile. European Journal of Protistology 79, 125798.

Slouf, V., Chábera, P., Olsen, J., Martin, E., Qian, P., Hunter, C. and Polívka, T. 2012. Photoprotection in a purple phototrophic bacterium mediated by oxygen-dependent alteration of carotenoid excited-state properties. Proceedings of the National Academy of Sciences of the United States of America 109(22), 8570–8575.

Song, W., Wemheuer, B., Zhang, S., Steensen, K. and Thomas, T. 2019. MetaCHIP: community-level horizontal gene transfer identification through the combination of best-match and phylogenetic approaches. Microbiome 7, 36.

Sorokin, D., Tourova, T., Panteleeva, A. and Muyzer, G. 2012. Desulfonatronobacter acidivorans gen. nov., sp. nov. and Desulfobulbus alkaliphilus sp. nov., haloalkaliphilic heterotrophic sulfate-reducing bacteria from soda lakes. International Journal of Systematic and Evolutionary Microbiology 62, 2107–2113.

Stamatakis, A. 2014. RAxML version 8: a tool for phylogenetic analysis and post-analysis of large phylogenies. Bioinformatics 30(9), 1312–1313.

Starr, S. and Muscatello, A. 2020. Mars in situ resource utilization: a review. Planetary and Space Science 182, 104824.

Steinegger, M. and Söding, J. 2017. MMseqs2 enables sensitive protein sequence searching for the analysis of massive data sets. Nature Biotechnology 35(11), 1026–1028.

Stober, I., Zhong, J. and Bucher, K. 2023. From freshwater inflows to salt lakes and salt deposits in the Qaidam Basin, W China. Swiss Journal of Geosciences 116(1), 5.

Sun, Y.Y., Liang, Y., Liu, H., Liu, J., Ji, J.L., Ke, X., Liu, X.B., He, Y.X., Wang, H.Y., Zhang, B., Zhang, Y.S., Zhuang, G.S., Pei, J.L., Li, Y.X., Quan, C., Li, J.X., Aitchison, J.C., Liu, W.G. and Liu, Z.H. 2023. Mid-Miocene sea level altitude of the Qaidam Basin, northern Tibetan Plateau. Communications Earth & Environment 4(1), 3.

Sunagawa, S., Mende, D., Zeller, G., Izquierdo-Carrasco, F., Berger, S., Kultima, J., Coelho, L., Arumugam, M., Tap, J., Nielsen, H., Rasmussen, S., Brunak, S., Pedersen, O., Guarner, F., de Vos, W., Wang, J., Li, J., Dore, J., Ehrlich, S., Stamatakis, A. and Bork, P. 2013. Metagenomic species profiling using universal phylogenetic marker genes. Nature Methods 10(12), 1196–1199.

Szydlowski, L., Bulbul, A., Simpson, A., Kaya, D., Singh, N., Sezerman, U., Labaj, P., Kosciolek, T. and Venkateswaran, K. 2024. Adaptation to space conditions of novel bacterial species isolated from the International Space Station revealed by functional gene annotations and comparative genome analysis. Microbiome 12(1), 190.

Tarnas, J., Mustard, J., Lollar, B., Bramble, M., Cannon, K., Palumbo, A. and Plesa, A. 2018. Radiolytic H2 production on Noachian Mars: Implications for habitability and atmospheric warming. Earth and Planetary Science Letters 502, 133–145.

Uritskiy, G., DiRuggiero, J. and Taylor, J. 2018. MetaWRAP-a flexible pipeline for genome-resolved metagenomic data analysis. Microbiome 6, 158.

Vavourakis, C., Andrei, A., Mehrshad, M., Ghai, R., Sorokin, D. and Muyzer, G. 2018. A metagenomics roadmap to the uncultured genome diversity in hypersaline soda lake sediments. Microbiome 6, 168.

Veloso, M., Waldisperg, A., Arros, P., Berríos-Pastén, C., Acosta, J., Colque, H., Varas, M., Allende, M., Orellana, L. and Marcoleta, A. 2023. Diversity, taxonomic novelty, and encoded functions of Salar de Ascotán microbiota, as revealed by metagenome-assembled genomes. Microorganisms 11(11), 2819.

Wang, H.R., Zhang, L.X., Tian, C., Fan, S., Zheng, D.C., Song, Y.H., Gao, P. and Li, D.P. 2024. Effects of nitrogen supply on hydrogen-oxidizing bacterial enrichment to produce microbial protein: Comparing nitrogen fixation and ammonium assimilation. Bioresource Technology 394, 130199.

Wang, J., Wang, F., Chu, L., Wang, H., Zhong, Z., Liu, Z., Gao, J. and Duan, H. 2014. High genetic diversity and novelty in eukaryotic plankton assemblages inhabiting saline lakes in the Qaidam Basin. PLoS One 9(11), e112812.

Wang, Y., Li, F.C., Zhao, J., Chen, H., Jiang, P. and Tang, X. 2019. Microbial community diversity and vertical distribution in a columnar sediment of Maluku Strait. Journal of Atmospheric Science Research 2(2), 51–58.

Weller, G., Kysela, B., Roy, R., Tonkin, L., Scanlan, E., Della, M., Devine, S., Day, J., Wilkinson, A., di Fagagna, F., Devine, K., Bowater, R., Jeggo, P., Jackson, S. and Doherty, A. 2002. Identification of a DNA nonhomologous end-joining complex in bacteria. Science 297(5587), 1686–1689.

West, P., Probst, A., Grigoriev, I., Thomas, B. and Banfield, J. 2018. Genome-reconstruction for eukaryotes from complex natural microbial communities. Genome Research 28(4), 569–580.

Williams, A., Eigenbrode, J., Millan, M., Williams, R., Mcintosh, O., Teinturier, S., Roach, J., Malespin, C., McAdam, A., Mahaffy, P., Bryk, A., Buch, A., Boulesteix, D., Chou, L., Dworkin, J., Fox, V., Franz, H., Freissinet, C., Glavin, D., House, C., Johnson, S., Lewis, J., Mojarro, A., Navarro-Gonzalez, R., Pozarycki, C., Steele, A., Summons, R., Szopa, C., Thorpe, M. and Vasavada, A. 2026. Diverse organic molecules on Mars revealed by the first SAM TMAH experiment. Nature Communications 17(1), 2748.

Wu, Y., Liao, L., Wang, C., Ma, W., Meng, F., Wu, M. and Xu, X. 2013. A comparison of microbial communities in deep-sea polymetallic nodules and the surrounding sediments in the Pacific Ocean. Deep-Sea Research Part I-Oceanographic Research Papers 79, 40–49.

Yu, T., Luo, Y., Tan, X., Zhao, D., Bi, X., Li, C., Zheng, Y., Xiang, H. and Hu, S. 2024. Global marine cold seep metagenomes reveal diversity of taxonomy, metabolic function, and natural products. Genomics Proteomics & Bioinformatics 22(2), qzad006.

Zhang, H., Wei, T., Zhang, J., Li, Q., Fu, L., He, L. and Wang, Y. 2024. Microbial mediation of cryptic methane cycle in semiclosed marine water column. The Innovation Geoscience 2(3), 100082.

Zhang, R., Wang, C., Wang, X., Mu, D. and Du, Z. 2019a. Jannaschia formosa sp. nov., isolated from marine saltern sediment. International Journal of Systematic and Evolutionary Microbiology 69(7), 2037–2042.

Zhang, W., Chen, C., Yuan, Y., Su, D., Ding, L., Epstein, S. and He, S. 2019b. Pararhodobacter marinus sp. nov., a bacterium isolated from marine sediment in the East China Sea and emended description of the genus Pararhodobacter. International Journal of Systematic and Evolutionary Microbiology 69(10), 3293–3298.

Zhang, Z.F., Huang, J.E., Phurbu, D., Qu, Z.S., Liu, F. and Cai, L. 2025. A deep metagenomic atlas of Qinghai-Xizang Plateau lakes reveals their microbial diversity and salinity adaptation mechanisms. Cell Reports 44(11).

Zhaxybayeva, O. and Nesbo, C. 2026. Impact of horizontal gene transfer on adaptations to extreme environments. Journal of Molecular Biology 438(4), 169403.

Zheng, J., Hu, B., Zhang, X., Ge, Q., Yan, Y., Akresi, J., Piyush, V., Huang, L. and Yin, Y. 2023. dbCAN-seq update: CAZyme gene clusters and substrates in microbiomes. Nucleic Acids Research 51(D1), D557–D563.

Zhou, Y., Mara, P., Cui, G., Edgcomb, V. and Wang, Y. 2022. Microbiomes in the Challenger Deep slope and bottom-axis sediments. Nature Communications 13(1), 1515.

Zou, H., Shi, M., Zhang, T., Li, L., Li, L. and Xian, M. 2017. Natural and engineered polyhydroxyalkanoate (PHA) synthase: key enzyme in biopolyester production. Applied Microbiology and Biotechnology 101(20), 7417–7426.

