## Supplementary methods, results, tables, figures; files of main figures for "Microbial evolution, biogeochemical functions, and environmental adaptations in a desert saline lake on the Qinghai–Tibet Plateau": Supplemental information.docx

### **Supplementary methods and results**

#### **1. Detection of concentration of oxidized inorganic nitrogen species**

Nitrate- and nitrite-nitrogen concentrations in the filtered liquid samples, diluted 30-fold (considering the limited amount of filtered pore water), were below the detection limits of the colorimetric assay (Hach reagent 2107169) based on diazonium salts. Consistently, no discernible color development was observed by the WG-752N UV–visible spectrophotometry (INESA, Shanghai, China) after a 20-fold dilution and reaction with Griess reagent (Hakobyan et al., 2022), indicating extremely low levels of these oxidized inorganic nitrogen species (< 0.06 mg/L).

#### **2. Raw Illumina sequencing data processing**

Illumina paired-end shotgun metagenomic sequencing generated 41.0–44.1 Gb raw data per sample, corresponding to approximately 273–294 million reads per metagenome.

Raw reads were trimmed to remove adapters and then filtered using fastp (v.0.23.4) (Chen et al., 2018) with parameters “-f 1 -t 0 -F 1 -T 0 -w 24 -c -q 20 -u 20 -g -W 5 -3 -l 50”. Fastuniq was used to eliminate duplicated paired short reads with default parameters (Xu et al., 2012). Processed reads were aligned to the laboratory contamination control metagenome (He et al., 2025) using BWA, and alignment files were subsequently processed with Samtools and Bedtools to remove reads mapping to the control sequences. Reads that did not align to the control metagenome were retained as the final clean reads.

#### **3. Phylogenetic analysis of representative bacterial OTUs**

The “SILVA_138_SSURef_NR99_tax_silva” dataset was imported into QIIME2, and the V4 region was extracted. Taxonomic annotations for the reference sequences were assigned using the silva-138-99-tax.qza classifier (<https://data.qiime2.org/2022.8/common/silva-138-99-tax.qza>).

For Acidobacteriota Subgroup 26, a total of 38 reference sequences and 33 OTU sequences were selected to construct a maximum-likelihood phylogenetic tree, in which 28 OTUs clustered with “AY225650”. These OTUs were subsequently compared against the SILVA 138 database using BLASTn, and 17 OTUs with the highest likelihood of affiliation to Subgroup 26 were retained. These sequences, together with reference sequences showing > 97% identity in BLASTn results and the closest matches retrieved from the NCBI core_nt database, were used to reconstruct a refined maximum-likelihood phylogenetic tree (Fig. S4). In addition, the most abundant OTU in sample S3 (6e92436d4db1b61aaeda1ab8741ee007b7ad20b1; Supplementary Table S2) was selected to construct the maximum-likelihood phylogenetic tree with 12 closely related reference sequences (a part in Fig. 2H).

For the two OTUs assigned to *Flexithrix*, a maximum-likelihood phylogenetic tree was reconstructed using these OTUs together with 78 reference sequences affiliated with Flammeovirgaceae. The two OTUs clustered with JN428535. BLASTn searches against the NCBI core_nt database further indicated that MF995916, which is not included in the SILVA database (Supplementary Table S3), was more closely related to OTU 3ac9f48dcf8d6e4cee30cbeddb84e168a1732c89 than JN428535 was. Although no SILVA reference sequence exceeding 97% similarity was identified for this OTU, it was therefore not treated as a novel sequence in Fig. 2H. The two OTUs shared a 117-bp exact overlap, and the shorter OTU, 0f72228aa9d8e0759f0407f260079499fa65130f, was also most closely related to MF995916.

For the OTU assigned to *Candidatus* Falkowiibacteriota (32edb73b26b29e60beaf68da8634d7c41d08cc18), which accounted for 1.70% of the S4 community, a maximum-likelihood tree was reconstructed together with 82 reference sequences from this phylum. This OTU clustered with JF738100, HM481307, and HM481357. BLASTn searches against the core_nt database indicated that FM867974, derived from hydrothermal sediments and not represented in the SILVA reference set, was the closest sequence. In the pruned tree shown in Fig. 2H, JF738100 and FM867974 were retained.

For Phycisphaerales, 21 of 41 OTUs were further assigned to the Phycisphaeraceae Urania-1B-19 marine sediment group. Three of these OTUs each exceeded 1% relative abundance in S1. A maximum-likelihood tree was reconstructed using these OTUs together with 956 reference sequences from this order. OTU 1499986676230991738e51e6d3e97e5f25a35504 clustered with the deep-sea sequences KM454365 and KX172650, whereas OTUs 15588ae15124dd21e4c2886a03549dc7be62a33b and cb57cd80ab16a6da337a7cff95369ddadf3b1132 formed a separate clade with marine-sediment sequences AY627529, JQ580169, and JX227329. OTU 15588ae15124dd21e4c2886a03549dc7be62a33b was identified as novel; its closest core_nt match showed a maximum percent identity of 96.32% (E value, 3 × 10^-54^), and the reference sequences were almost exclusively derived from marine environments. For the longer OTU cb57cd80ab16a6da337a7cff95369ddadf3b1132, BLASTn searches against core_nt suggested that several sequences from the endolithic community of Mono Island (Puerto Rico), which are not included in the SILVA reference set, were the closest relatives; KT973342 was therefore retained in the simplified tree shown in Fig. 2H.

Fig. S5 provides a more detailed view of the phylogenetic relationships between representative high-abundance OTUs from different bacteria phyla and their closest reference sequences.

#### **4. Phylogenetic analysis of *Candidatus* Woesearchaeota MAGs’ protein sequences**

For *dfx* (K05919; superoxide reductase/desulfoferrodoxin), the 126-aa protein sequence encoded by S4-bin11 was first queried against the NCBI NR database using BLASTp, and the top 100 homologous sequences were retrieved, most of which were bacterial. Additional BLASTp searches restricted to Archaea and *Candidatus* Woesearchaeota were then performed, yielding 100 homologous sequences from each search. After dereplication, a total of 274 reference sequences, mostly 120-130 aa in length, were retained and used together with the S4-bin11 sequence to reconstruct a maximum-likelihood phylogeny using IQ-TREE v.2.2.0 with the parameters “-m MFP -B 1000 -alrt 1000 -T AUTO --bnni --mem 100G”. S4-bin11 *dfx* tended to cluster with 6 sequences from Methanobacteriati (clade bootstrap: 88.2%).

For *gapN* (K00131; glyceraldehyde-3-phosphate dehydrogenase), the 545-aa protein sequence encoded by S4-bin3 was first queried against the NCBI NR database using BLASTp to retrieve the top 100 homologous sequences, followed by an additional BLASTp search restricted to Archaea to obtain another 100 sequences. After dereplication, a total of 199 reference sequences, annotated as aldehyde dehydrogenase family proteins or NADP-dependent glyceraldehyde-3-phosphate dehydrogenases, were retained and used together with the S4-bin3 and S4-bin11 sequences to reconstruct a maximum-likelihood phylogeny using IQ-TREE.

Unrooted topology tests did not provide statistically robust support for a closer affinity of the S4-bin3 and S4-bin11 *gapN* sequences to archaeal homologs over bacterial homologs. Both archaeal-affinity and bacterial-affinity local constrained topologies were compatible with the data under the tested substitution models (IQ-TREE AU topology tests; constrained ML candidate trees forcing S4-bin3/S4-bin11 to group with the nearest 2, 3, 5, or 10 archaeal or bacterial homologs; tests repeated under Q.yeast+I+I+R6 selected by ModelFinder and under LG+F+R6 as a model-sensitivity analysis). Therefore, the phylogenetic placement of the two MAG-derived *gapN* sequences should be regarded as unresolved rather than confidently archaeal-affiliated or bacterial-affiliated.

#### **5. Phylogenetic analysis of *aoxB* sequences (AioA protein sequences)**

Contig-derived sequences annotated as *aoxB* by KofamScan were combined with 214 reference sequences (see Supplementary Table S6) to reconstruct a maximum-likelihood phylogeny, in order to infer the potential taxonomic origin of sequences, especially for those not assigned to MAGs. Phylogenetic analysis was performed using RAxML (v.8.2.12) under the PROTGAMMALG model. Maximum-likelihood trees were inferred with 1000 rapid bootstrap replicates using the -f a option, and bootstrap support values were mapped onto the best-scoring tree.

Based on the clustering patterns of the *aoxB* sequences from the contigs with the reference sequences in the phylogenetic tree (Fig. S9A), together with their distribution among the MAGs, these sequences are likely derived from Alphaproteobacteria, Gammaproteobacteria, Deinococci, and Desulfuromonadia. BLASTp searches against the NR database indicated that the sequence most closely related to the Deinococci *aoxB* was an *aoxB* from a Trueperaceae bacterium (MDZ7708468.1), which was recovered from Salar de Ascotán, a high-altitude arsenic-rich salt flat exposed to intense ultraviolet radiation in the Atacama Desert, Chile (Veloso et al., 2023), suggesting a broadly similar environmental context to Eboliang Hu saline lake samples.

The *aoxB* sequence in S2-bin12 matched only five Thermodesulfobacteriota *aoxB* sequences (BLASTp against NR database). Sequences with > 64% identity were almost exclusively from Gammaproteobacteria or Betaproteobacteria, whereas the closest hit was IAA34024.1 from a Geothermobacteraceae MAG, with 81.65% identity. The phylogenetic tree constructed from these six sequences and four representative sequences selected from the Beta/Gamma-proteobacterial clade containing the S2-bin12 *aoxB* in Fig. S9A (one being WP_109675809.1 at the base of this clade and the other three being *aoxB* sequences annotated from contigs recovered in this study) showed that IAA34024.1 and OGU17269.1, both derived from subsurface samples, clustered with Proteobacterial sequences, whereas the Thermodesulfobacteriota sequences outside this clade originated from marine sediments (Fig. S9B).

#### **6. Characteristic metabolic patterns of species in communities**

##### 6.1 Genome-informed prediction of species life-history strategy tendencies

Functional traits were assigned to three life-history strategy categories—high growth yield (Y), resource acquisition (A), and stress tolerance (S)—following the previous conceptual frameworks (Cheng et al., 2024; Malik et al., 2020; Wood et al., 2018). For each strategy, non-redundant gene sets were constructed (see Supplementary Table S7) and combined with KofamScan annotations of each MAG to obtain total gene copy numbers.

For the acquisition (A) strategy, the total abundance of CAZy families was additionally incorporated (Sorouri et al., 2024). The gene copy numbers (and CAZy totals for A) were first standardized across genomes using Z-scores. For the A strategy, Z-scores of gene copy number and CAZy abundance were summed and re-standardized to derive a unified acquisition propensity score. Yield (Y) and stress-tolerance (S) propensity scores were derived from Z-score–standardized total gene copy numbers. Finally, each genome was assigned its dominant life-history strategy based on the highest standardized propensity score among Y, A, and S.

A total of 16 MAGs were included in the comparative analysis, comprising all 13 high-quality MAGs and three additional dominant medium-quality MAGs (*Marinobacter guineae* S1-bin4, 88.75% completeness; *Limnobacter profundi* S3-bin9, 81.93%; Myxococcota bacterium S4-bin21, 88.38%). This dataset encompassed seven of the ten dominant MAGs identified in the community (Fig. S7A). The remaining three dominant MAGs, all medium-quality Thermodesulfobacteriota genomes, were excluded because this phylum was already represented by two high-quality genomes (S1-bin1 and S4-bin12) in the analysis. These genomes were used to infer life history strategy tendencies of the corresponding taxa.

Across 15 genomes with completeness > 88%, the total number of annotated CAZys showed a significant positive correlation with genome size (Fig. S10A; Spearman R² = 0.27, *p* = 0.026). As shown in Fig. S10B, the functional gene composition of dominant MAGs suggested that the corresponding taxa predominantly favored Y- or A-strategies, whereas only *M. guineae* exhibited a tendency toward an S-strategy.

Genes involved in osmotic pressure regulation (shown in Fig. 5B), which do not overlap with gene sets representing the three life history strategies, were subsequently incorporated into the analysis. Based on the combined profiles of these four gene categories, hierarchical clustering was performed for the 16 genomes. The two Cyanobacteriota genomes clustered most closely with each other, reflecting their relative enrichment in S-strategy-associated genes and a marked depletion of osmotic pressure regulation transporters. In contrast, genome pairs affiliated with the same orders—Desulfuromonadales (S4-bin12 and S1-bin1) and Balneolales (S4-bin30 and S4-bin27)—displayed clear functional divergence, despite clustering within a group characterized by a higher abundance of osmotic pressure regulation genes (Fig. S10C).

##### 6.2 Genomic basis underlying ecological dominance and adaptation of dominant MAGs

Based on the relative enrichment of dominant MAGs across the four functional gene categories (genes related to high growth yield, resource acquisition, stress tolerance and osmotic pressure regulation) (Supplementary methods and results 6.1, Fig. S10), together with their distinct metabolic features compared to their closest phylogenetic reference genomes, —particularly with respect to substrate utilization and element cycling—the ecological basis underlying their dominance can be inferred.

In addition to the strong capacity for coping with environmental stress reflected in Fig. S10, the *M. guineae* genome S1-bin4 exhibits a highly flexible organic matter metabolism (Fig. S15). This genome uniquely encodes phenol 2-monooxygenase, enabling the conversion of benzene to phenol and further to catechol, as well as the subsequent degradation of catechol into pyruvate and acetaldehyde/acetyl-CoA; these metabolic capabilities were exclusive to this MAG among all recovered genomes. Moreover, it is the only Pseudomonadota genome in the dataset encoding acetyl-CoA synthetase, and it likely represents one of the key community members mediating the conversion of monosaccharides to pyruvate via the Entner–Doudoroff pathway (Fig. S15).

In samples S1 and S2, the Thermodesulfobacteriota MAG S1-bin1 represented the most abundant genome within that phylum. In addition to encoding nitrogenase and GS–GOGAT (*glnA* and *gltBD*), S1-bin1 harbors two distinct glutamate dehydrogenases (*gdhA*, K00262; *GDH2*, K15371) (Fig. S16). The presence of both ammonia-assimilatory routes suggests a competitive advantage in ammonium assimilation relative to other Thermodesulfobacteriota genomes. Compared with the reference genome GCA_002869665.1, S1-bin1 contains a duplicated *GDH2* copy and an expanded repertoire of *mcp* genes (10 copies versus 0 in the reference). S1-bin1 also scored highest for genes implicated in osmotic-pressure regulation among the 16 MAGs (Fig. S10).

The Flammeovirgaceae MAG S2-bin 24 was the only member among the five Bacteroidota genomes to encode an *amt* transporter together with a GLUD1_2-type glutamate dehydrogenase (K00261). Compared with 14 reference Flammeovirgaceae genomes (Fig. S24), this MAG additionally encoded a tryptophan 2-monooxygenase (implicated in L-tryptophan oxidation), a multicomponent Na⁺/H⁺ antiporter complex (mnhA–G), and a suite of CRISPR-associated proteins corresponding to Type I (K19114, K19115, K19116) and Type III (K07016, K09002, K19138, K19139, K19140) CRISPR–Cas systems (Fig. S17).

Compared with the other Myxococcota MAG in the dataset, phototrophic S4-bin21 uniquely encoded *amt* and two distinct glutamate dehydrogenases (absent from the closest reference genome GCA_036788295.1). S4-bin21 also encoded enzymes implicated in aromatic compound degradation, including phenylacetyl-CoA monooxygenase and the ring-cleavage-associated proteins PaaG and PaaZ, and it possessed the alkane 1-monooxygenase involved in aerobic alkane oxidation. Two distinct xanthine dehydrogenase systems (*xdhAB* and *yagRST*), not found in the reference genome, were detected and may oxidize xanthine to urate. In addition, S4-bin21 encodes complete Embden–Meyerhof, gluconeogenesis, and pentose phosphate pathways, as well as the glycogen degradation module (M00855) (Fig. S18); notably, the glycogen phosphorylase gene *glgP* is present in three copies in S4-bin21 compared with a single copy in the reference genome (Fig. S25).

The *Pseudooceanicola* MAG S3-bin8 potentially participates in the ring-cleavage degradation of 4-hydroxybenzoate, nitrate/nitrite reduction (harboring both *nasBD* and *nirK*), and thiosulfate oxidation. Although its genome size is smaller than that of the reference genome (GCF_000688295.1), S3-bin8 encodes additional functional capacities, including *napAB*, a complete SOX system (Fig. S19), and an increased copy number of the DNA mismatch repair protein MutS, which is present in two copies.

The *Limnobacter profundi* MAG S3-bin9 lacks a complete Entner–Doudoroff pathway but retains the capacity to catabolize phosphorylated gluconate to C3 intermediates (pyruvate and D-glyceraldehyde-3-phosphate) and encodes a complete TCA cycle. The genome also encodes pathways for β-oxidation of acetylated medium-chain fatty acids and for ammonia recovery from amino acid catabolism, as well as an assimilatory nitrite-to-ammonium reduction pathway and a SOX thiosulfate-oxidation system (Fig. S20). Relative to the larger reference genome (GCF_013004065.1), S3-bin9 shows copy-number expansions (one additional copy each) in four genes implicated in the conversion of 2-hydroxymuconate semialdehyde (a catechol ring-cleavage product) to pyruvate and acetyl-CoA (*dmpH*, *praC*, *bphI*, *bphJ*), and an additional copy of *dnaB* (replicative DNA helicase).

The *Pseudomonas alloputida* MAG S3-bin2 encoded an expanded repertoire of aromatic compound degradation genes and harbored three copies of the long-chain alkane monooxygenase. Compared with the larger reference genome, S3-bin2 carried an additional anthranilate 1,2-dioxygenase involved in anthranilate degradation, and exhibited copy-number expansions in three genes (*pcaB*, *pcaC*, *pcaD*) implicated in the conversion of 3,4-dihydroxybenzoate to 3-oxoadipate (Fig. S21). Four genes associated with the conversion of glycerone phosphate to pyruvate (*tpiA*, *gpmI*, *eno*, *pyk*) were likewise more abundant, and the genome also contained additional copies of the sulfate/thiosulfate transport genes *cysU* and *cysW*.

#### **7. Comparative genomic analysis of Phototrophicaceae MAGs**

The Phototrophicaceae MAGs recovered in this study (S2-bin4, S2-bin5, and S4-bin9) were compared with four genomes classified as the genus CAIRDJ01 in GTDB, which were derived from different water layers of the same freshwater lake. For phylogenetic analysis, the type species *Phototrophicus methaneseepsis* ZRK33 and seven additional genomes classified within the same family were included as references. Genome size and GC content suggest that these three newly recovered MAGs likely represent at least two distinct species, although they are more closely related to one another than to the other reference genomes (Fig. S22). Compared with the other genomes within this genus, the higher-completeness MAG S2-bin4 exhibited distinctive metabolic features, including the capacity to oxidize glucose and oxaloacetate and to reduce nitrite via both NrfAH and NirK, corresponding to assimilatory reduction to ammonium and dissimilatory reduction to nitric oxide, respectively (Fig. S13). BLASTp searches indicated that the closest NR homologs of its PufL protein, the photosynthetic reaction center L subunit, were from a Chloroflexota bacterium associated with Atlantic Ocean plastic particles (MEO0560538.1; 77.95% identity), whereas the closest homolog among Phototrophicales proteins was from an alkaline sulfidic hot spring (PJF26966.1; 65.30% identity). By contrast, *Phototrophicus methaneseepsis* ZRK33 has been reported to possess a photosystem II oxygen-evolving complex (Zheng et al., 2024), which is fundamentally different from the anoxygenic photosystem encoded by S2-bin4. Thus, the Phototrophicaceae MAGs recovered in this study represent rare genomes within this order that appear to mediate anoxygenic photosynthesis via the RC-LH1 complex.

#### **8. Comparative genomic analysis of Cyanophyceae MAGs**

The closest genome to S4-bin16 was GCA_004299065.1, whose taxonomic placement remains unresolved (GTDB assigns it to the genus *Sodalinema*, whereas NCBI classifies it as *Phormidium*). We therefore selected the two *Sodalinema* genomes available in the Genome database, eight *Phormidium* genomes (including three with ANI values > 83% relative to S4-bin16), and four *Geitlerinema* genomes from RefSeq as reference genomes for comparative genomic analysis. Similarly, the closest genome to S4-bin13 was GCF_000317615.1, whose taxonomic placement is also ambiguous (GTDB assigns it to *Halothece*, whereas NCBI classifies it as *Dactylococcopsis*). Accordingly, all six *Dactylococcopsis* genomes available in the Genome database and two *Halothece* genomes were included as reference genomes.

Phylogenetic analyses suggested that the brine-water environment may have strongly shaped the single-copy conserved protein profile of the S4-bin16 lineage (Fig. S23), consistent with its disputed taxonomic placement. Based on the functional gene profiles of the 24 genomes, the brine-water-associated clade tended to invest more broadly in osmotic-stress tolerance than reference genomes from non-saline environments, including sucrose synthesis (K00695, *SUS*) (Ehira et al., 2014; Santos-Merino et al., 2023), glucosylglycerol synthesis (K03692 + K05978, *E2.4.1.213* + *stpA*) (Luo et al., 2022), potassium uptake (K10716, *kch*) (Hagemann, 2011), and glycine betaine/proline transport (K02000–K02002; *proVWX*) (Rashid et al., 2023). In addition, genes related to iron uptake and arsenic/tellurium resistance (*feoABC*, *arsB*, *terD*, *arsC*), nitrogen fixation (*nif* cluster), and hydrogen sulfide oxidation (*sqr*) showed a similar distribution pattern. Compared with S4-bin13, S4-bin16 additionally encoded a NAD⁺-reducing hydrogenase (*hoxHY*) (Fig. S14). Relative to its two closest genomes (*Phormidium* sp. SL48-SHIP and *Sodalinema* sp. BLK5-1.bin.222), S4-bin16 also encoded an iron-siderophore transport system (*fepDGC*), subtype I-U factors of the Type I CRISPR-Cas system (*csb1*, *csb2*, *csx17*), and a Type I restriction-modification system (*hsdR*, *hsdS*, *hsdM*). S4-bin13 additionally encoded nitrogenase, nitric oxide reductase, and subtype I-U factors relative to *Dactylococcopsis salina* PCC 8305.

#### **9. Phylogenetic analysis of the novel Flammeovirgaceae MAG**

S2-bin24 represents a genome affiliated with an unclassified genus within the class Cytophagia and family Flammeovirgaceae (red value = 0.79985), characterized by a relatively small genome size and high GC content (Fig. S24A). To date, most Flammeovirgaceae genomes deposited in RefSeq have been recovered from marine samples, including seawater, sediments, endosymbionts of marine eukaryotes. In a maximum-likelihood phylogeny reconstructed from 14 genomes designated as “reference genomes” in the Genome database, together with S2-bin24 and using *Cytophaga aurantiaca* DSM 3654 (Cytophagaceae) as the outgroup, this MAG showed a relatively distant relationship to the other genomes. When the reference set was expanded to include 75 GTDB genomes from this family, the phylogenetic tree placed S2-bin24 and GCA_025056215.1 (from a hot spring, 2.8 Mbp with completeness of 99.45% and GC percentage of 36.94%) on a distinct branch (bootstrap: 99.1%), potentially representing a deeply diverged ancient lineage (Fig. S24B).

#### **10. Comparative genomic analysis of “*Candidatus* Kuafubacteriaceae JAHQVK01” MAGs**

S4-bin21 was classified by GTDB-Tk as a member of the genus “JAHQVK01”, which, according to the most recent GTDB release (R232), should be assigned to the newly proposed family *Candidatus* Kuafubacteriaceae, members of which all possess an anoxygenic photosystem II (Li et al., 2023). The maximum-likelihood phylogeny (for this genus, GCA_011526105.1 as the outgroup) showed that S4-bin21 was most closely related to genomes derived from stromatolite biomes. We compared the functional gene repertoire of S4-bin21 with that of the other four genomes in this genus available in the database. S4-bin21 encoded a unique GLUD1_2-type glutamate dehydrogenase (K00261); BLASTp searches against the NR database indicated that its closest homolog was HJL41477.1 from a marine sediment-derived Myxococcales bacterium (LLY-WYZ-16_1), whereas the other genomes in this genus used an NADP^+^-dependent glutamate dehydrogenase (K00262) (Fig. S25).

#### **11. Estimation of species divergence times under a molecular clock**

To estimate species divergence times, the MCMCTree program in PAML (v.4.10.7) was used (Dos Reis and Yang, 2019; Yang and Rannala, 2006). Divergence time constraints for specific prokaryotic lineages, inferred in previous studies and retrieved from TimeTree (<http://www.timetree.org>) (Kumar et al., 2017), were used to calibrate the maximum-likelihood trees. The resulting trees were visualized with tvBOT (Xie et al., 2023).

First, a maximum-likelihood phylogeny was reconstructed for the newly recovered *Marinobacter guineae* genome in this study (Fig. 5B), together with *Marinobacter guineae* M3B, two *Marinobacter persicus* strains, and *Halomonas ventosae* CECT 5797, based on 43 commonly conserved proteins identified by CheckM. The resulting alignment contained 6,646 amino-acid positions. Divergence times of 1,064 Ma for the split between the *Marinobacter* lineage and *Halomonas ventosae*, and 150 Ma for the divergence within the *Marinobacter* lineage (Marin et al., 2017), were used as calibration points (soft calibration). Two independent MCMC runs were performed, and convergence was evaluated from between-run agreement, Gelman–Rubin potential scale-reduction factors and effective sample sizes.

The estimated divergence time between the two *Marinobacter guineae* genomes, which originated from habitats that have remained geographically isolated for a long time, was 10.47 Ma (95% HPD: 2.18–20.68) (Fig. S26A), close to the estimated divergence time of approximately 11.04 Ma for the *Oceanicaulis* genus (Marin et al., 2017).

Second, maximum-likelihood phylogenies were similarly reconstructed for several representative MAG lineages from the community and their reference genomes, including the putative novel phototrophic *Oceanicaulis* MAG described in this study (Fig. 5A), the putative high-salinity clade within *Candidatus* Woesearchaeota UBA583 (Fig. 5C), two genomes classified as genus “PKUG01” in GTDB (of which the only database genome originated from estuary sediment), and a newly recovered *Pseudooceanicola* genome that was most closely related to *Pseudooceanicola nanhaiensis* (ANI = 86.9%). Divergence times of 1194 Ma between *Pseudooceanicola nanhaiensis* and *Oceanicaulis alexandrii* (Wang and Luo, 2021), and 2,540.0–2,805.8 Ma between the Pseudomonadota lineage containing these taxa and the Thermodesulfobacteriota lineage represented by *Desulfuromusa kysingii* (Marin et al., 2017; Sheridan et al., 2003), were used as calibration points. The estimated divergence times between the newly recovered genomes and their closest reference genomes, as well as among themselves, were all greater than 129 Ma (Fig. S26B), indicating independent evolutionary histories that were much longer than the divergence observed between *Marinobacter guineae* genomes and among *Oceanicaulis* lineages.

#### **12. Marine affinity of eukaryotes**

Further BLASTn searches against the NCBI core nucleotide database indicated that the closest sequence to OTU4 was KX160006, which was derived from starfish tissue (FioRito et al., 2016). OTU9 was most closely related to sequences MW843499 and MW843500 from microbial mats in an aquatic system in the Atacama Desert; these strains were previously proposed to have shared a marine ancestor and to have evolved into distinct species under environmental stress in the isolated inland saline lake, including high UV radiation (Schiwitza et al., 2021). Notably, the same environment has also yielded MAGs corresponding to four uncultured species of *Dactylococcopsis* deposited in the Genome database, which are closely related to S4-bin13 (Fig. S23: GCA_964463655.1, GCA_964489465.1, GCA_964493825.1, GCA_964346685.1), as well as the closest homolog of AoxB annotated in the Trueperaceae MAG from this study (MDZ7708468.1). Taken together, these OTUs provide representative examples that some eukaryotic members of the community also exhibit strong marine affinity.

#### **13. Arsenic metabolizing genes in MAGs**

Genes encoding three types of arsenate reductases were abundant in the MAG set recovered in this study. Among the 46 MAGs, 32 harbored at least one arsenate reductase, and 11 of them also possessed the capacity to methylate arsenite, thereby producing non-toxic organoarsenical compounds. Four distinct distribution patterns of these genes were observed across the MAG set, and the corresponding MAGs are indicated in Fig. S27A. Figure S27B further summarizes the species distribution and relative abundances of arsenic-metabolizing genes. Among arsenate reductases, the thioredoxin-dependent arsenate reductase (K03741) was the most prevalent (TPM levels are also higher than any genes related to nitrate/nitrite reduction), whereas the respiratory arsenate reductase (K28466) was most commonly encoded by Thermodesulfobacteriota genomes. Although *arsP* has not been widely functionally characterized in cultured organisms, its high prevalence in the present community is notable and exceeds that of *arsM*, in contrast to previous observations from marine sediments (Li et al., 2024). This pattern may indicate that methylarsenite tends to be exported after formation and that this capability is required by most community members.

### **Supplementary figures**

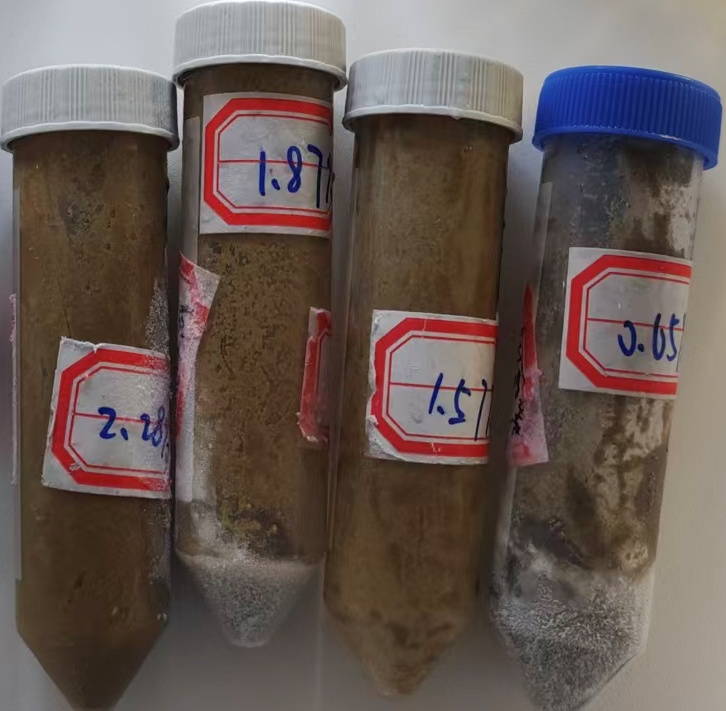

**S1**

**S2**

**S3**

**S4**

#### Figure S1. A photo of sediments sampled from the four sites.

The dark yellow, gravel-like sediments showed no significant reaction with sodium sulfide solution under alkaline conditions (pH: 11–12). Following mixing and agitation with 1 M sulfuric acid, the sediments were gradually corroded; however, the resulting pale-yellow supernatant exhibited no detectable reaction with 0.01 M potassium thiocyanate (KSCN). In addition, the absorbance of the supernatant decreased markedly across the 250–350 nm wavelength range (from 1.371 to 0.386). These observations suggest that the coloration of the sediments is more likely attributable to humic substances rather than iron-bearing minerals.

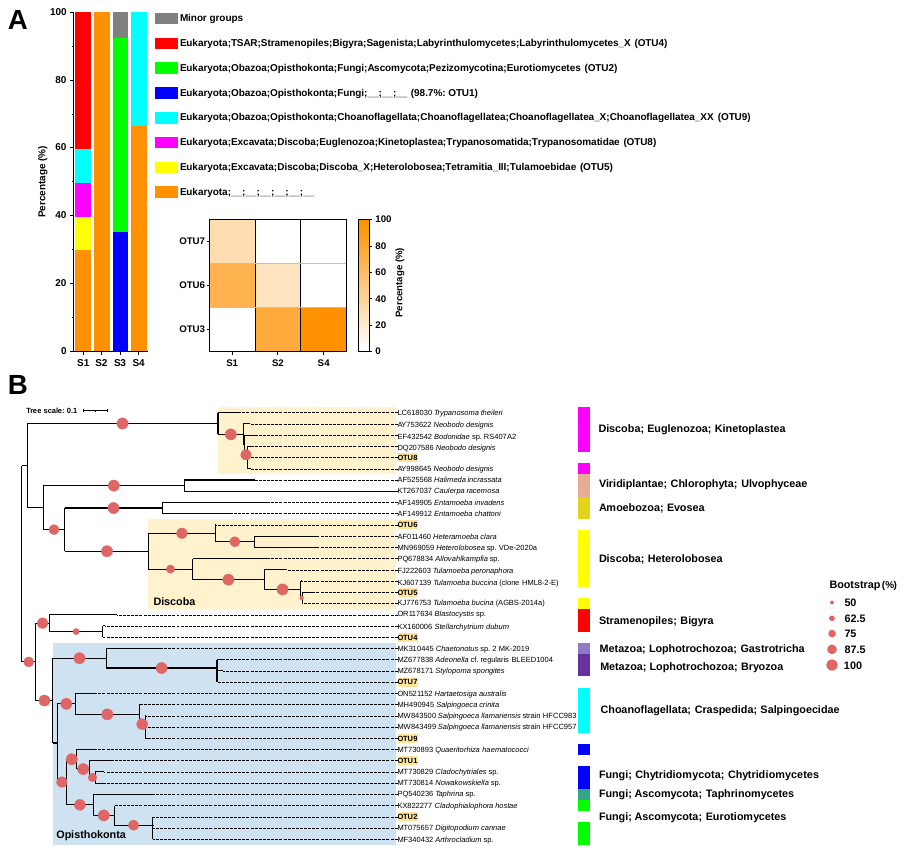

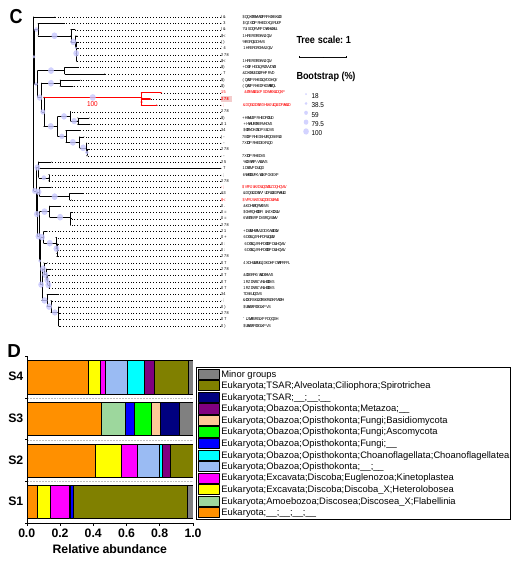

#### Figure S2. Eukaryotic community composition in sediments.

Based on 18S rRNA gene V9-region sequences: (A) Relative abundances of eukaryotic taxa with corresponding OTU identifiers; OTU1 assigned only to unclassified “Fungi” accounts for 98.7% of sequences. (B) Maximum-likelihood phylogeny constructed from the eight dominant OTUs together with reference 18S sequences. (C): OTU3, which is 75 bp in length, exhibited a sequence similarity of 97.06% to several Apicomplexa (parasitic protists) 18S rRNA gene sequences (e.g., NCBI accession numbers JX131298 and AY327258); however, it clustered with bacterial reference sequences affiliated with *Candidatus* Peregrinibacteria bacterium GW2011_GWA2_33_10 (KX123604) and *Clostridium pasteurianum* DSM 525 (NR_104822) in the phylogenetic tree. Given this ambiguous phylogenetic placement, OTU3 was not included in Figure B. (D): Eukaryotic community composition in sediments based on all 18S rRNA gene sequences (Only dominant taxa with a relative abundance of > 5% in at least one sample are shown. “Minor groups” denotes the combined category of OTUs other than dominant taxa and eukaryotic OTUs annotated as “Eukaryota;__;__;__;__” in the PR2 database).

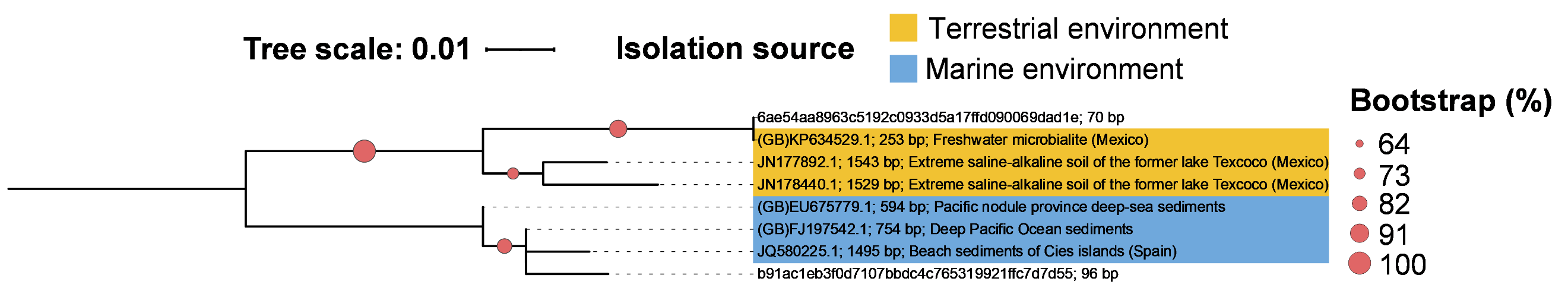

#### Figure S3. Maximum-likelihood phylogeny of representative V4-region OTUs assigned to the phyla “Nitrospirota” and non-redundant reference sequences from the SILVA 138 database and NCBI core nucleotide database (GeneBank accession shown).

The environmental origins of reference sequences (Terrestrial or marine environment) are indicated by distinct colors. In the phylogenetic tree reconstructed from 1,204 Nitrospirota reference sequences, sequence “6ae54aa8963c5192c0933d5a17ffd090069dad1e” clustered with JN177892.1 and JN178440.1. In BLASTn searches, KP634529.1 (not included in the SILVA database) was the closest sequence to this OTU (fast minimum-evolution / neighbor-joining analysis). Sequence “b91ac1eb3f0d7107bbdc4c765319921ffc7d7d55” clustered with JQ580225.1; in BLASTn searches, FJ197542.1 (not included in SILVA “Ref” dataset) showed the highest identity to this OTU (100%), whereas additional sequences with 98.95% identity, such as EU675779.1, were mostly derived from deep-sea environments across the four oceans. These eight sequences were realigned, trimmed, and used to reconstruct the maximum-likelihood tree shown here.

#### Figure S4. Maximum-likelihood phylogeny of representative V4-region OTUs assigned to “Acidobacteriota Subgroup 26” and non-redundant reference sequences from the SILVA 138 database and NCBI core nucleotide database.

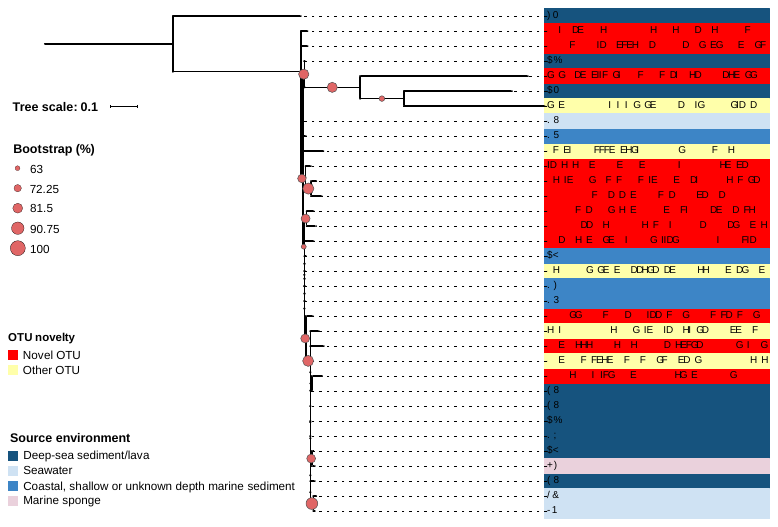

OTU novelty and the environmental origins of reference sequences are indicated by distinct colors (“Other OTU” means non novel OTU; legend indicates color–environment mapping).

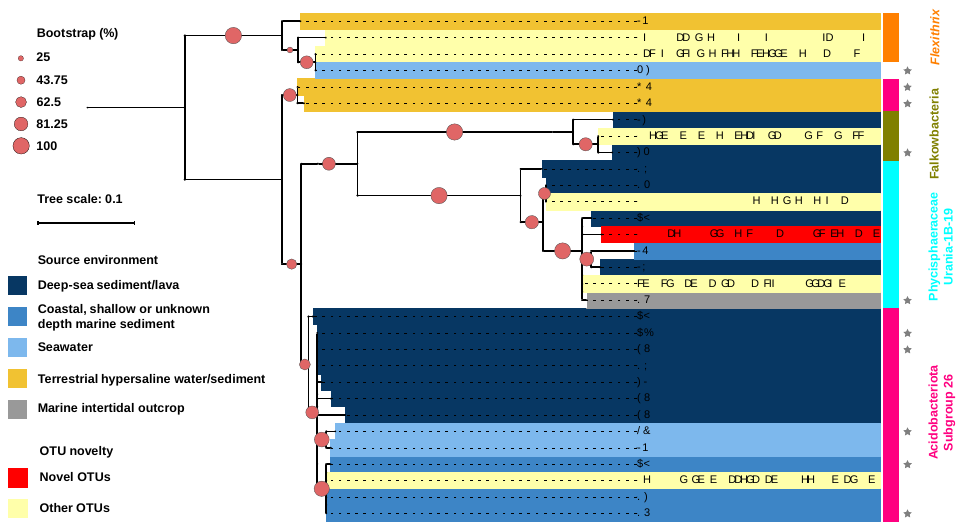

#### Figure S5. Maximum-likelihood phylogeny of representative high-abundance OTUs (relative abundance > 1% in at least one sample) and their closest reference sequences.

OTU novelty and the environmental origins of reference sequences are indicated by distinct colors (legend indicates color–environment mapping). The gray stars indicate sequences that are not included in the SILVA 138 reference set.

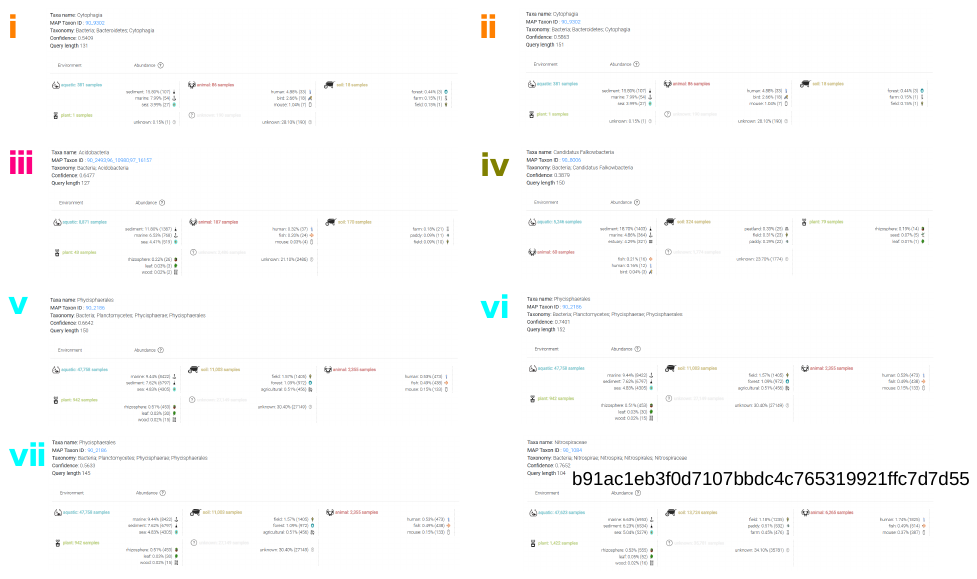

#### Figure S6. Environmental sample-occurrence profiles of MicrobeAtlas reference OTUs matched to representative bacterial OTUs.

Eight representative OTU sequences (i-ⅶ corresponds to the identification of OTU in Figure 2H) were submitted to the MicrobeAtlas MAPseq search tool, and the number and proportion of MicrobeAtlas samples containing the matched OTU across annotated habitat and sub-habitat categories were displayed. Percentages represent the fraction of all samples positive for the matched OTU that belonged to each environmental category; they do not represent mean relative abundance, habitat-normalized prevalence or sequence identity. The profiles therefore provide database-wide ecological context for the matched reference OTUs, used as supporting evidence for environmental affinity of the OTUs.

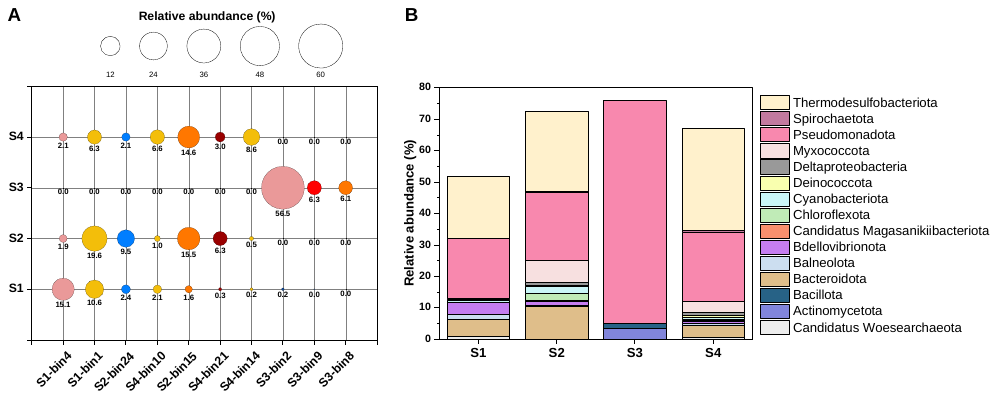

#### Figure S7. Relative abundances of dominant MAGs and MAG-derived phylum composition across samples.

(A) Relative abundances of the 10 dominant MAGs in each sample. (B) Phylum-level relative abundances inferred from MAGs (Deltaproteobacteria class with unresolved phylum-level affiliation).

#### Figure S8. Maximum-likelihood phylogeny of *dfx* protein sequence encoded by S4-bin11 and reference sequences.

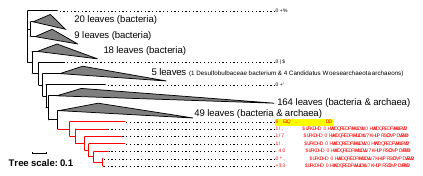

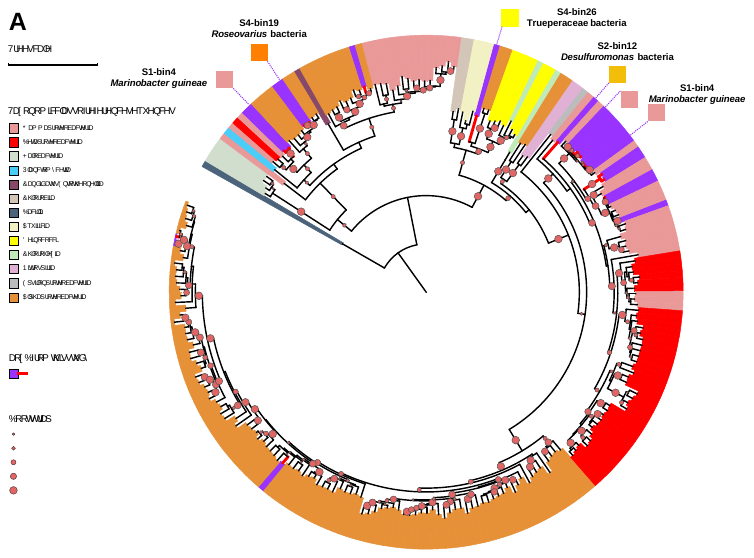

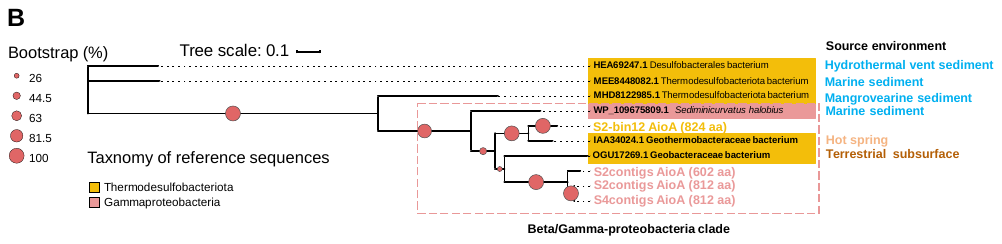

#### Figure S9. Phylogenetic analysis of *aoxB*.

(A) Maximum-likelihood phylogeny of *aoxB* protein sequences from the contigs and 214 reference sequences. (B) Maximum-likelihood phylogeny of Thermodesulfobacteriota AioA and representative AioA from Beta/Gamma-proteobacterial clade.

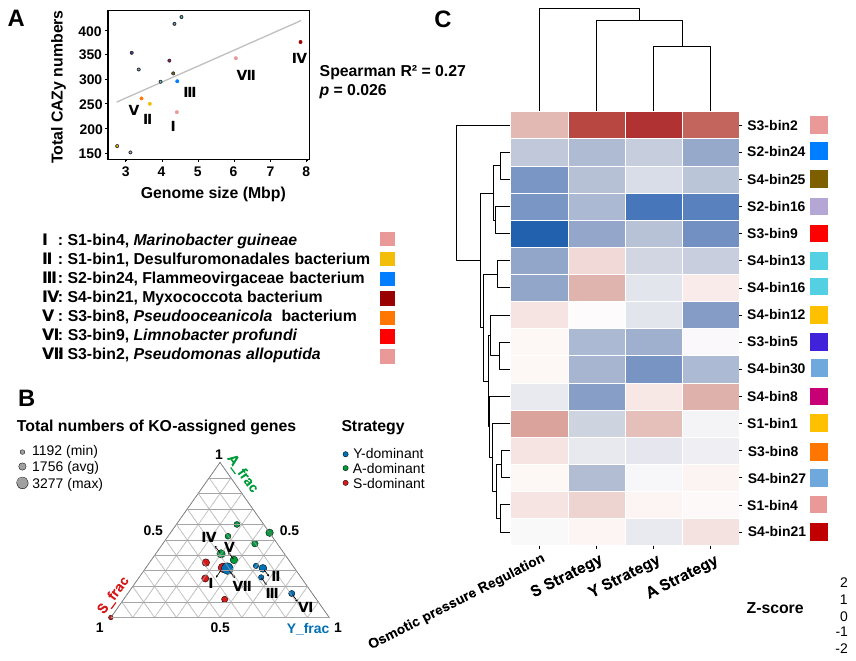

#### Figure S10. Predicted species life-history strategies.

(A) Correlation between total numbers of annotated CAZys and genome size across the 15 MAGs. (B) Inferred life-history strategy preferences of dominant MAGs based on the abundance of strategy-defining gene sets. (C) Hierarchical clustering heatmap of 16 genomes across four functional dimensions: scores for the three life-history strategies (Y/A/S) and a separate score for osmotic-pressure-regulation genes (gene set shown in Fig. 5B). Color scale denotes normalized scores (Z score).

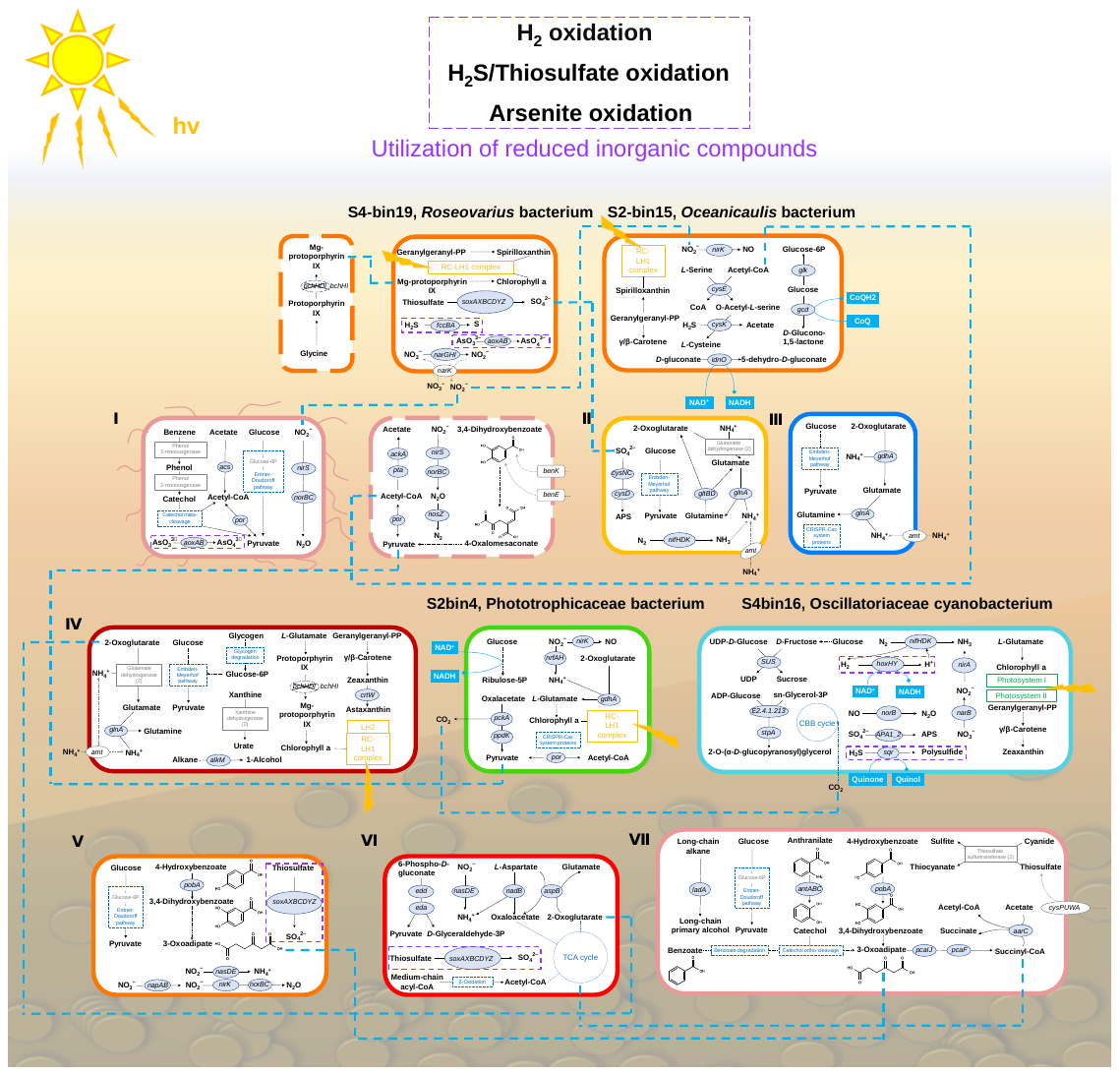

#### Figure S11. Characteristic metabolic patterns of representative MAGs.

MAGs I–VII correspond to the MAG numbers used in Fig. S10. Cyanobacteriota are the phototrophic autotrophs in the community; although their relative abundance was always < 3%, the metabolic map of S4-bin16 (Oscillatoriaceae cyanobacterium) is also shown. In addition, other phototroph-associated heterotrophs are shown, including the dominant MAG S4-bin21 (IV, Myxococcota bacterium) and S2-bin15 (*Oceanicaulis* bacterium), as well as S4-bin19 (*Roseovarius* bacterium) and S2-bin4 (Phototrophicaceae bacterium), which showed higher completeness and lower contamination than the other two MAGs assigned by GTDB-Tk to the “CAIRDJ01” genus. In the metabolic maps, the utilization of reduced inorganic compounds (including hydrogen oxidation, H_2_S/thiosulfate oxidation, and arsenite oxidation) is highlighted in purple boxes. Potential interspecies material exchange is indicated by cyan dashed lines. Note that the positions of genes or gene clusters do not necessarily reflect their precise localization within cellular substructures. For further details, see Figs. S12–S21.

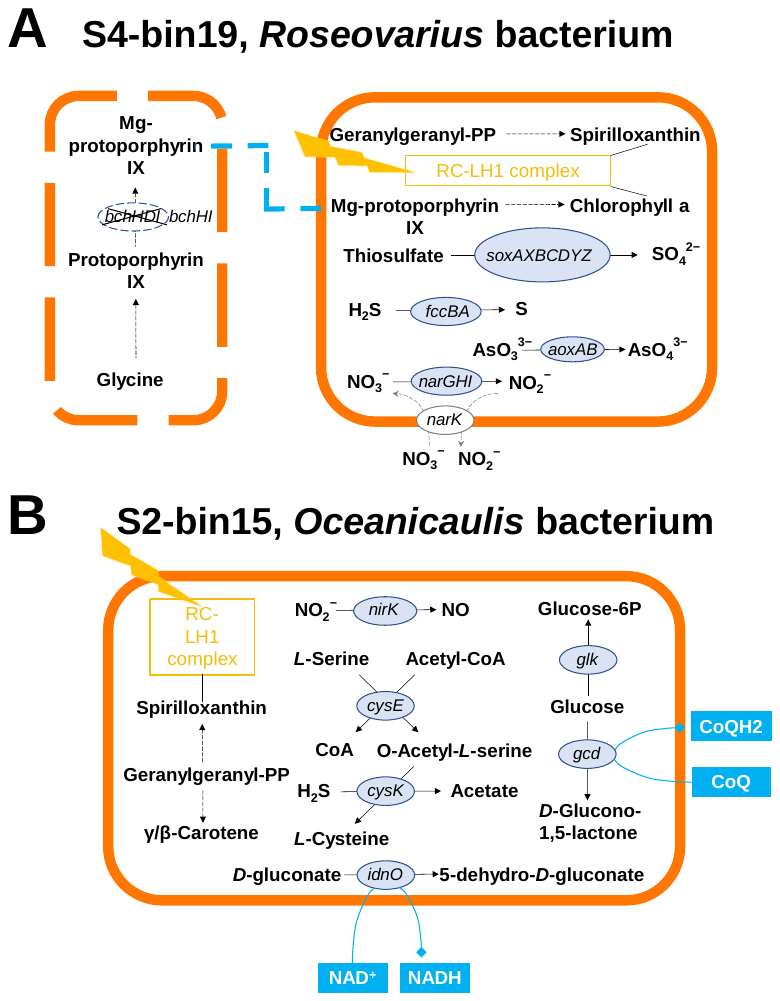

#### Figure S12. Simplified metabolic schematics of putatively phototrophic alphaproteobacterial MAGs harboring the RC-LH1 complex. Black dashed lines indicate multi-step reactions mediated by multiple enzymes, and gray curved lines indicate transporter-mediated movement of molecules or ions (the same convention applies to Figs. S13–S21).

(A) S4-bin19, *Roseovarius* bacterium. Contig-level annotations also identified genes implicated in the glycine-dependent synthesis of Mg-protoporphyrin IX in *Roseovarius*; although the magnesium chelatase subunit *bchD* is absent, this conversion is shown in the dashed box.

(B) S2-bin15, *Oceanicaulis* bacterium. The conversion of protochlorophyllide to bacteriochlorophyll a is not shown here because it is presented in Fig. 5A.

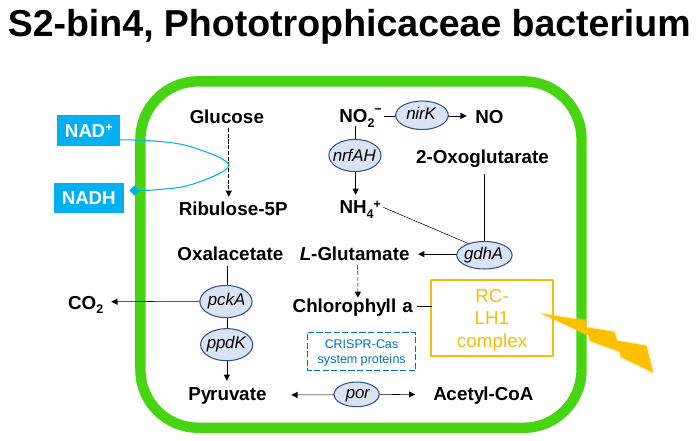

#### Figure S13. Simplified metabolic schematic of MAG S2-bin4 (Phototrophicaceae bacterium).

Compared with other CAIRDJ01 genomes, S2-bin4 not only harbors the RC-LH1 complex but also shows enhanced carbon utilization capacity, including oxidation of glucose (M00006: K00036 + K01057 + K00033) and oxaloacetate (K01610), nitrite reduction pathways (assimilatory reduction to ammonium and dissimilatory reduction to nitric oxide), L-glutamate-dependent chlorophyll a biosynthesis, and CRISPR-Cas systems.

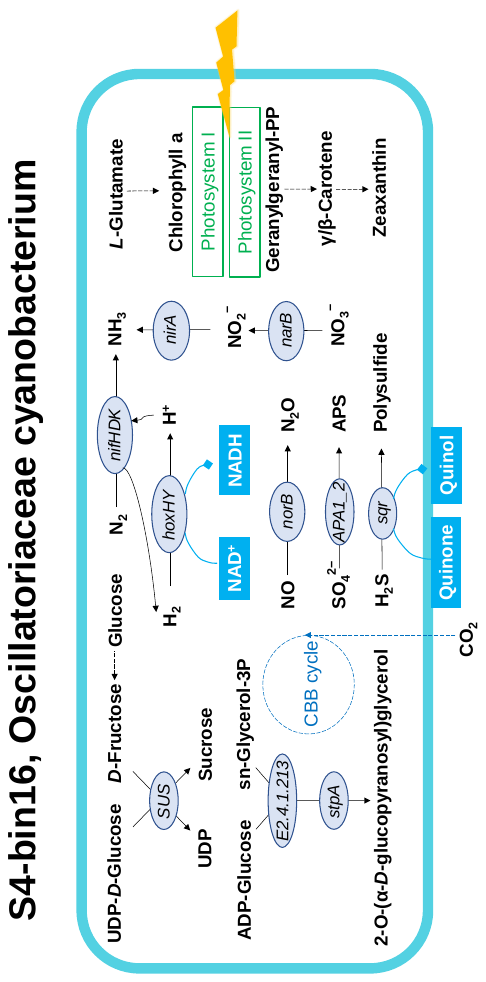

#### Figure S14. Simplified metabolic schematic of MAG S4-bin16 (Oscillatoriaceae cyanobacterium).

It may adapt to high-salinity conditions by accumulating osmoprotective compounds, whereas the capacity for sucrose or glucosylglycerol biosynthesis is rare in the community. It may also participate in a range of biochemical redox reactions.

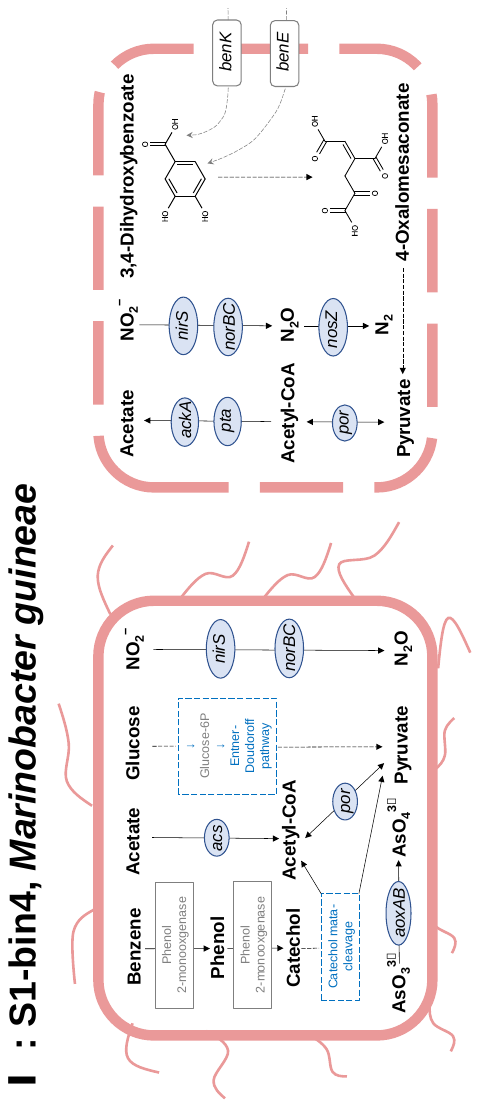

#### Figure S15. Simplified metabolic schematic of MAG S1-bin4 (*Marinobacter guineae*).

Some Marinobacter-affiliated functional genes annotated on contigs were not assigned to the *M. guineae* MAG during assembly and binning, but instead clustered into another MAG (completeness 84.45%, contamination 2.69%). Although both topology-based phylogenetic placement and ANI values indicate that this MAG also belongs to the genus *Marinobacter*, it failed the GUNC quality assessment and was therefore excluded from the analyses described above (S1-bin5, *Marinobacter* MAG abandoned). Nevertheless, the functional genes recovered from this MAG point to a higher functional and taxonomic diversity of *Marinobacter* within the community. These genes suggest potential involvement in acetate production via the Pta–AckA pathway, the independent degradation of 3,4-dihydroxybenzoate into pyruvate and oxaloacetate, and the reduction of nitrous oxide to dinitrogen (the dashed box on the right side).

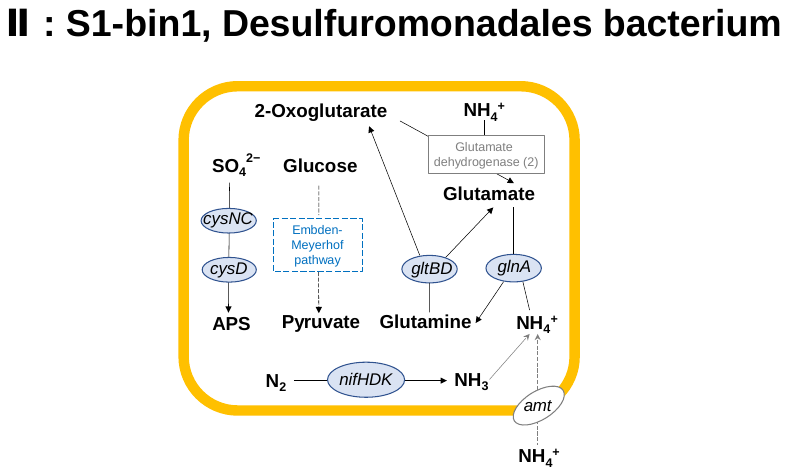

#### Figure S16. Simplified metabolic schematic of MAG S1-bin1 (Desulfuromonadales bacterium).

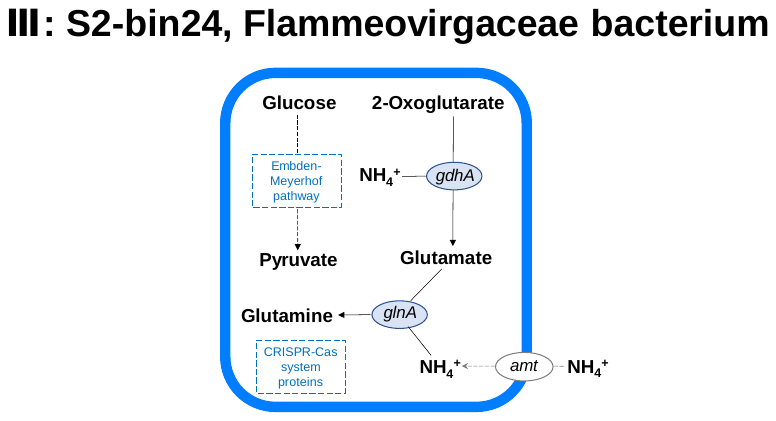

#### Figure S17. Simplified metabolic schematic of MAG S2-bin24 (Flammeovirgaceae bacterium).

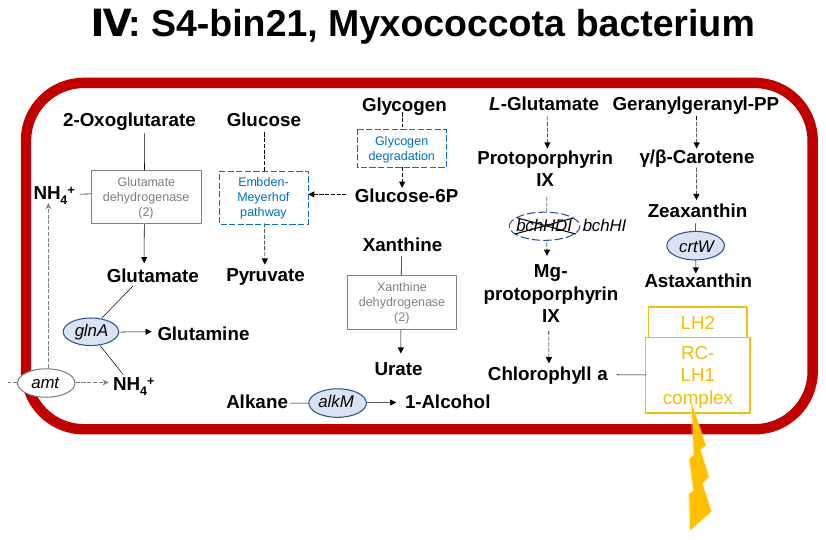

#### Figure S18. Simplified metabolic schematic of MAG S4-bin21 (Myxococcota bacterium).

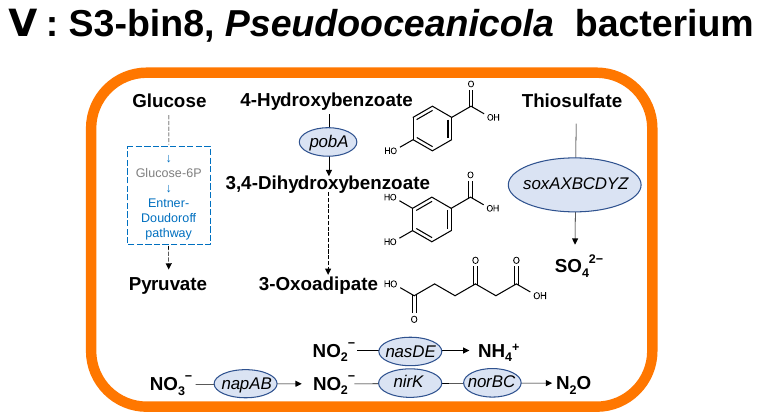

#### Figure S19. Simplified metabolic schematic of MAG S3-bin8 (*Pseudooceanicola* bacterium).

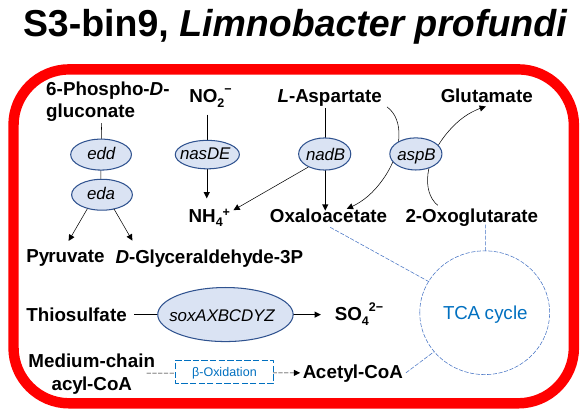

#### Figure S20. Simplified metabolic schematic of MAG S3-bin9 (*Limnobacter profundi*).

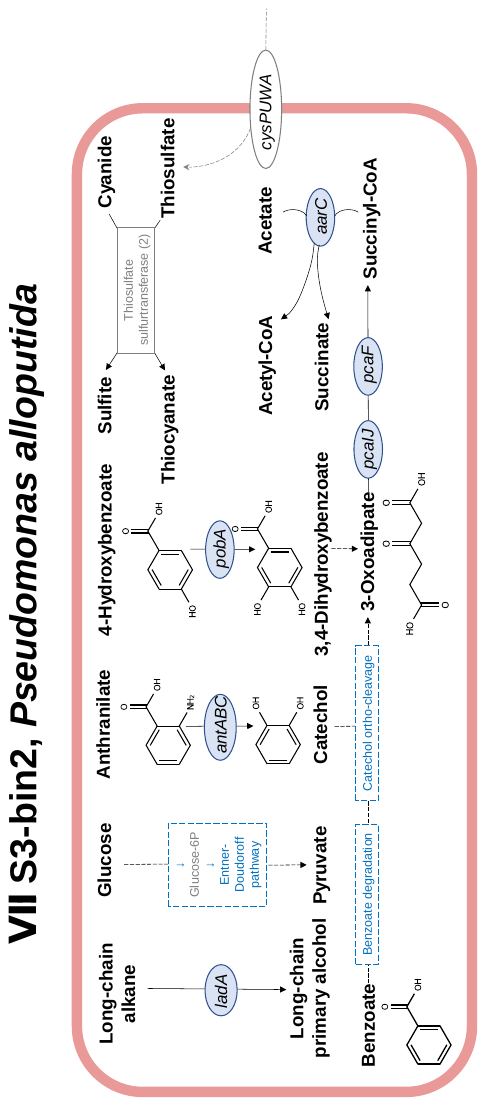

#### Figure S21. Simplified metabolic schematic of MAG S3-bin2 (*Pseudomonas alloputida*).

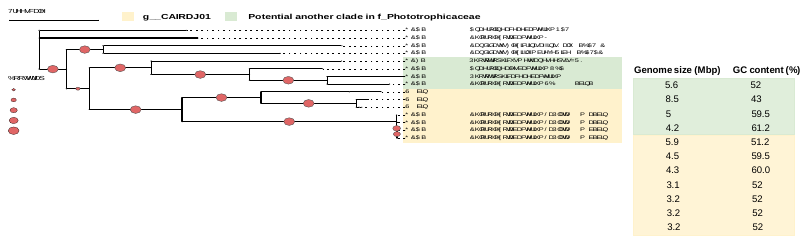

#### Figure S22. Maximum-likelihood phylogeny of representative Phototrophicaceae MAGs.

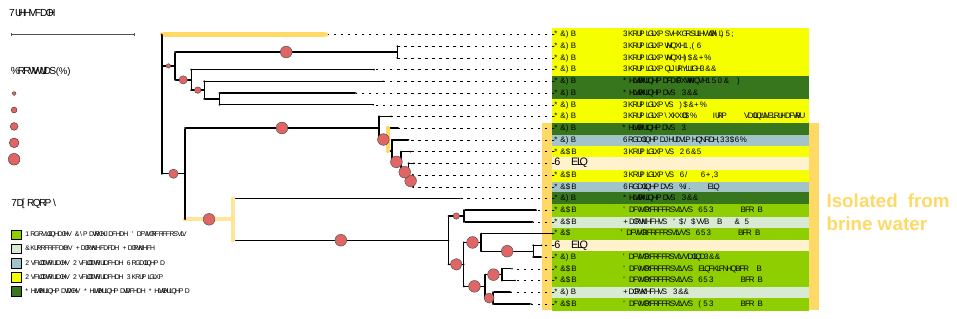

#### Figure S23. Maximum-likelihood phylogeny of the 2 reconstructed Cyanophyceae MAGs from this study and their reference genomes (based on the concatenated alignment of 43 commonly conserved proteins predicted by CheckM).

#### Figure S24. Phylogenetic analysis of the novel Flammeovirgaceae MAG S2-bin24.

(A) Maximum-likelihood phylogeny using 14 Flammeovirgaceae “reference genomes” from the NCBI Genome database and *Cytophaga aurantiaca* DSM 3654 (GCF_000379725.1) as the outgroup. (B) Maximum-likelihood phylogeny using 75 Flammeovirgaceae genomes from GTDB.

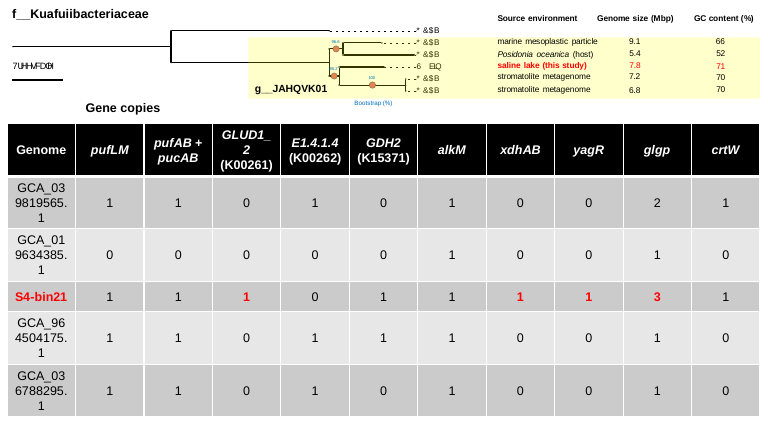

#### Figure S25. Comparative genomic analysis of “*Candidatus* Kuafubacteriaceae JAHQVK01” MAGs.

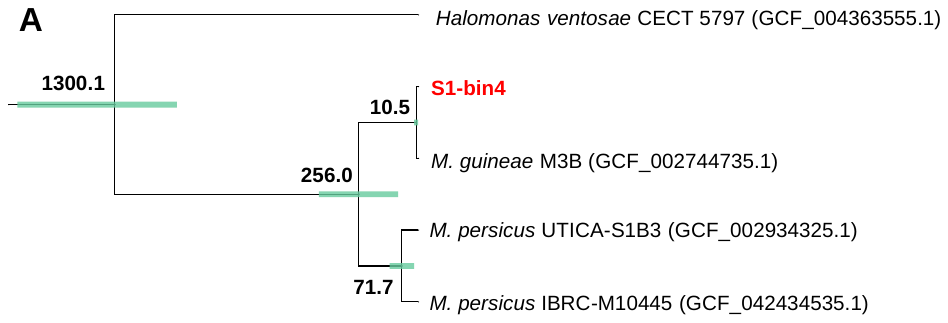

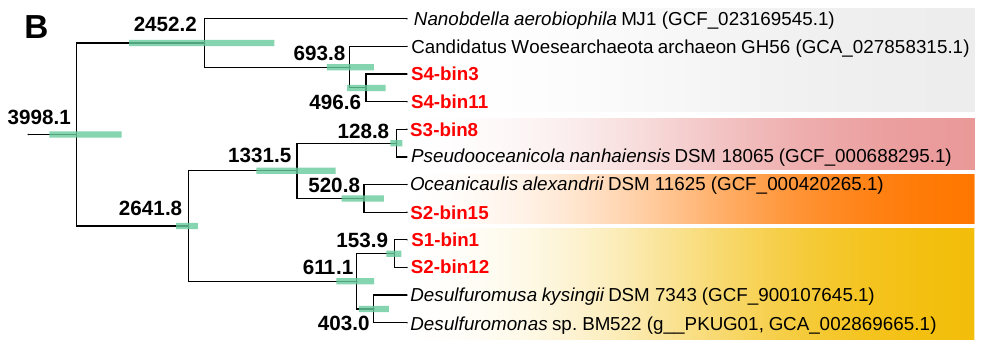

#### Figure S26. Molecular-clock estimates of divergence times for phylogenetic clusters containing newly recovered MAGs.

(A) Time-calibrated phylogeny for S1-bin4 (*Marinobacter guineae*). (B) Time-calibrated phylogeny for other representative novel MAGs. Bold labels at internal nodes indicate posterior mean divergence times (Ma), and horizontal bars indicate 95% highest posterior density (HPD) intervals.

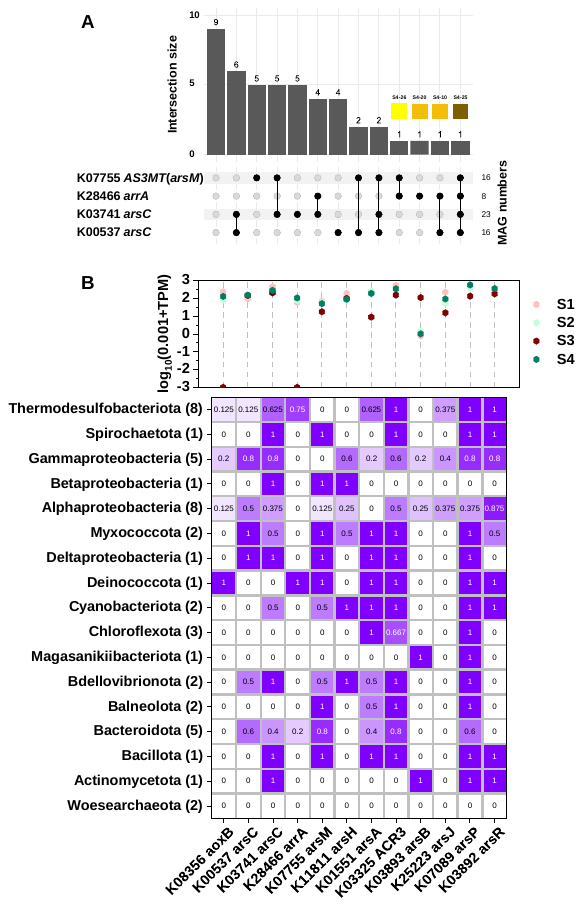

#### Figure S27. Distribution of arsenic metabolizing genes in MAGs.

(A) Co-occurrence patterns of arsenate reductase genes (K00537, K03741, and K28466) and arsenite methyltransferase (*arsM*, K07755) across the 46 MAGs.

(B) Relative abundances (TPM) and prevalence of arsenic-metabolizing genes across MAG lineages. The gene *arsH* encodes a methylarsenite-specific oxidase; *arsA* forms an arsenite efflux pump that exports arsenite from the cytoplasm; *ACR3* and *arsB* also mediate arsenite transport; *arsJ* and *arsP* function as organoarsenical efflux permeases; and *arsR* encodes an arsenite-responsive repressor that regulates *ars* operon transcription.

### **References**

Chen, S., Zhou, Y., Chen, Y. and Gu, J. 2018. fastp: an ultra-fast all-in-one FASTQ preprocessor. Bioinformatics 34(17), 884–890.

Cheng, S., Xue, W., Gong, X., Hu, F., Yang, Y. and Liu, M. 2024. Reconciling plant and microbial ecological strategies to elucidate cover crop effects on soil carbon and nitrogen cycling. Journal of Ecology 112(12), 2901–2916.

Dos Reis, M. and Yang, Z. (2019) EVOLUTIONARY GENOMICS, 2 EDITION: Statistical and Computational Methods. Anisimova, M. (ed), pp. 309–330.

Ehira, S., Kimura, S., Miyazaki, S. and Ohmori, M. 2014. Sucrose synthesis in the nitrogen-fixing cyanobacterium Anabaena sp. strain PCC 7120 is controlled by the two-component response regulator OrrA. Applied and Environmental Microbiology 80(18), 5672–5679.

FioRito, R., Leander, C. and Leander, B. 2016. Characterization of three novel species of Labyrinthulomycota isolated from ochre sea stars (Pisaster ochraceus). Marine Biology 163(8), 170.

Hagemann, M. 2011. Molecular biology of cyanobacterial salt acclimation. FEMS Microbiology Reviews 35(1), 87–123.

Hakobyan, L., Monforte-Gomez, B., Moliner-Martinez, Y., Molins-Legua, C. and Campins-Falco, P. 2022. Improving sustainability of the Griess reaction by reagent stabilization on PDMS membranes and ZnNPs as reductor of nitrates: application to different water samples. Polymers 14(3), 464.

He, Y.H., Baltar, F. and Wang, Y. 2025. Seasonal variability in community structure and metabolism of active deep-sea microorganisms. ISME J 19(1), wraf214.

Kumar, S., Stecher, G., Suleski, M. and Hedges, S. 2017. TimeTree: a resource for timelines, timetrees, and divergence times. Molecular Biology and Evolution 34(7), 1812–1819.

Li, L., Huang, D., Hu, Y., Rudling, N., Canniffe, D., Wang, F. and Wang, Y. 2023. Globally distributed Myxococcota with photosynthesis gene clusters illuminate the origin and evolution of a potentially chimeric lifestyle. Nature Communications 14(1), 6450.

Li, Z., He, Y., Zhang, H., Qian, H. and Wang, Y. 2024. Biotransformations of arsenic in marine sediments across marginal slope to hadal zone. Journal of Hazardous Materials 480, 135955.

Luo, Q., Duan, Y. and Lu, X. 2022. Biological sources, metabolism, and production of glucosylglycerols, a group of natural glucosides of biotechnological interest. Biotechnology Advances 59, 107964.

Malik, A., Martiny, J., Brodie, E., Martiny, A., Treseder, K. and Allison, S. 2020. Defining trait-based microbial strategies with consequences for soil carbon cycling under climate change. ISME J 14(1), 1–9.

Marin, J., Battistuzzi, F., Brown, A. and Hedges, S. 2017. The timetree of prokaryotes: new insights into their evolution and speciation. Molecular Biology and Evolution 34(2), 437–446.

Rashid, F., Crémazy, F., Hofmann, A., Forrest, D., Grainger, D., Heermann, D. and Dame, R. 2023. The environmentally-regulated interplay between local three-dimensional chromatin organisation and transcription of proVWX in E. coli. Nature Communications 14(1), 7478.

Santos-Merino, M., Yun, L. and Ducat, D. 2023. Cyanobacteria as cell factories for the photosynthetic production of sucrose. Frontiers in Microbiology 14, 1126032.

Schiwitza, S., Gutsche, L., Freches, E., Arndt, H. and Nitsche, F. 2021. Extended divergence estimates and species descriptions of new craspedid choanoflagellates from the Atacama Desert, Northern Chile. European Journal of Protistology 79, 125798.

Sheridan, P., Freeman, K. and Brenchley, J. 2003. Estimated minimal divergence times of the major bacterial and archaeal phyla. Geomicrobiology Journal 20(1), 1–14.

Sorouri, B., Scales, N., Gaut, B. and Allison, S. 2024. Sphingomonas clade and functional distribution with simulated climate change. Microbiology Spectrum 12(5), e00236–00224.

Veloso, M., Waldisperg, A., Arros, P., Berríos-Pastén, C., Acosta, J., Colque, H., Varas, M., Allende, M., Orellana, L. and Marcoleta, A. 2023. Diversity, taxonomic novelty, and encoded functions of Salar de Ascotán microbiota, as revealed by metagenome-assembled genomes. Microorganisms 11(11), 2819.

Wang, S. and Luo, H. 2021. Dating Alphaproteobacteria evolution with eukaryotic fossils. Nature Communications 12(1), 3324.

Wood, J., Tang, C. and Franks, A. 2018. Competitive traits are more important than stress-tolerance traits in a cadmium-contaminated rhizosphere: a role for trait theory in microbial ecology. Frontiers in Microbiology 9, 121.

Xie, J., Chen, Y., Cai, G., Cai, R., Hu, Z. and Wang, H. 2023. Tree Visualization By One Table (tvBOT): a web application for visualizing, modifying and annotating phylogenetic trees. Nucleic Acids Research 51(W1), W587–W592.

Xu, H., Luo, X., Qian, J., Pang, X., Song, J., Qian, G., Chen, J. and Chen, S. 2012. FastUniq: a fast de novo duplicates removal tool for paired short reads. PloS One 7(12), e52249.

Yang, Z. and Rannala, B. 2006. Bayesian estimation of species divergence times under a molecular clock using multiple fossil calibrations with soft bounds. Molecular Biology and Evolution 23(1), 212–226.

Zheng, R., Wang, C. and Sun, C. 2024. Deep-sea in situ and laboratory multi-omics provide insights into the sulfur assimilation of a deep-sea Chloroflexota bacterium. MBio 15(4), e00004–00024.
